# Elucidating Human Milk Oligosaccharide biosynthetic genes through network-based multiomics integration

**DOI:** 10.1101/2020.09.02.278663

**Authors:** Benjamin P. Kellman, Anne Richelle, Jeong-Yeh Yang, Digantkumar Chapla, Austin W. T. Chiang, Julia Najera, Bokan Bao, Natalia Koga, Mahmoud A. Mohammad, Anders Bech Bruntse, Morey W. Haymond, Kelley W. Moremen, Lars Bode, Nathan E. Lewis

## Abstract

Human Milk Oligosaccharides (HMOs) are abundant carbohydrates fundamental to infant health and development. Although these oligosaccharides were discovered more than half a century ago, their biosynthesis in the mammary gland remains largely uncharacterized. Here, we used a systems biology framework that integrated glycan and RNA expression data to construct an HMO biosynthetic network and predict glycosyltransferases involved. To accomplish this, we constructed models describing the most likely pathways for the synthesis of the oligosaccharides accounting for >95% of the HMO content in human milk. Through our models, we propose candidate genes for elongation, branching, fucosylation, and sialylation of HMOs. We further explored selected enzyme activities through kinetic assay and their co-regulation through transcription factor analysis. These results provide the molecular basis of HMO biosynthesis necessary to guide progress in HMO research and application with the ultimate goal of understanding and improving infant health and development.

**Significance statement:** With the HMO biosynthesis network resolved, we can begin to connect genotypes with milk types and thereby connect clinical infant, child and even adult outcomes to specific HMOs and HMO modifications. Knowledge of these pathways can simplify the work of synthetic reproduction of these HMOs providing a roadmap for improving infant, child, and overall human health with the specific application of a newly limitless source of nutraceuticals for infants and people of all ages.

## 1 Introduction

Human milk is the “gold standard” of nutrition during early life ^1–3^. Beyond lactose, lipids, and proteins, human milk contains 11-17% (dry weight) oligosaccharides (Human Milk Oligosaccharides, HMOs)^4,5^. HMOs are milk bioactives known to improve infant immediate and long-term health and development^2,6^. HMOs are metabolic substrates for specific beneficial bacteria (e.g., *Lactobacillus* spp. and *Bifidobacter* spp.), and shape the infant’s gut microbiome ^2,7^. HMOs also impact the infant’s immune system, protect the infant from intestinal and immunological disorders (e.g., necrotizing enterocolitis, HIV, etc.), and may aid in proper brain development and cognition ^2,6,8,9^. In addition, recent discoveries show that some HMOs can be beneficial to humans of all ages, e.g. the HMO 2’-fucosyllactose (2’FL) protecting against alcohol-induced liver disease^10^.

The biological functions of HMOs are determined by their structures ^6^. HMOs are unconjugated glycans consisting of 3–20 total monosaccharides draw from 3-5 unique monosaccharides: galactose (Gal, A), glucose (Glc, G), N-acetylglucosamine (GlcNAc, GN), fucose (Fuc, F) and the sialic acid N-acetyl-neuraminic acid (NeuAc, NN) (**Figure** 1A). All HMOs extend from a common lactose (Galβ1-4Glc) core. The core lactose can be extended at the nonreducing end, with a β-1,3-GlcNAc to form a trisaccharide. That intermediate trisaccharide is quickly extended on its non-reducing terminus with a β-1,3-linked galactose to form a type-I tetrasaccharide (LNT) or a β-1,4-linked galactose to form a type-II tetrasaccharide (LNnT). Additional branching of the trisaccharide or tetrasaccharide typically occurs at the lactose core by addition of a β-1,6-linked GlcNAc to the Gal residue. Lactose or the elongated oligosaccharides can be further fucosylated in an α-1,2-linkage to the terminal Gal residue, or α1,3/4-fucosylated on internal Glc or GlcNAc residues, and α-2,3-sialylated on the terminal Gal residue or α-2,6-sialylated on external Gal or internal GlcNAc residues^6,8^(**Figure** 1B).

**Figure 1.**
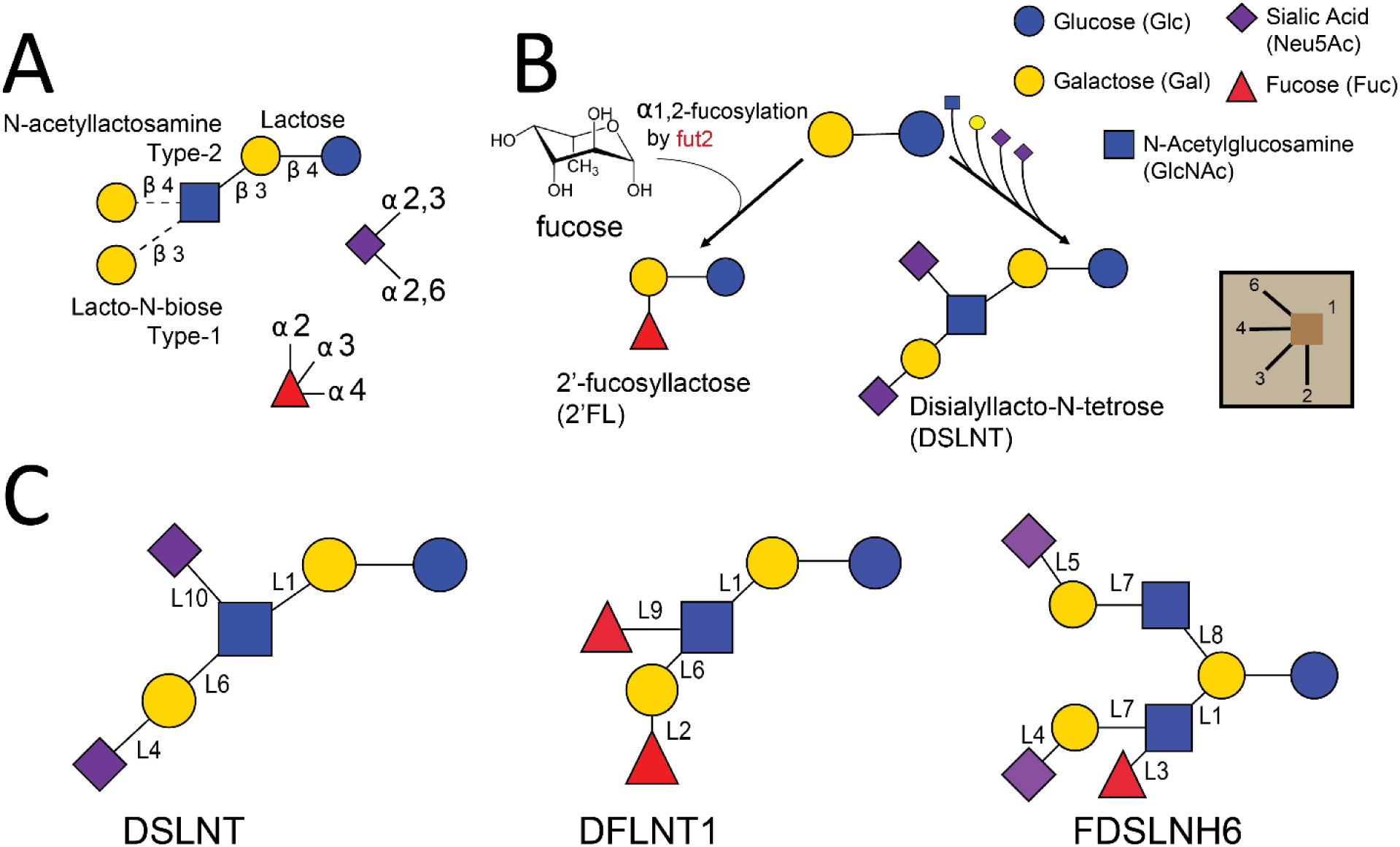
HMO blueprint and synthesis. **(A)** HMOs are built from a combination of the five monosaccharides D-glucose (Glc, blue circle), D-galactose (Gal, yellow circle), N-acetyl-glucosamine (GlcNac, blue square), L-fucose (Fuc, red triangle), and sialic acid (N-acetyl-neuraminic acid (NeuAc), purple diamond). Lactose (Gal-β-1,4-Glc) forms the reducing end and can be elongated with several Lacto-N-biose or N-acetyllactosamine repeat units (Gal-β-1,3/4-GlcNAc). Lactose or the polylactosamine backbone can be fucosylated with α-1,2-, α-1,3-, or α-1,4-linkages or sialylated in α-2,3- or α-2,6-linkages ^2^. **(B)** Small HMOs can be fucosylated to make 2’FL while larger HMOs can be synthesized by the extension of the core lactose with N-acetylactosamine (type-I) or lacto-N-biose (type-II) and subsequent decoration of the extended core with sialic acid to make more complex HMOs, such as DSLNT. **(C)** Three HMOs in this study: DSLNT, isomer 1 of DFLNT, isomer 6 of FDSLNH; isomer structures represent predictions from this study (see Methods, **Figure** S 12). Each monosaccharide-linking glycosidic bond is labeled (L1, L2,…L10) according to the linkage reactions listed in **Table** 1.

Despite decades of study, many details of HMO biosynthesis remain unclear. While the many possible monosaccharide addition events above are known, the order of the biosynthetic steps and many of the enzymes involved are not known (**Table** 1). For example, the lactose core is extended by alternating actions of β-1,3-N-acetylglucosaminyltransferases (b3GnT) and β-1,4-galactosaminyltransferases (b4GalT) while β-galactoside sialyltransferases (SGalT) and α-1,2-fucosyltransferases (including the FUT2 ‘secretor’ locus) are responsible for some sialylation and fucosylation of a terminal galactose, respectively ^11^. However, each enzymatic activity in HMO extension and branching can potentially be catalyzed by multiple isozymes in the respective gene family. Direct evidence of the specific isozymes performing each reaction *in vivo* is extremely limited.

**Table 1.**
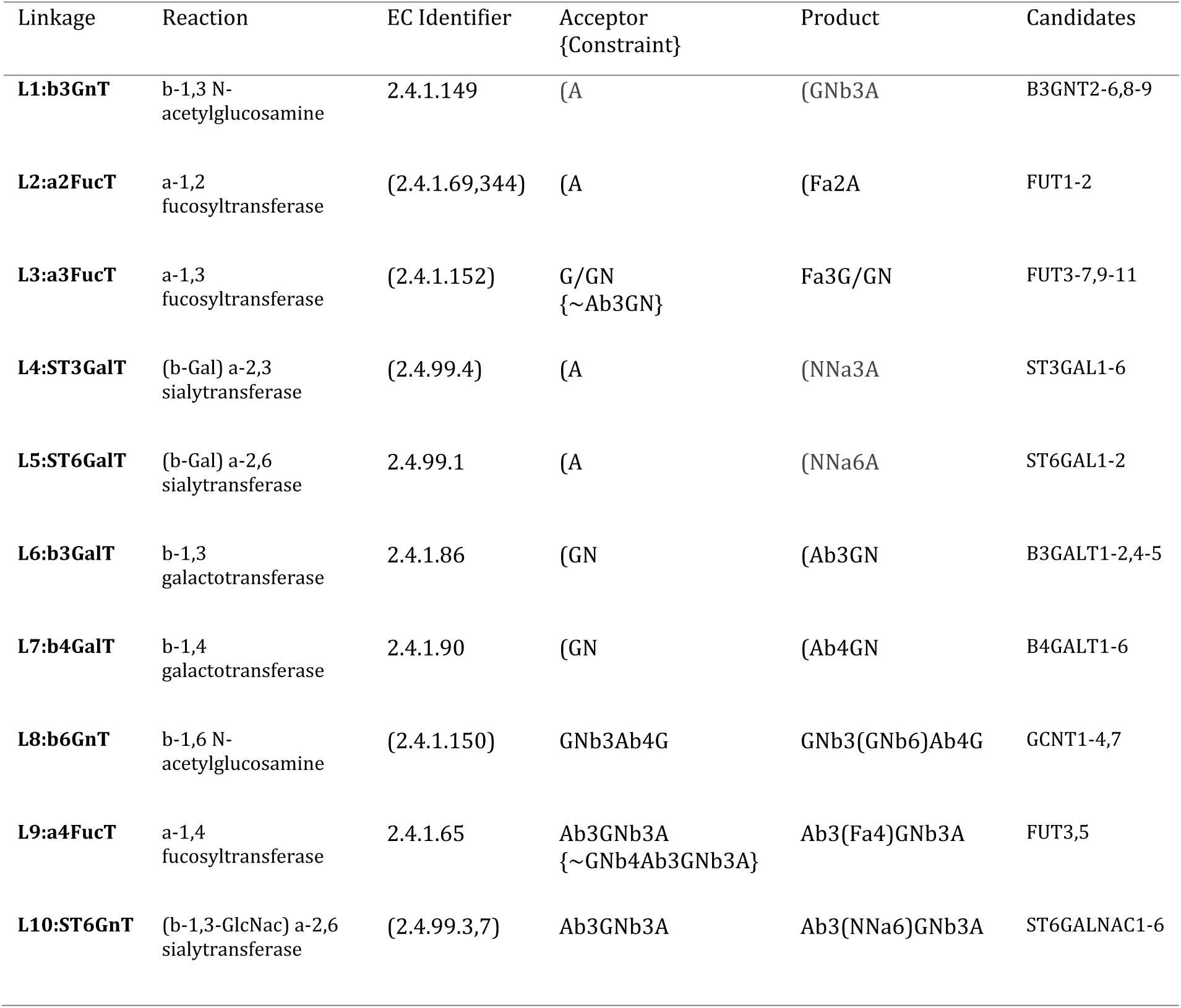
Glycosylation reactions examined. We studied here several candidate glycosyltransferases expressed in our samples to identify candidates for 10 elementary reactions (see Methods, **Table** S 1). Acceptor, product and constraint are represented in LiCoRR^12^: monosaccharides include Gal (A), Fuc (F), Glc (G), GlcNAc (GN), Neu5Ac (NN). Additionally, “)” and “(“indicate initiation and termination of a branch respectively, “[X/Y]” indicates either monosaccharide, and “∼” indicates a negation. An asterisk “*” indicates an imperfect match between the EC number and reaction. Background colors correspond to the monosaccharide added: GlcNAc (blue), Fuc (red), Neu5Ac (purple), and Gal (yellow).

Here we leverage the heterogeneity in HMO composition and gene expression across human subjects to refine our knowledge of the HMO biosynthetic network. Milk samples were collected from 11 lactating women across two independent cohorts between the 1st and 42nd day post-partum. (see methods). Gene expression profiling of mammary epithelial cells was obtained from mRNA present in the milk fat globule membrane interspace. Absolute and relative concentrations of the 16 most abundant HMOs was measured. Starting from a scaffold of all possible reactions^13–18^, we used constraint-based modeling^19,20^ to reduce the network to a set of relevant reactions and most plausible HMO structures when not known^21^ to form the basis for a mechanistic model^13,14,22^. This resulted in a ranked ensemble of candidate biosynthetic pathway topologies. We then ranked 44 million candidate biosynthesis networks to identify the most likely network topologies and candidate enzymes for each reaction by integrating sample-matched transcriptomic and glycoprofiling data from the 11 subjects. For this we simulated all reaction fluxes and tested the consistency between changes in flux and gene expression to determine the most probable gene isoforms responsible for each linkage type. We followed with direct observations through fluorescence activity assays to confirm our predictions. Finally, we performed transcription factor analysis to delineate regulators of the system. The resulting knowledge of the biosynthetic network can guide efforts to unravel the genetic basis of variations in HMO composition across subjects, populations, and disorders using systems biology modeling techniques.

## 2 Results

### 2.1 Abundances of HMOS and their known enzymes do not correlate

While α-1,2-fucosylation of glycans in humans can be accomplished by both FUT1 and FUT2, only FUT2 is expressed in mammary gland epithelial cells (**Table** S 1). FUT2, the “secretor” gene, is essential to ABH antigens^23–25^ as well as HMO ^2,26,27^ expression. We confirmed that non-functional FUT2 in “non-secretor” subjects guarantees the near-absence of α-1,2-fucosylated HMOs like 2’FL and LNFP1 (Fig2C). But, examining only subjects with functional FUT2 (Secretors), we found FUT2 expression levels and the concentration (nmol/ml) of HMOs containing α-1,2-fucosylation do not correlate in sample-matched microarray and glycomic measurements by HPLC (**Figure** 2). Generalized Estimating Equations (GEE) showed no significant positive association (2’FL Wald p = 0.056; LNFPI Wald p = 0.34). FUT1 could catalyze this reaction but its expression was not detected in these samples. We hypothesized that to successfully connect gene expression to HMO synthesis, one must account for all biosynthetic steps and not solely rely on direct correlations.

**Figure 2.**
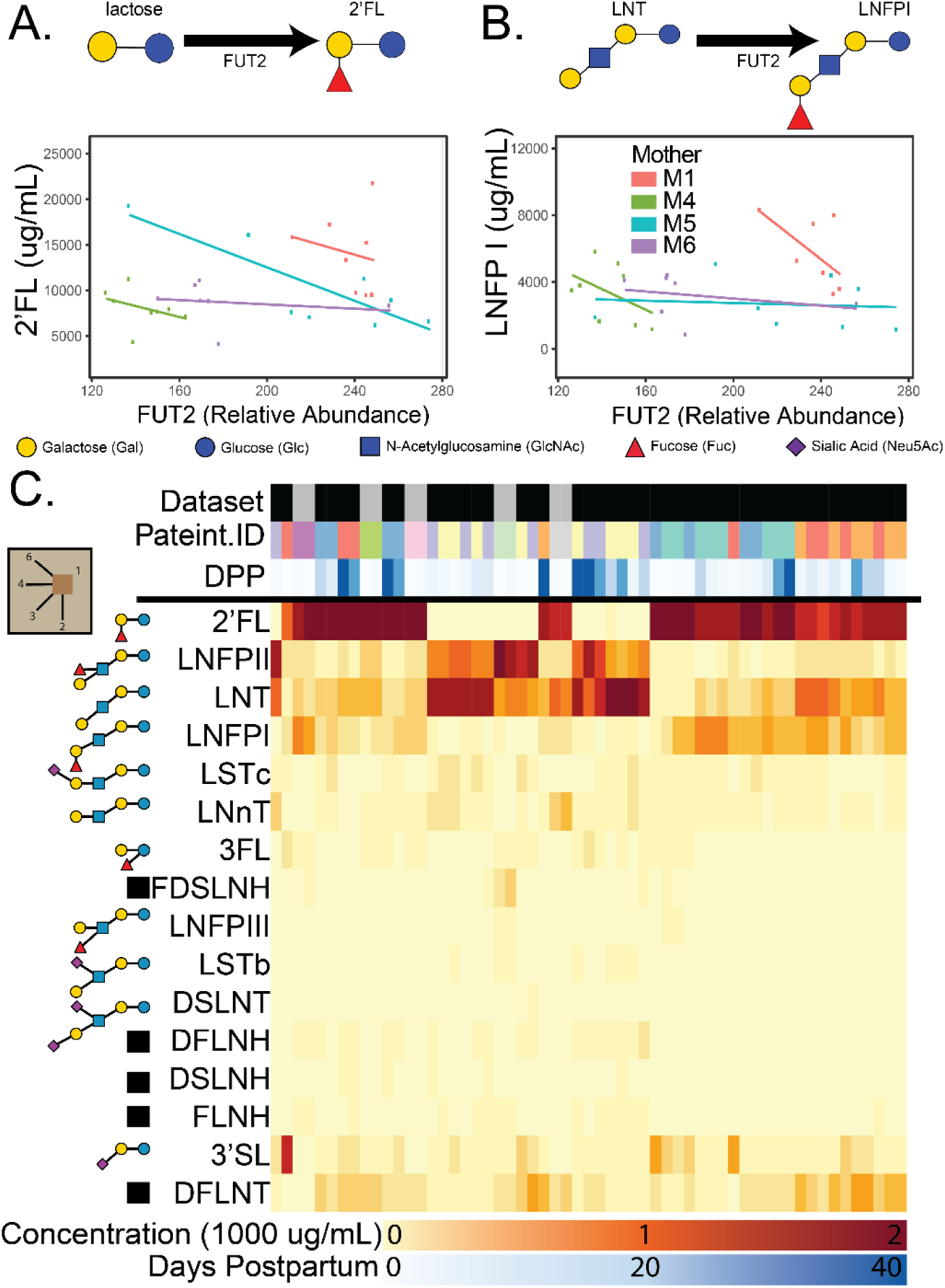
FUT2 expression should increase 2’FL and LNFPI which require the enzyme but there is no significant positive association. Direct comparison of FUT2 gene expression and concentrations (nmol/mL) of α-1,2-fucose containing HMOs, 2’FL **(A)** and LNFPI **(B)**, in sample-matched microarray and HPLC reveal no significant association in secretor women from cohort 1 sampled between day 1 and 42 post-partum. Trendlines and points are colored by subject. Linear trends were used to illustrate the intuition of the GEE approach used to estimate these associations across subjects. Non-secretor mothers were excluded due to non-functional FUT2. **(C)** A heatmap of all HMO concentrations across cohort 1 and cohort 2 (top-bar black and grey respectively). Known HMO structures are shown to the left of each row while uncharacterized structures are indicated with a black box. For proposed isomers of uncharacterized structures, see **Figure** S 12.

### 2.2 High-performing candidate biosynthetic models are supported by gene expression and predicted model flux across subjects

We built and examined models for HMO biosynthesis in human mammary gland epithelial cells. From the basic reaction set (**Figure** 3A), we generated the complete reaction network (**Figure** 3B) containing all possible reactions and HMOs with up to nine monosaccharides. The Complete Network was trimmed to obtain a Reduced Network (**Figure** 3D; **Figure** S 9, **Table** S 2) by removing reactions unnecessary for producing the observed oligosaccharides. Candidate models (**Figure** 3E) were built, capable of uniquely recapitulating the glycoprofiling data from milk using two independent cohorts--cohort 1 with 8 samples from 6 mothers between 6 hours and 42 days postpartum ^28,29^ and cohort 2 with 2 samples per mother on the 1st and second day after birth ^30^. Mixed integer linear programming was used to identify subnetworks with the minimal number of reactions from the Reduced Network. We identified 44,984,988 candidate models that can synthesize the measured oligosaccharides. Each candidate model contains 43-54 reactions (19.5-24.4% of the reactions in the Reduced Network (**Table** S 3)). These models covered all the feasible combinations of HMO synthesis by the 10 known glycosyltransferase families (**Figure** 1D) that could describe the synthesis of the HMOs in this study.

**Figure 3.**
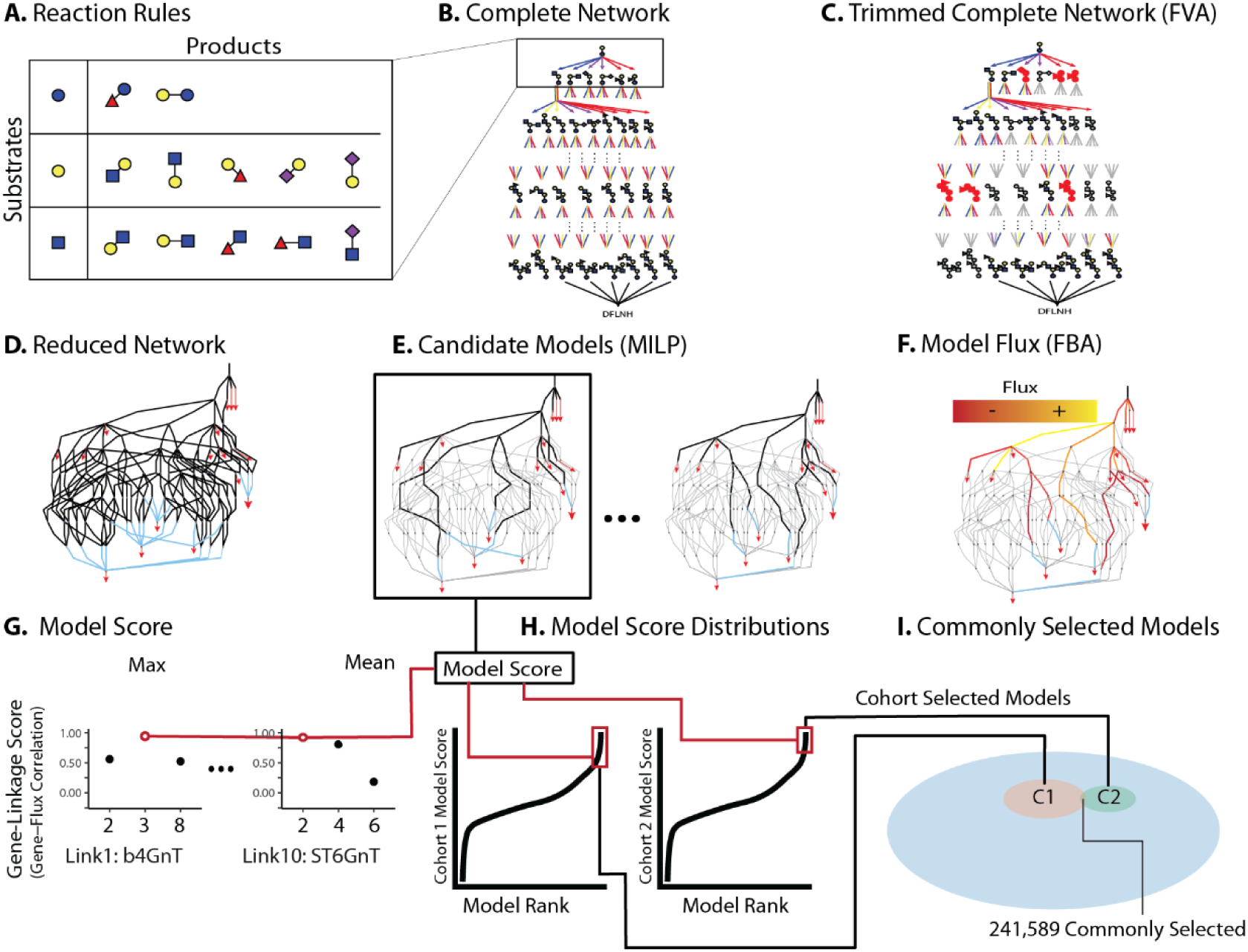
Overview of Computational Methods for Model Assembly (A-F) and Assessment (G-I). (A) To build the candidate models of HMO biosynthesis, reaction rules were defined to specify all possible monosaccharide additions. (B) The Complete Network includes all oligosaccharides and reactions resulting from the iterative addition of monosaccharides to a root lactose. (C) Using Flux Variability Analysis, the Complete Network was trimmed, removing reactions that cannot reach experimentally-measured HMOs, to produce a (D) Reduced Network **Figure** S 9; red triangles are observed HMOs blue lines are “sink reactions” joining alternative isomers (**Figure** S 12). (E) From the Reduced Network, Mixed Integer Linear Programming (MILP) was used to extract Candidate Models, each representing a subnetwork capable of uniquely synthesizing the observed oligosaccharide profile using a minimal number of reactions; black clines are reactions retained in a candidate model. (F) Flux Balance Analysis was used to estimate flux through each reaction necessary to simulate the measured oligosaccharide concentrations. (G) Model scores were computed as the average maximum correlation between linkage-specific candidate genes and normalized flux through that linkage (**Figure** S 11, S1.1.4). (H) Model scores were parameterized on cohort 1 (left) and cohort 2 (right) data (see Methods). High-performing models, 95th percentile of scores, are highlighted in red. (I) Of the >40 million models considered (blue), 2.66 and 2.32 million models were high-performing when parameterized on data from cohort 1 or cohort 2, respectively. Nearly 250,000 models consistently explained the relationship between predicted flux and expression data from both cohort 1 and cohort 2. These commonly selected models were analyzed for common structural features.

To identify the most likely biosynthetic pathways for HMOs, we computed a model score for each candidate model using the glycoprofiling and transcriptomic data from the two independent cohorts, after excluding low-expression gene candidates. Genes were excluded when expression was undetected in over 75% of microarray samples and the independent RNA-seq^31^ measured low expression relative to the GTEx^32^: TPM<2 and 75^th^ percentile Lemay < GTEx Median TPM. Specificity and expression filtration reduced the candidate genes from 54 to 24 (see supplemental results, **Table** S 1, **Figure** S 7); three linkages (L2, L5 and L9) were resolved by filtration alone indicating that FUT2, ST6GAL1 and FUT3 respectively perform these reactions.

Following low-expression filtering, we compared flux-expression correlation. Leveraging sample-matched transcriptomics and glycomics datasets, we computed model scores indicating the capacity of each candidate gene to support corresponding reaction flux. The model score was computed by first identifying for each reaction, the candidate gene that shows the best Spearman correlation between gene expression and normalized flux; flux was normalized as a fraction of the input flux to limit the influence of upstream reactions (**Figure** S 11, S1.1.4). The highest gene-linkage scores, for each reaction, for each model were averaged to obtain a model score (**Figure** 3G, see Methods). The model scores indicate consistency between gene expression and model-predicted flux. The high-performing models (z(model score)>1.646) were selected for further examination (**Figure** 3H, see Methods). Though quantile-quantile plots indicated the model score distributions were pseudo-gaussian, variation in skew resulted in slightly different numbers of high-performing models for the two different subject cohorts. Specifically, we found 2,658,052 high-performing models from cohort 1 and 2,322,262 high-performing models using cohort 2 (**Figure** 3I, **Table** S 4). We found 241,589 high-performing models common to cohort 1 and cohort 2. The model scores of commonly high-performing models are significantly correlated (Spearman R_s_=0.2, p<2.2e-16) and a hypergeometric enrichment of cohort 1 and cohort 2 selected models shows the overlap is significant relative to the background of 44 million models (p<2.2e-16). We analyzed these 241,589 commonly high-performing models and determined which candidate genes were common in high-performing models.

To determine the most important reactions (**Figure** 4) in the reduced network, we asked which reactions were most significantly and frequently represented among the top 241,589 high-performing models. We then filtered to retain only the top 5% of most important paths from lactose to each observed HMO (see Methods). The most important reactions form the Summary Network (**Figure** 4). Here, HMO biosynthesis naturally segregates into type-I backbone structures, with β -1,3-galactose addition to the GlcNAc-extended core lactose, and type-II structures, with β -1,4-galactose addition to the GlcNAc-extended core lactose. As expected, LNFPI, LNFPII, LSTb and DSLNT segregate to the type-I pathway while LNFPIII and LSTc are found in the type-II pathway (see Methods for HMO definitions). The Summary Network suggests resolutions to large structurally ambiguous HMOs (FLNH5, DFLNT2, DFLNH7, and DSLNH2) by highlighting their popularity in high-performing models. The Summary Network also shows three reactions of high comparable strength projecting from GlcNAc-β1,3-lactose to LNT, LNnT and a bi-GlcNAc-ylated lactose (HMO8, **Figure** 4, **Table** S 2) suggesting LNT may be bypassed through an early β-1,3-GlcNAc branching event; a previously postulated alternative path^33^. We checked for consistency with previous work^34^ and found that (1) the single fucose on the reducing-end Glc residue is always α-1,3 linked, (2) for monofucosylated structures, the non-reducing terminal β-1,3-galactose is α-1,2-fucosylated, (3) all galactose on the β-1,6-GlcNAc is always β-1,4 linked while all galactose on the β-1,3-GlcNAc are either β-1,3/4 linked. With the exception FDSLNH1, (4) no fucose is found at the reducing end of a branch and (5) all α-1,2-fucose appear on a β-1,3-galactose and not β-1,4-galactose in monofucosylated structures with more than four monosaccharides; suggesting that FDSLNH1 is an unlikely isomer. The summary network also suggests that most HMOs have type-I LacNAc backbones. To address the potential over-representation of type-I HMOs in our models, we examined the distribution of type-I and type-II in tetra- and pentasaccharides with known structures. Across samples, the median abundance of type-II HMOs, LNnT, LNFPIII and LSTc were 3.33%, 0.041%, and 2.68% of total nmol/mL while type-I HMOs of the same size, LNT, LNFPI, LNFPII, and LSTc, was 15.3%, 9.39%, 7.45% and 0.45% respectively. This confirms the greater abundance of type-I HMOs compared with the type-II structures in the glycomic profiles (**Figure** 2C). This Summary Network thus provides orientation in this underspecified space.

**Figure 4.**
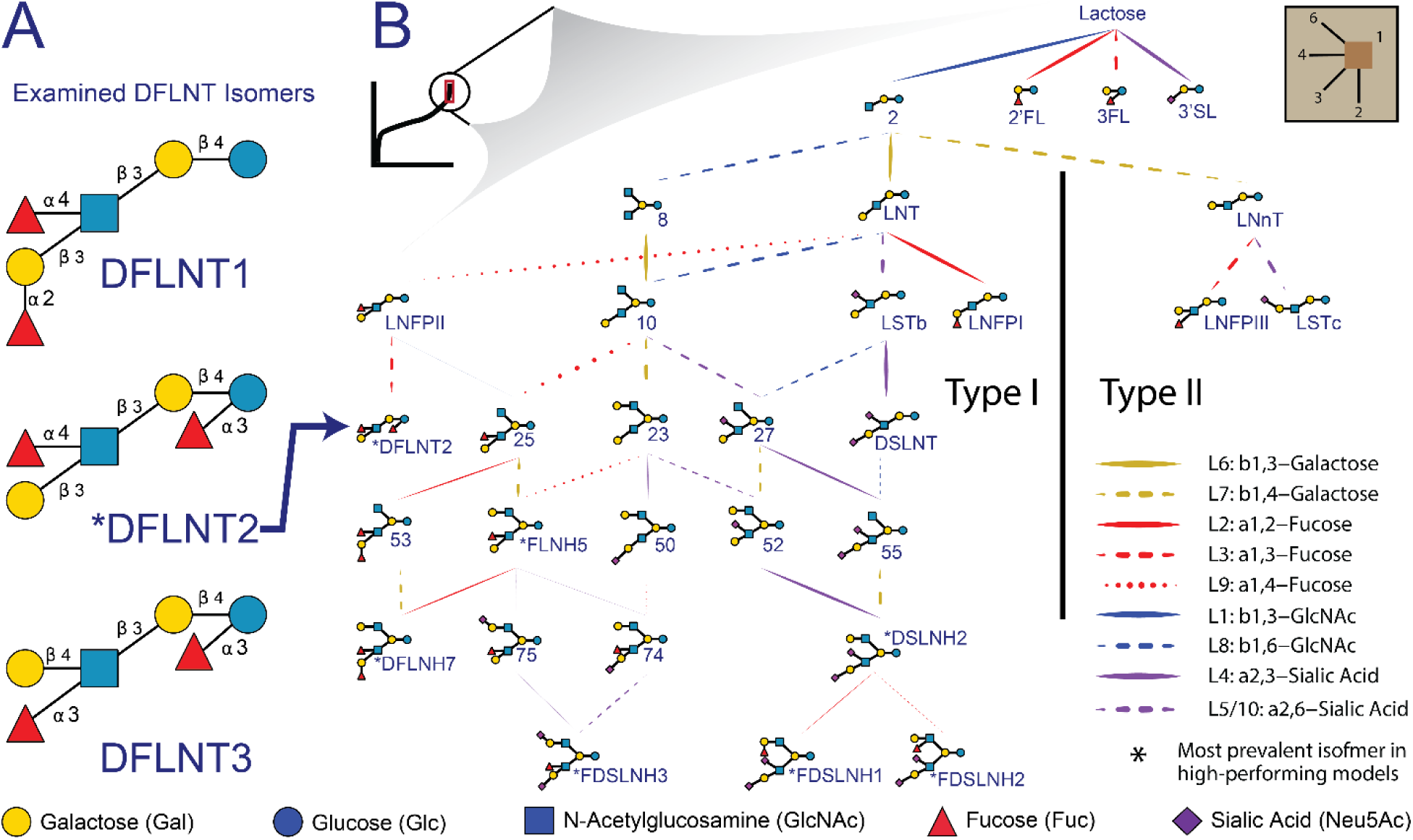
Summary Network of the most important reactions in the reduced network. Observed, intermediate and candidate HMOs most important to commonly high-preforming networks were selected from the Reduced Network (**Figure** 3D; Supplemental Methods S1.1.2). (A) Several ambiguous isomers (**Figure** S 12) were preferred(**Figure** S 1) in the commonly high-performing models. (B) A summary network was constructed from reaction importance; an aggregation of the proportion of high-performing models that include a reaction, and the enrichment of a reaction in the high-performing model set (see Methods). Line weight indicates the relative importance of each reaction. Line color corresponds to the monosaccharide added at each step and line type corresponds to the linkage type. The Summary Network naturally segregates into type-I and type-II backbone structures. For measured HMO definitions (e.g. FDSLNH and DSLNT) see Methods, for intermediate HMO definitions (e.g. 8, 10, or 25) see **Table** S 2, for uncertain structures (e.g. DFLNH7, FLNH5) see **Figure** S 12.

### 2.3 Glycosyltransferases are resolved by ranking reaction consistency across several metrics

We further analyzed the high-performing models to identify the glycosyltransferases responsible for each step in HMO biosynthesis (**Table** 1). As previously described, not all members of a gene family were examined in this analysis. Some genes were excluded due to their well characterized irrelevance (e.g. FUT8) and others, like FUT1, were excluded due to low expression in lactating breast epithelium (see **Table** S 1, methods and supplemental results for the detailed inclusion criteria). To determine the genes preferred for each reaction, we used three metrics to quantify the association between candidate gene expression and predicted flux. These were (1) *proportion (PROP -* the relative proportion of models best explained by a candidate gene, **Figure** S 4), (2) *gene linkage score (GLS* - the average Spearman correlation between gene expression and flux), and (3) *model score contribution (MSC -* an estimate of the gene-influence indicated by the Pearson correlation between model score and gene linkage score) (**Figure** 5A, **Figure** S 5). For each candidate gene, we generated a reaction support score (**Figure** 5B, see Methods); the pooled significance of the maxima of PROP, GLS and MSC across both cohorts.

**Figure 5.**
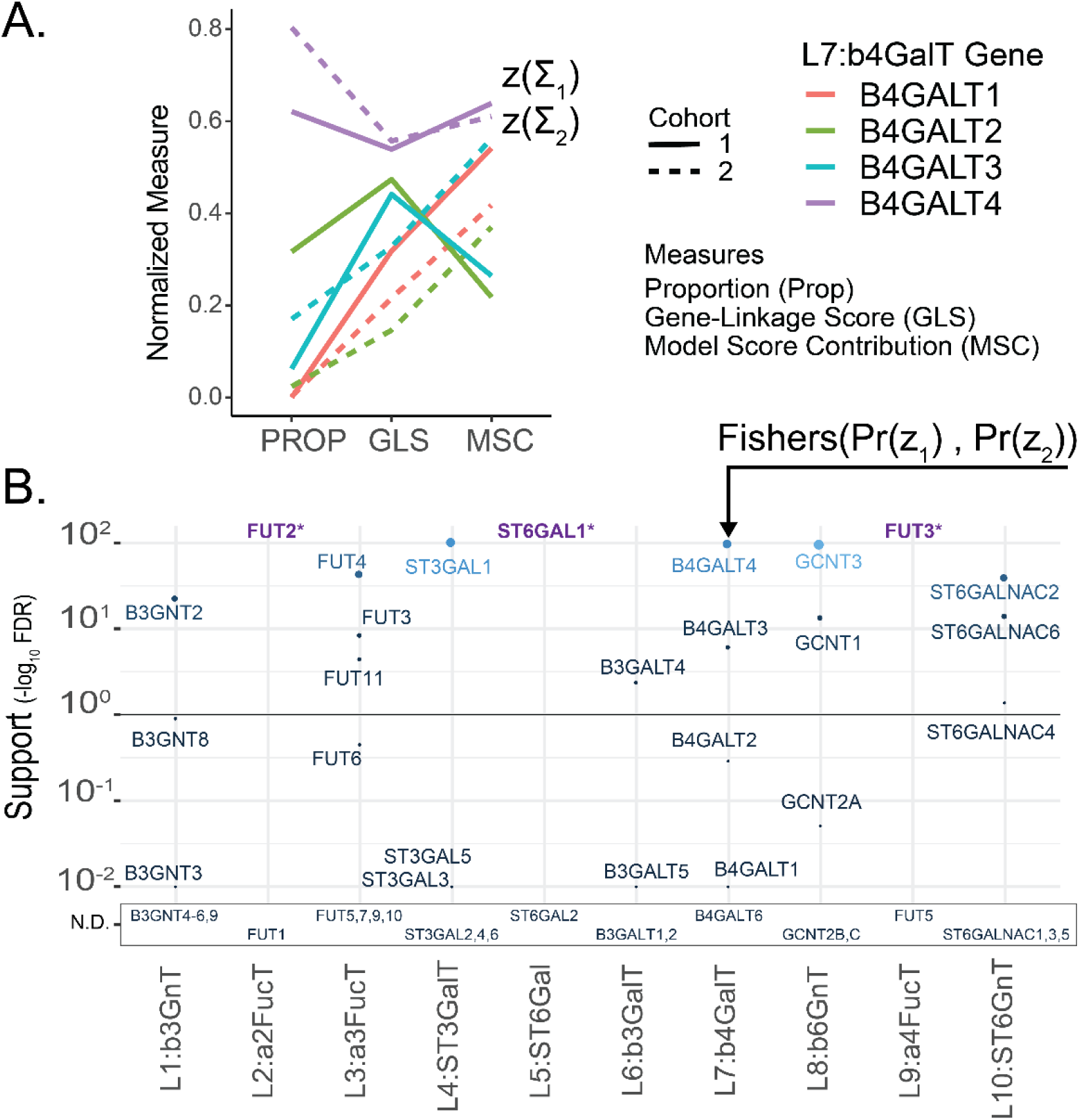
Gene expression correlation with model flux predicts enzymes involved in HMO biosynthesis. (A) To determine the gene expression that best explains flux through each reaction in each glycomics-transcriptomics matched sample, we examined the proportion of high-performing models were each gene was most flux-correlated (PROP, **Figure** S 4), we also examined the gene-linkage score (GLS) and Model Score Contribution (MSC). For this visual, each measure was max-min normalized between 0 and 1. Genes were selected based on high performance on all three measures across cohorts (line type). (B) We summarize the three performance scores from panel A across cohorts into a single support score (see Methods). Briefly, “Support” is p-value for the sum of PROP, GLS and MSC z-scores (relative to a permuted background), Fisher-pooled across cohorts then False Discovery Rate (FDR) corrected across genes (see Methods). Unmeasured genes appear below the plot in the Not Determined (N.D.) box. Genes selected by default (purple, “*”) as the only measured gene candidate (**Table** 1)

Three reactions, L2 (FUT2), L5 (ST3GAL1) and L9 (FUT3), were matched to genes by default as they were the only gene candidates remaining following gene expression filtering (**Table** S 1, Supplementary Results). At least one gene showed significant support (q<0.1) for each remaining reaction. GCNT3 shows highly significant support (q<0.001) and nearly 100% of models selected this isoform over GCNT2C or GCNT1 (**Figure** S 4). B4GALT4 is the most significantly supporting gene for the L7: b4GalT reaction (**Figure** 5B). In both cohort 1 and 2, B4GALT4 outperforms all other isoforms in all three metrics. B4GALT4 expression best explains flux in 62% and 80% (PROP) of high-performing models using cohort 1 and 2 data respectively (**Figure** S 4). B4GALT4 also has the highest MSC and GLS (z>5.6) of any isoforms. Interestingly, while B4GALT1 is highly expressed and fundamental to lactose synthesis in the presence of α-lactalbumin and lactation in general^35,36^, it showed negligible support for the L7 reaction (**Figure** 5B). Considering the reaction support score, all linkages show at least one gene for each reaction that significantly explains behavior across cohorts (**Figure** 5B).

### 2.4 Kinetic assays confirm selected genes and expand our scope

Towards validating and expanding our gene-reaction predictions, glycosyltransferase enzyme activity assays were performed using the NTP-Glo™ Glycosyltransferase assay format from Promega. We used linkage L1:b3GnT and L10:ST6GnT to validate our selections and examined every plausible isoform of the ST3GAL for its ability to perform the linkage L4:ST3GalT reaction. Five acceptors were used: (1) lactose to examine activity on the initial HMO acceptor, (2) LNT and (3) LNnT to establish which enzymes would act on larger type-I and type-II tetrasaccharides, (4) Gal β1,3-GalNAc to determine specificity for non-HMO O-type glycans, and (5) a GlcNAc-β1,3-Gal-β1,4-GlcNAc-β1,3-Gal-β1,4-Glc pentasaccharide structure to test the formation of a non-reducing terminal type-I (Gal-b1,3-) cap on a longer acceptor. We explored the activities of various gene products to perform specific glycosyltransferase reactions crucial to HMO biosynthesis (**Figure** 6, **Table** S 5).

**Figure 6.**
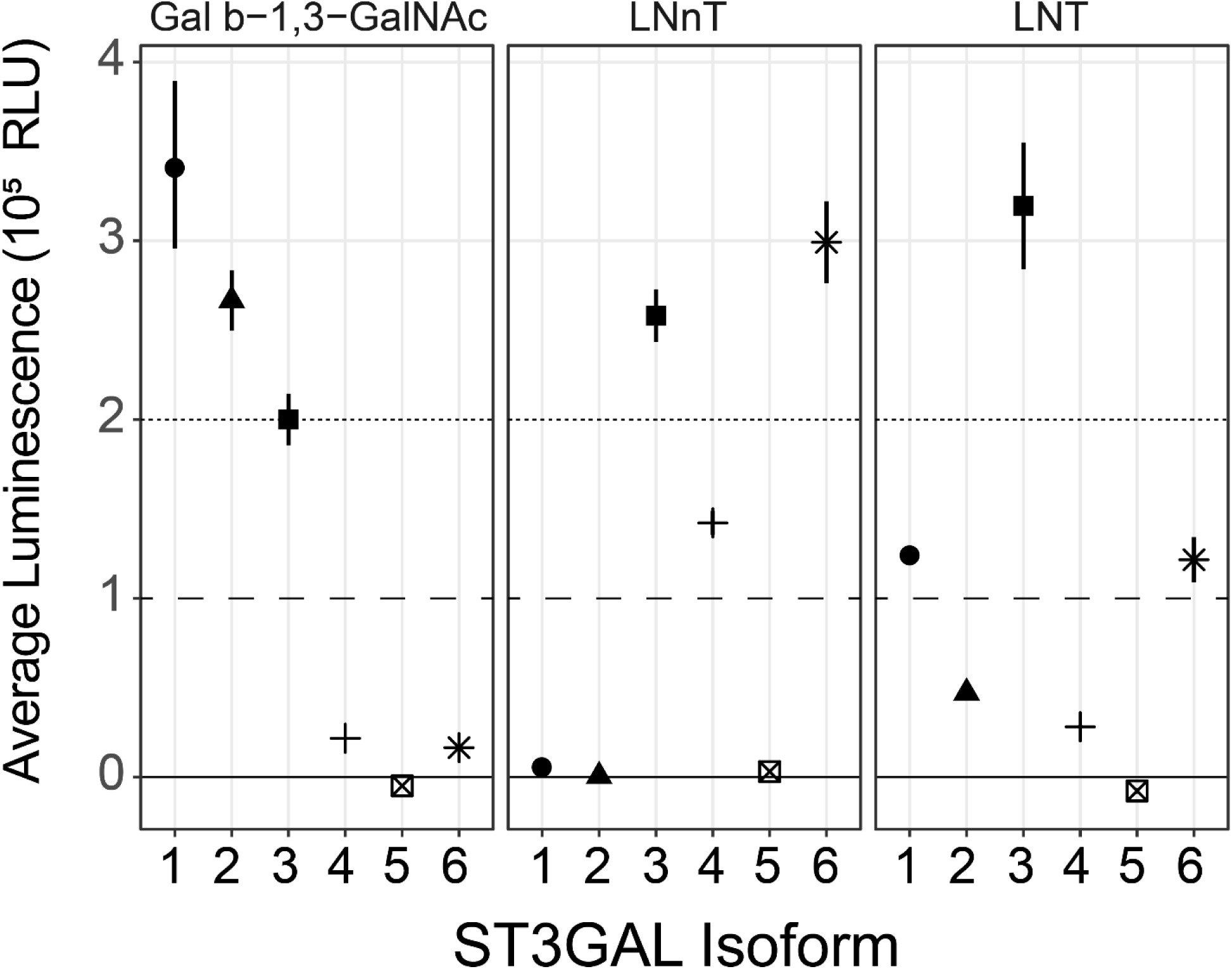
Results of the CMP-Glo™ Glycosyltransferase Assay to test GT candidates on relevant HMO acceptors. Average luminescence below 10,000 is considered weak activity, and activity above 200,000 is considered very high activity. Reported luminescence values were background corrected and 95% confidence intervals are shown. For complete details see **Table** S 6. Shapes correspond to ST3GALT isoforms

In the cross-cohort aggregate analysis (**Figure** 5B), B3GNT2 is selected as a reasonable candidate to catalyze flux through the L1:b3GnT reaction. The B3GNT2 support score is nearly 100 times more significant than B3GNT8, the next most associated gene. Consistent with the predictions that b3GnT should convert lactose into the precursor to LNT and LNnT, the UDP-Glo™ assay showed B3GNT2 had high activity toward lactose as an acceptor. We further found that B3GNT2 could add a β1,3-GlcNAc to LNnT as is necessary for poly-lacNAc HMOs. The cross-cohort aggregate analysis (**Figure** 5B) selected ST6GALNAC2 to perform L10, the α2,6 addition of sialic acid to the internal β1,3-GlcNAc; necessary for the biosynthesis of LSTb from LNT and possibly DSLNT from LSTa. However, the CMP-GLO™ assay highlighted a negligible activity of ST6GALNAC2 toward LNT even at very high enzyme input indicating that this enzyme does not convert LNT to LSTb. We did not test if it can convert LSTa to DSLNT. In contrast, ST6GALNAC5 was effectively able to use LNT as an acceptor, although we did not confirm the formation of the LSTb structure. ST6GALNAC5 could not be considered in the support score calculation because it was only measured in cohort 2; expression was greater than zero in 1 of 12 samples.

Finally, we tested the affinities of plausible ST3GAL isoforms to sialylate LNT, LNnT or β1,3-GlcNAc (**Table** S 5). The multi-cohort analysis (**Figure** 5B) implicates ST3GAL1 as the best candidate for this reaction. The CMP-Glo™ assay indicated that ST3GAL1 has limited activity toward LNT but high activity toward Gal β1,3-GlcNAc suggesting ST3GAL1, *in vitro*, is more involved in non-HMO O-type glycan biosynthesis. ST3GAL2 showed a similar but less substantial pattern. ST3GAL3 showed the strongest activity for sialylation both LNT and LNnT suggesting it could synthesize LSTa from LNT. ST3GAL6 shares a similar but lesser activity for LNT and LNnT.

We analyzed the original expression profiles to determine which genes were sufficiently expressed to actuate this activity. STGAL1, 3 and 5 were strongly expressed in nearly 100% of samples across both cohorts; ST3GAL2 and 4 show zero expression in 75% of samples in at least one cohort (**Figure** S 6). ST3GAL3 was highly expressed and effective at catalyzing the L4 reaction for LNT and LNnT while ST3GAL1 was highly expressed and weakly catalyzed sialylation of LNT making ST3GAL3 the most likely candidate for L4 reaction on LNT and LNnT.

### 2.5 Selected glycosyltransferases share transcriptional regulators across independent predictions

To explore the transcriptional regulation during lactation, we used two orthogonal approaches for transcription factor (TF) discovery. We used Ingenuity Pathway Analysis (IPA) to predict upstream regulatory factors based on differential expression associated with each HMO. IPA analyzed all genes differentially expressed with HMO abundance, not only HMO glycogenes; these differential expression patterns formed HMO specific gene expression signatures. Additionally, we used MEME for *de novo* motif discovery in the promoter regions of HMO glycogenes and TOMTOM to map those discovered motifs to known TFs. We validated these predictions by examining transcriptional regulators selected by both MEME and IPA (**Figure** S 13, see Methods).

IPA discovered 57 TFs significantly (|z|≥3; p < 0.001) associated with the 16 HMO-specific gene expression signatures. We performed differential expression on HMO substructure abundance and substructure abundance ratios^17^; IPA found 66 and 49 TFs significantly (|z|≥3; p < 0.001)) associated with HMO substructure and substructure ratio specific gene expression signatures. Using MEME, we identified three putative TF regulatory sites (TF motifs I, II and III) for 6 selected glycosyltransferases responsible for the HMO biosynthesis (**Table** 2 and **Figure** S 15). TOMTOM calculated that these putative binding sites were significantly associated with six known TFs (IKZF1, SP1, EGR1, ETS1, ETV4 and ERG) that were also predicted by IPA as regulators of gene signatures associated with HMO concentration (**Figure** 7, **Figure** S 16) or HMO glycan substructures abundance (**Figure** S 17). SP1, EGR1, ETS1, ETV4 and ERG are all predicted to positively influence expression associated with the biosynthetically related HMOs: 3’SL, 3FL, LSTb and DSLNT; 3’SL and 3FL share a common substrate (lactose) while LSTb is a likely precursor to DSLNT. The motif-level analysis showed opposing regulation between IKZF1: upregulating gene expression signatures associated with the 3’SL and LSTb substructure abundance^17^ (X34 and X62 respectively, see **Figure** S 19) and downregulating gene expression associated with GlcNAC-lactose, LNT and LNFPI substructure abundance (X18, X40 and X65 respectively, see **Figure** S 19), while EGR1, ERG and ETS1 have the opposite predicted impact (**Figure** S 17). The motif-level predictions are consistent with the HMO-level predictions of upregulation on 3’SL and LSTb while adding an additional point of contrast. While EGR1, ERG and ETS1 are predicted to increase production of sialylated HMOs, they may have the opposite impact on LNFPI. Thus, we detect signatures of multiple transcription factors that could coordinate the regulation of the genes we identified to contribute to HMO biosynthesis (see supplemental discussion).

**Table 2.** TF motif (MEME) and IPA upstream regulator integrated results.

| LINKAGE | REACTION<br>SUPPORT<br>SCORE<br>SELECTED<br>CANDIDATE | MEME<br>TF MOTIF<br>(P-VALUE)*1 | JASPAR TF<br>(P-VALUE)*2 | IPA PREDICTED<br>TF*3 |
| --- | --- | --- | --- | --- |
| L1:B3GNT | B3GNT2 | TF Motif-II<br>(1.39e-12) | SP1 (4.96e-05)<br>EGR1 (2.17e-05) | Y<br>Y |
| L2:A2FUCT | FUT2 | TF Motif-III<br>(3.63e-16) | IKZF1 (7.62e-04) | Y |
| L3:A3FUCT | FUT11 | TF Motif-II<br>(1.00e-7) | SP1 (4.96e-05)<br>EGR1 (2.17e-05) | Y<br>Y |
| L4:ST3GAL<br>T | ST3GAL1 | TF Motif-I<br>(2.76e-11) | ETV4 (1.24e-03)<br>ETS1 (3.01e-04)<br>ERG (3.50e-04) | Y<br>Y<br>Y |
| L7:B4GALT | B4GALT4 | TF Motif-II<br>(7.67e-11) | SP1 (4.96e-05)<br>EGR1 (2.17e-05) | Y<br>Y |
| L10:<br>ST6GNT | ST6GALNAC2 | TF Motif-II<br>(1.08e-7) | SP1 (4.96e-05)<br>EGR1 (2.17e-05) | Y<br>Y |

\*1. The P-value (see **Figure S 15**) is the significance of the selected GT to the MEME identified TF motif.
\*2. The P-value (see **Table S 7**) is the significance of known TF associated with the MEME identified TF motif.
\*3. The IPA upstream regulator analyses were conducted on the three different sets of DEGs: 16 HMOs, 19 glycan motifs, and 4 differential motifs (see Methods for details). Based on the Z-score predicted by IPA using the gene expression data, we selected the significant TFs with IPA predicted activation score $|Z \text{ value}| \geq 3$ in this study. Note, 'Y' denotes the known TF is presented in the indicated dataset (HMO (**Figure 7**, **Figure S 16**), Motif (**Figure S 17**), or differential motif (**Figure S 18**)) of the IPA predicted TF and 'N' means the TF doesn't present in the dataset of IPA predicted TF.

**Figure 7.**
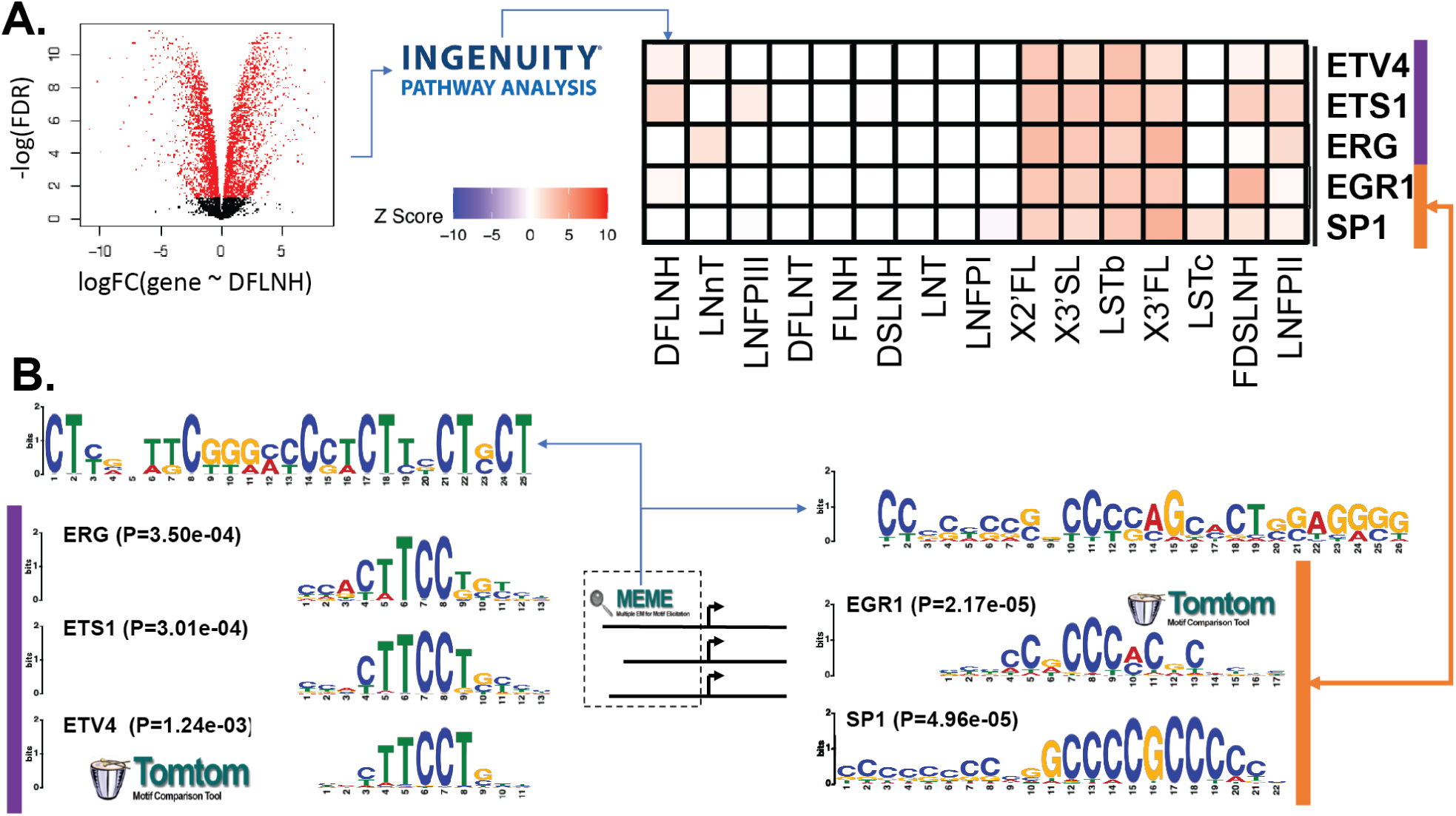
de novo promoter-enriched TF motifs and IPA predicted TFs using differential expression analyses with respect to 16 HMOs. (A) MEME identified TF motifs and 5 known TFs (ETV4, ETS1, EGR1, SP1, and ERG) associated with them (see **Table** S 6). MEME-discovered TFs were cross-referenced with known TF binding sites using TOMTOM. Logos for the matched known and discovered motifs are shown in the top and bottom of each subpanel; the p-value is a logo matching significance calculated by TOMTOM. (B) Subset of a biclustering of activation z-score computed by IPA indicating the likelihood that a TF activates (z>0) or inhibits (z<0) an HMO concentration signature (gene expression associated with changes in HMO concentration). The full biclustering can be found in the supplement (**Figure** S 16)

## 3 Discussion

By integrating sample-matched quantitative oligosaccharide measurements and gene expression data using computational models of HMO biosynthesis, we resolved genes responsible for 10 elementary reactions in human mammary gland epithelial cells. The modeling-based strategy was essential since simple correlations failed to capture the simplest HMO-gene associations, given the complex interactions of glycosyltransferases in the HMO biosynthetic pathway. Because the pathway characterization is still incomplete, we built >44 million candidate models that uniquely recapitulate glycoprofiling data in two independent cohorts. Candidate model flux, i.e. activity of each reaction, was predicted for each model and compared to sample-matched gene expression data. We used the consistency between gene expression and predicted flux across cohorts in high-performing models to select genes for each fundamental reaction. Analysis of these models suggested glycosyltransferase genes, thus providing a clearer picture of the enzymes and regulators of HMO biosynthesis in mammary epithelial cells. The clarification of the pathways and enzymes involved in HMO biosynthesis will be an invaluable resource to help (1) discover the maternal genetic basis of health-impacting^1,2,5,6,37–46^ HMO composition heterogeneity^7,26,47,48^ and (2) drive chemoenzymatic synthesis ^49–53^ and metabolic engineering for manufacturing HMOs for food ingredients, supplements and potential therapeutics^54–59^ (see supplemental discussion).

Of the three fucosylation reactions, two were determined using expression data alone while the third required additional insight from the flux-expression comparison or, support score. Consistent with studies in blood^23–25^ and milk^26,47,60^ types, we selected FUT2 as the gene supporting the α1,2-fucosylation (L2:a2FucT) linkage reaction. FUT1 was ruled out due to non-expression (**Table** S 1, supplemental results). In the second fucosylation reaction, FUT3, FUT4 and FUT11 all show significant support for α1,3-fucosylation (L3:a3FucT) linkage formation. FUT11 is more commonly considered an N-glycan-specific transferase^61^ and therefore a less likely candidate. Both FUT3 and FUT4 prefer to fucosylate the inner GlcNAc of a type-I polylactosamine^62^. FUT3 prefers neutral type-I polylactosamine while FUT4 also fucosylates the sialylated form^63,64^; the charge preferences are inverted for type-II polylactosamine acceptors^65^. Prudden et. al.^52^ used FUT9 to perform this reaction, consistent with its ability to transfer α1,3 fucose to the distal GlcNAc of a neutral polylactosamine^61–63^. The four HMO structures with α1,3-Fucose in the Summary Network (**Figure** 4) include 3FL (neutral inner fucosylation), LNFPIII (neutral distal fucosylation), DFLNT2 (neutral inner fucosylation), and FDSLNH2 (sialylated and neutral distal fucosylation). FUT9 showed negligible expression in RNA-Seq (3^rd^ Quartile TPM=0.37, **Table** S 1), yet it is highly expressed (TPM>10) brain and stomach^32^. Therefore, it is likely that the distal fucosylation is conducted by another enzyme *in vivo* while the inner fucosylation is likely performed by either FUT3 or FUT4. FUT3 was also chosen for the α1,4-fucosylatoin (L9:a4FucT) by default due to the non-expression of FUT5, confirmed by RNA-Seq (**Table** S 1, supplemental results). FUT3 adds an α1,4-fucose to the GlcNAc of a neutral type-I chain to form the Lewis-A or Lewis-B group and adds an α1,3-fucose to the GlcNAc of a type-II chain^63,64^. Usage of FUT3 would provide a parsimonious explanation for the fucosylation of both type-I and type-II HMOs like LNFPII (Fuc-α1,4-LNT (type-I)) and LNFPIII (Fuc-α1,3-LNnT (type-II)).

One of two sialyltrasferases was clearly resolved with expression data alone, the other required additional examination. ST6GAL1 was chosen by default to support the α2,6-sialylation (L5:ST6GalT) reaction due to the non-expression of ST6GAL2 (**Table** S 1). ST6GAL1 sialylates galactose in HMOs^52^. For the second sialylation reaction, our flux-expression comparison selected ST6GALNAC2 and ST6GALNAC6 as the significant supporters of α2,6 sialylation (L10:ST6GnT). Through a kinetic assay, we confirmed that ST6GALNAC2 (previously shown to accept core-1 O-glycans^66,67^) fails to sialylate LNT. Though our kinetic assay shows that ST6GALNAC5 (known to sialylate GM1b^68^) can sialylate LNT, it was not expressed in this context (**Table** S 1, supplemental results). ST6GALNAC3 expression was not observed in microarrays but could not be ruled out due to RNA-Seq expression (**Table** S 1, supplemental results); it sialylates the GalNAc of NeuAc-α2,3-Gal-β1,3-GalNAc-α1-O-Ser/Thr and NeuAc-α2,3-Gal-β1,3-GalNAc-β1,4-Gal-β1,4-Glc-β1-Cer when the inner galactose is not sialylated (e.g. GD1a or GT1b)^69–72^ but has not been shown to transfer to a GlcNAc. The last ganglioside-accepting family gene, ST6GALNAC6, has broader activity accepting several gangliosides (GM1b, GD1a, and GT1b)^69^ and sialylating the GlcNAc of LNT-ceramide^73^. Considering the broader activity, clear expression and computational selection, ST6GALNAC6 is the most likely candidate, though ST6GALNAC3 should not be ruled out. In the third reaction, ST3GAL1 shows significant support for α2,3-sialylation (L4:ST3GalT) reactions while ST3GAL3 shows negligible consistency in the flux-expression comparison. Yet, *in vitro*, ST3GAL3 was most effective at sialylating both LNT and LNnT in kinetic assays while ST3GAL1 weakly sialylated LNT. ST3GAL4, which prefers type-II acceptors^74– 76^, was used previously to perform this reaction *in vitro*^52^, but it was not expressed on the microarrays nor RNA-Seq. ST3GAL3 can accept type-I, type-II and type-III acceptors including LNT and prefer type-I acceptors^74,75,77^ while ST3GAL1 accepts type-I, type-III and core-1 acceptors but not type-II^74,75,78^. The kinetic assays and previous literature show ST3GAL3 is more capable than ST3GAL1 at catalyzing this reaction, while ST3GAL1 expression was found to be the only plausible candidate based on estimated flux through this reaction. If ST3GAL1 were responsible for this reaction, its inability to sialylate type-II HMO could partially explain the lack of sialylation and larger structures in the type-II HMO branch. Both ST3GAL1 and ST3GAL3 remain plausible candidate genes, and further *in vivo* studies are needed. Both galactosylation reactions required further examination of flux-expression relationships. We found B3GALT4 to significantly support the type-I β -1,3-galactose addition (L6:b3GalT). B3GALT4 can transfer a galactose to GalNAc in the synthesis of GM1 from GM2^79^. Unlike B3GALT5, there is no evidence that B3GALT4 can transfer galactose to a GlcNAc^80^. B3GALT5, has been shown to transfer a β -1,3-galactose to GlcNAc to form LNT *in vitro*^81^. B3GALT5 expression measured for cohort 1 microarray was much lower than expression in cohort 2 and the independent RNA-Seq^31^ suggesting that the probes in the first microarray may have failed (**Table** S 1, supplemental results). While both B3GALT4 and B3GALT5 seem plausible, given the historical failures of B3GALT4 to perform this reaction and our likely failure to measure and evaluate B3GALT5, B3GALT5 may be the stronger candidate for this reaction. In the second galactosylation reaction, the flux-expression comparison found B4GAL4 and B3GALT3 most significantly supports the type-II definitive β-1,4-galactose addition (L7:b4GalT). These gene-products can synthesize LNnT-ceramide^82^. Additionally, in the presence of α-lactalbumin (highly expressed during lactation), B4GALT4 shows an increased affinity for GlcNAc acceptors suggesting during lactation it is more likely to perform the L7 reaction^82,83^. B4GALT1 and B4GALT2 synthesize lactose in the presence of α-lactalbumin during lactation^35,36^, but B4GALT1 expression was not correlated with L7 flux and B4GALT2 was not expressed (**Table** S 1). We note that associations between B4GALT1 expression L7 flux may be masked due to its consistent high. Therefore, flux-expression correlation should not be used to exclude B4GALT1 as a candidate for the L7 reaction. Doing so, B4GALT4, B4GALT3 and possibly B3GALT1 remain the most plausible candidates.

Finally, both GlcNAc additions required flux-expression examinations. B3GNT2 showed significant support in the flux-expression comparison. In our kinetic assays, B3GNT2 demonstrated high activity towards lactose as an acceptor. Previously, B3GNT2 has performed the β-1,3-GlcNAc addition (L1:b3GnT) on multiple glycan types including several HMOs: lactose, LNnT, polylactosamine-LNnT^84^. The agreement of literature, kinetic assays and flux-expression analysis indicate B3GNT2 is an appropriate choice for this reaction. In the second GlcNAc reaction, GCNT3 and GCNT1 most significantly support the branching β-1,6-GlcNAc addition (L8:b6GnT). While GCNT2B can effectively transfer the branching GlcNAc to the inner galactose of LNnT^52,85^, it was not expressed in the cohort microarrays or independent RNA-Seq. GCNT1 transfers a branching GlcNAc to the GalNAc of a core-1 O-glycan^86,87^ while GCNT3 acts on core-1 and the galactose of the LNT-like core-3 structure^88,89^. GCNT3 is also specifically expressed in mucus-producing tissues^88,89^ like lactating mammary gland epithelium. Interestingly, GCNT3 acts on galactose of the GlcNAc-β1,3-Gal-β1,4-Glc trisaccharide (predistally) while GCNT2 acts on the central galactose of the LNnT or LNT tetrasaccahride (centrally)^85^. Therefore, reliance on GCNT3 for the branching reaction would explain the noncanonical branched tetrasaccharide (HMO8, **Figure** 4) suggesting a third major branch from GlcNAc-β1,6-lactose, distinct from LNT and LNnT. Predistal addition of the branched GlcNAc may also explain the lack of branched type-II structures since B4GALT4 cannot act on branched core-4 structures^90^. HMO biosynthesis with GCNT3 and B4GALT4 could explanation the type-I bias seen in the Summary Network (**Figure** 4).

Our results show consistency with experimental validation here and the published literature. Further direct empirical studies will be invaluable to confirm each gene-reaction association and the complete biosynthesis network. Such studies would include further clinical cohort studies and the development of mammary organoid models capable of producing HMOs. Such experimental systems can clarify the impact of mammary-tissue specific genes, cofactors, and HMO chaperones like α-lactalbumin ^82,83^ on glycosyltransferase activity. Therefore, further development of authentic *in vitro* cell and organoid models will be invaluable to finalizing our model of HMO biosynthesis.

## 4 Conclusion

By using systems biology approaches, different omics data can be integrated, as shown here to predict gene-reaction relations even in highly uncertain and underdetermined networks. Of the ten fundamental reactions we aimed to resolve and reduce (**Table** 1), we succeeded in narrowing the candidate substantially for each one. The newly reduced space of HMO biosynthetic pathways and knowledge of the enzymes and their regulation will enable mechanistic insights into the relationship of maternal genotype and infant development. Finally, once essential HMOs are identified, the knowledge presented here on the HMO biosynthetic network can provide insights for large-scale synthesis of HMOs as ingredients, supplements, or potential therapeutics to further help improve the health of infants, mothers, and people of all ages.

## 5 Author contribution

BK, AR, LB and NEL designed and performed the study and wrote the manuscript. ABB performed preliminary analysis. AR performed modeling analyses. BK analyzed and interpreted the models analyses. MAM and MWH provided samples. DC, JYY, JN, KM and LB performed expression, purification of glycosyltransferases and kinetic assays. AR, BK, and NK performed literature surveys to determine appropriate candidate genes for each reaction. BK and BB performed motif-level analysis. BK and AWTC performed transcription factor analysis.

## Supporting information

Table S1 - Expression

Table S2 - Parameterized Network

## 6 Acknowledgments

Special thanks to Frederique Lisacek and Andrew McDonald for their input on navigating this interdisciplinary topic. Additional thanks to Philip Spahn, Hooman Hefzi, Krystyna Kolodziej and Caressa Robinson for help editing this manuscript. This work was supported by a Lilly Innovation Fellows Award (A.R.), the Novo Nordisk Foundation provided to the Center for Biosustainability at the Technical University of Denmark (NNF10CC1016517: N.E.L.), NIGMS (R35 GM119850: N.E.L., P41GM103390, P01GM107012, and R01GM130915 to K.W.M.), NICHD (R21 HD080682: L.B.) and USDA (USDA/ARS 6250-6001; M.W.H). This work is a publication of the U.S. Department of Agriculture/Agricultural Research Service, Children’s Nutrition Research Center, Department of Pediatrics, Baylor College of Medicine, Houston, Texas. The contents of this publication do not necessarily reflect the views or policies of the U.S. Department of Agriculture, nor does mention of trade names, commercial products, or organizations imply endorsement from the U.S. government.

## 7 Materials and methods

### 7.1 Milk sample Collection

Samples were collected following Institutional Review Board approval (Baylor College of Medicine, Houston, TX). Lactating women 18-35 years of age with uncomplicated singleton pregnancy, vaginal delivery at term (>37 weeks), Body Mass Index <26 kg/m^2^ without diabetes, impaired glucose tolerance, anemia, or renal or hepatic dysfunction were given informed consent before sample collection. Description of the protocols used to collect milk samples and the diversity of subjects present in both datasets. Cohort 1 consists of 8 samples for each of the 6 subjects (48 samples total) including milk from 4 secretor mothers and 2 non-secretor mothers spanning from 6 hrs to 42 days postpartum. Sample collection was previously described^28,29^. Cohort 2 consists of 2 samples over each of the 5 (10 samples total) including samples from 4 secretor mothers and 1 non-secretor mother spanning 1 to 2 days postpartum. Sample collection was previously described^30^.

### 7.2 Illumina MRNA microarrays & Glycoprofiling

All expression and glycoprofiling measurements were sample-matched. Therefore, comparisons across data-types occurred within each individual sample described in the previous section. Not all samples in these studies have both microarray and glycoprofile measurements, only the samples described in the previous section have matched glycomics and transcriptomics data.

mRNA was isolated from TRIzol-treated milk fat in each sample. Expression in cohort 1 was measured using HumanHT-12 v4 Expression Beadchip microarrays (Illumina, Inc.) with ∼44k probes. Extraction of mRNA and measurement of expression in milk samples was performed as previously described ^28,29^. Gene expression data for cohort 1 were retrieved from the Gene Expression Omnibus at accession: GSE36936. Cohort 2 gene expression data were measured using a Human Ref-8 BeadChip array (Illumina, Inc) with ∼22k probes. Extraction of mRNA and related methods were previously described ^30^. Expression data for cohort 1 can be accessed at accession: GSE12669. Both microarrays were background corrected. The cohort 1 microarray was normalized using cubic spline normalization and the cohort 2 microarray was normalized using the robust spline normalization.

As previously described^41,91^, HMO composition and abundance data were collected using high-performance liquid chromatography (HPLC) with 2-aminobenzamide (CID: 6942) derivatization and a raffinose (CID:439242) standard. 16 HMOs were measured using retention time and commercial standards including 2-fucosyllactose (2’FL), 3-fucosyllactose (3FL), 3-sialyllactose (3’SL), lacto-N-tetraose (LNT), lacto-N-neotetraose (LNnT), lacto-N-fucopentaose (LNFP1, LNFP2 and LNFP3), sialyl-LNT (LSTb and LSTc), difucosyl-LNT (DFLNT), disialyllacto-N-tetraose (DSLNT), fucosyl-lacto-N-hexaose (FLNH), difucosyl-lacto-N-hexaose (DFLNH), fucosyl-disialyl-lacto-N-hexaose (FDSLNH) and disialyl-lacto-N-hexaose (DSLNH). Technicians were blinded to sample metadata. HMO composition and abundance measurement for cohort 1 were fully described in ^17^. Measurements for cohort 2 are previously unpublished and used the same methodology.

### 7.3 Software

Modeling of HMO biosynthesis was performed in Matlab 2016b using the CobraToolbox ^92^. All analysis of biosynthetic models, interpretation and statistics were performed in R v3.5 and v3.6. In R, we used *bigmemory, bigalgebra* and *biganalytics* to handle the millions of models and associated statistics ^93^. We used *metap* for pooling p-values^94^.

### 7.4 Generation and scoring of glycosylation network models

Here we attempt to determine the genes responsible for making HMOs through the construction and interrogation of models of their biosynthesis. Similar to the other biosynthetically constrained glycomic models like the milk metaglycome^21^, Cartoonist^95^ and several N-glycome simulations^13,96–98^, we began with a set of elementary reactions. Enumerating all feasible permutations of the elementary reaction (**Figure** 3A; S1.1.1), we delineated every possible reaction series from lactose to each of the 16 most abundant HMOs. Of the measured HMOs, 11 have fully determined molecular structures, while the remaining five have multiple candidate structures (**Figure** 1C, **Figure** S 12)^6,8,34,99–101^. The set of all possible reactions leading to characterized and ambiguous structures formed the Complete Network (**Figure** 3B; Supplemental Methods S1.1.1). Though non-lysosomal glycosidase^102–104^ reactions are not explicitly specified, they are implicitly encoded in the flux. To reduce the Complete Network to a more manageable size, we identified and removed all reactions that do not lead to observed oligosaccharides using Flux Variability Analysis (FVA; Supplemental Methods S1.2.4;^105–107^). This trimming (**Figure** 3C; Supplemental Methods S1.1.2) defines the Reduced Network (**Figure** 3D; Supplemental Methods S1.1.2). The Reduced Network describes many candidate models that can uniquely simulate the HMO abundance collected through High-Performance Liquid Chromatography (HPLC). A mixed integer linear programing (Supplemental Methods S1.2.5;^108,109^) approach was employed to extract candidate models from the Reduced Network capable of uniquely recapitulating the HPLC data with minimal reactions (**Figure** 3E; Supplemental Methods S1.1.3). The reactions of each candidate model were parameterized to determine the necessary flow of material (flux) through each reaction to reproduce the measured oligosaccharide profiles (**Figure** 3F; Supplemental Methods S1.1.3; S). The models were ranked by the consistency between the predicted flux and the expression of genes believed to be associated with each reaction (**Figure** 3G; Supplemental Methods S1.1.4). This consistency is evaluated by the Spearman correlation of changes in flux and gene expression across subjects (**Figure** 3H; Supplemental Methods S1.1.4.1).

### 7.5 Candidate Model Ranking, Model Selection and Selection Validation

Model scores, indicating the consistency between flux and gene expression (S1.1.4.1), were used to rank candidate models (S1.1.4.2). The distribution of model scores computed from each dataset were approximately normal, as evidenced by their linear Q-Q plots. This permitted the construction of a background normal distribution of model scores (**Figure** S 20). We then selected high-performing models, those with z-score normalized model scores greater than 1.646 (i.e., greater than the top 5% of scores from a normal distribution) for further study. Model selection was performed on scores computed independently for cohort 1 and cohort 2. Commonly high-performing models were those that perform well in both cohort 1 and cohort 2. Hypergeometric enrichment was used to confirm that the top cohort 1 and cohort 2 models significantly overlapped. (see Supplemental Methods S1.1.4.2)

### 7.6 Summary Network Extraction from the Reduced Network

The Summary Network relates a heuristic selection of the most important reactions in the HMO biosynthesis network as measured by proportion of inclusion in the commonly high-performing models and enrichment in the commonly high-performing models relative to the background. Paths drawn from observed HMOs to the root lactose were scored for their aggregate importance. The top 5% of paths leading to each observed HMO were retained to form the Summary Network (Supplemental Methods see S1.1.4.3).

### 7.7 Ambiguous Gene Selection

We aimed to match 10 elementary glycosyltransferase reactions to the supporting genes (**Table** 1). Candidate genes were filtered from the relevant gene families to exclude gene products well known to perform unrelated reactions (**Table** 1). Candidate genes were first evaluated for expression in breast epithelium samples including microarrays in this study, independent RNA-Seq (GSE45669) ^31^ and comparison to global expression distributions in GTEx^32^; genes unmeasured by microarray in at least 75% of microarray samples (3rd Quartile, Q3) within each cohort were excluded unless they were non-negligibly expressed in the independent RNA-Seq (TPM_Lemay_>2 or TPM_Lemay_>Median(TPM_GTEx_) (see supplemental results, **Table** S 1, **Figure** S 7).

We used the model score definition, which quantifies how well the genes explain a model, i.e., if the expression of the genes are best correlated to the normalized flux of the reaction (**Figure** S 11, S1.1.4) they are proposed to support. We examined each gene contribution to the overall model score in three ways to determine a consensus support score for each gene-reaction association (see S1.1.5.2).

The first metric we examined was the proportion (PROP) of commonly high-performing models best explained by an isoform relative to the proportion of background models that select that same isoform. The second metric was the average gene-linkage score (GLS) in high-performing models, i.e., the Spearman correlation between the normalized flux (**Figure** S 11, S1.1.4) and gene expression of corresponding candidate genes. The gene-linkage score is a continuous measure of the consistency between each gene with the flux it was proposed to support. Because it considers every gene, not just the most flux-consistent gene, it is helpful for judging performance when the most flux-consistent gene is more ambiguous. The third metric was the model-score contribution (MSC). MSC quantifies the Pearson correlation between the gene-linkage score, the gene expression consistency with the normalized flux, and the overall model score (i.e., the average correlation of all most-flux-consistent genes). The model score indicates the frequency with which a gene is the most flux-consistent gene normalized by its contribution relative to the other most flux-consistent genes in that model.

An aggregate reaction support score was constructed to describe performance within each individual score (PROP, GLS, and MSC) and consistency across cohorts. To measure significance, the gene-linkage score matrix (i.e., Spearman correlation between each candidate gene and the corresponding normalized flux for each model) was shuffled (n=27) and all analyses rerun on each shuffle to generate a permuted background distribution for PROP, GLS and MSC; shuffling of the GLS matrix was done using a perfect minimal hash to remap all entries back to the GLS matrix in a random order^110^. Performance within each independent cohort was described as the sum of z-scores for each of three measures; z-score was calculated relative to the mean and standard deviations of these scores in the permutation results. Consistency across cohorts was determined by pooling p-values using the Fisher’s log-sum method ^94,111^. The score presented in **Figure** 5B is the -log_10_(FDR(cohort-pooled-p).

### 7.8 *In vitro* glycosyltransferase activity assays

Recombinant forms of the respective glycosyltransferases were expressed and purified as previously described^112^. Enzyme activity was determined using the UDP-Glo^™^ or UMP/CMP-Glo^™^ Glycosyltransferase Assay (Promega) that determined UDP/CMP concentration formed as a by-product of the glycosyltransferase reaction. Assays were performed according to the manufacturer’s instructions using reactions (10 µL) that consisted of a universal buffer containing 100 mM each of MES, MOPS, and TRIS, pH 7.0, donor (1 mM UDP-GlcNAc (Promega) for B3GNT2; 1 mM UDP-Gal (Promega) for B3GALT2; 0.2 mM CMP-SA (Nacalai USA Inc.) for ST3GAL1-6, ST6GALNAC2, and ST6GALNAC5), 1 mM acceptor (lactose (Sigma) and lacto-N-neotetraose (LNnT) (Carbosynth) for B3GNT2; lacto-N-tetraose (LNT, Bode lab) and pentasaccharide (GlcNAc-b1,3-Gal-b1,4-GlcNAc-b1,3-Gal-b1,4-Glc, Boons lab, University of Georgia) for B3GALT2; LNnT, LNT, and Gal-β1,3-GalNAc (Carbosynth) for ST3GAL1-6; LNT for ST6GALNAC2 and ST6GALNAC5. The B3GNT2 and B3GALT2 assays also contained 1 mg/ml BSA and 5 mM MnCl_2_. Assays were carried out for 1 h (B3GNT2, B3GALT2, ST6GALNAC2, and ST6GALNAC5) or 30 min (ST3GAL1-6) at 37 °C. Reactions (5 μL) were stopped by mixing with an equal volume of Detection Reagent (5 μL) in white polystyrene, low-volume, 384-well assay plates (Corning) and incubated for 60 min at room temperature. After incubation, luminescence measurements were performed using a GloMax Multi Detection System plate reader (Promega). The average luminescence was subtracted from the average luminescence of respective blank to correct for background. Background and reaction measurements were performed in triplicate.

### 7.9 Differential expression (DE) analysis

The differential expression analysis was conducted on three different datasets: 1) 16 different HMOs (2’FL, 3’SL, 3FL, FLNH, LNT, LNnT, LSTb, LNFP-III, LNFP-II, LNFP-I, DFLNT, LSTc, DSLNT, FDSLNH, DSLNH, DFLNH), 2) 19 glycan motifs (X18, X32, X34, X35, X37, X40, X62, X63, X64, X65, X66, X94, X106, X113, X120, X127, X141, X142, X143, see **Figure** S 19), and 3) 4 differential motifs for the difference (“conversion rate”) between related motifs (X65-X40, X106-X62, X63-X37, X62-X40, see **Figure** S 19). Substructure abundance for glycan motifs and conversion ratios were computed using Glycompare v1 ^17^. The gene expression data were downloaded from the Gene Expression Omnibus ^113^ (GSE36936). Specifically, for each HMO, motif or differential motif, we used concentration (e.g., HMO– *3FL*) as the predictor for gene expression in the differential expression analysis (e.g., “gene expression ∼ [3FL]”). The differential expression analysis was performed by fitting linear models using empirical Bayes method as implemented in the *limma* v3.40.6 in R v3.6.1 package ^114^ and p-values were adjusted for multiple testing using Benjamini-Hochberg (BH) method^115^. In this way, we determined gene-expression signatures indicative of each HMO and motif abundance.

### 7.10 Ingenuity Pathways Analysis (IPA) upstream regulator

Differential expression signatures indicative of differential abundance in 16 HMOs, 19 motifs and 4 differential motifs were analyzed to predict upstream regulators using Ingenuity Pathway Analysis (IPA, QIAGEN Inc.). Gene expression signatures indicative of HMO and motif abundance were defined as genes differentially expressed with abundance in the previous *limma* analysis(FDR q<0.05 and |Fold Change|>1.5).

### 7.11 *De novo* TF binding site motifs discovery and known TF binding site identification

We downloaded promoter sequences (file: “*upstream1000*.*fa*.*gz*”; version: GRCH38) from UCSC Genome Browser public database (https://genome.ucsc.edu/) for the O-glycosyltransferase genes used in this study (**Table** S 1). These promoter sequences included 1,000 bases upstream of annotated transcription starts of RefSeq genes with annotated 5’ UTRs. To conduct *de novo* TF binding site motifs discovery, we first applied the motif discovery program MEME ^116^ to identify candidate TF binding site motifs on the downloaded promoter sequences with default parameters. The 10 TF binding site motifs found by MEME were analyzed further for matches to known TF binding sites for mammalian transcription factors in the motif databases, JASPAR Vertebrates ^117^, via motif comparison tool, TOMTOM ^118^. The resulting discovered TF binding site motifs and their significantly associated known TF binding sites (**Table** S 6, **Table** S 7) for mammalian transcription factors were used further to compare with the IPA predicted upstream regulators.

## REFERENCES

1. Edmond, K. M. et al. Delayed Breastfeeding Initiation Increases Risk of Neonatal Mortality. Pediatrics 117, e380–e386 (2006).

2. Bode, L. Human milk oligosaccharides: every baby needs a sugar mama. Glycobiology 22, 1147–1162 (2012).

3. Jantscher-Krenn, E. & Bode, L. Human milk oligosaccharides and their potential benefits for the breast-fed neonate. Minerva Pediatr. 64, 83–99 (2012).

4. Coppa, G. V. et al. Changes in Carbohydrate Composition in Human Milk Over 4 Months of Lactation. Pediatrics 91, (1993).

5. Picciano, M. F. Nutrient Composition of Human Milk. Pediatr. Clin. North Am. 48, 53–67 (2001).

6. Bode, L. The functional biology of human milk oligosaccharides. Early Hum. Dev. 91, 619–622 (2015).

7. Azad, M. B. et al. Human Milk Oligosaccharide Concentrations Are Associated with Multiple Fixed and Modifiable Maternal Characteristics, Environmental Factors, and Feeding Practices. J. Nutr. 148, 1733–1742 (2018).

8. Kobata, A. Structures and application of oligosaccharides in human milk. Proc. Jpn. Acad. Ser. B Phys. Biol. Sci. 86, 731–747 (2010).

9. Etzold, S. & Bode, L. Glycan-dependent viral infection in infants and the role of human milk oligosaccharides. Curr. Opin. Virol. 7, 101–107 (2014).

10. Zhou, R. et al. Deficiency of intestinal α1-2-fucosylation exacerbates ethanol-induced liver disease in mice. Alcohol. Clin. Exp. Res. (2020) doi: 10.1111/acer.14405.

11. Kellman, B. P. et al. A consensus-based and readable extension of Linear Code for Reaction Rules (LiCoRR). bioRxiv 2020.05.31.126623 (2020) doi: 10.1101/2020.05.31.126623.

12. Kobata, A. Possible application of milk oligosaccharides for drug development. Chang Gung Med. J. 26, 621–636 (2003).

13. Spahn, P. N. et al. A Markov chain model for N-linked protein glycosylation – towards a lowparameter tool for model-driven glycoengineering. Metabolic Engineering vol. 33 52–66 (2016).

14. Liang, C. et al. A Markov model of glycosylation elucidates isozyme specificity and glycosyltransferase interactions for glycoengineering. Current Research in Biotechnology in press, (2020).

15. Liu, G. & Neelamegham, S. A computational framework for the automated construction of glycosylation reaction networks. PLoS One 9, e100939 (2014).

16. McDonald, A. G., Tipton, K. F. & Davey, G. P. A Knowledge-Based System for Display and Prediction of O-Glycosylation Network Behaviour in Response to Enzyme Knockouts. PLoS Comput. Biol. 12, e1004844 (2016).

17. Bao, B. et al. Correcting for sparsity and non-independence in glycomic data through a systems biology framework. bioRxiv 693507 2019) doi: 10.1101/693507.

18. Akune, Y. et al. Comprehensive analysis of the N-glycan biosynthetic pathway using bioinformatics to generate UniCorn: A theoretical N-glycan structure database. Carbohydr. Res. 431, 56–63 (2016).

19. Lewis, N. E., Nagarajan, H. & Palsson, B. O. Constraining the metabolic genotype-phenotype relationship using a phylogeny of in silico methods. Nat. Rev. Microbiol. 10, 291–305 (2012).

20. Burgard, A. P., Vaidyaraman, S. & Maranas, C. D. Minimal reaction sets for Escherichia coli metabolism under different growth requirements and uptake environments. Biotechnol. Prog. 17, 791–797 (2001).

21. Agravat, S. B., Song, X., Rojsajjakul, T., Cummings, R. D. & Smith, D. F. Computational approaches to define a human milk metaglycome. Bioinformatics 32, 1471–1478 (2016).

22. Spahn, P. N., Hansen, A. H., Kol, S., Voldborg, B. G. & Lewis, N. E. Predictive glycoengineering of biosimilars using a Markov chain glycosylation model. Biotechnol. J. 12, (2017).

23. Nishihara, S. et al. Molecular genetic analysis of the human Lewis histo-blood group system. J. Biol. Chem. 269, 29271–29278 (1994).

24. Kudo, T. et al. Molecular genetic analysis of the human Lewis histo-blood group system. II. Secretor gene inactivation by a novel single missense mutation A385T in Japanese nonsecretor individuals. J. Biol. Chem. 271, 9830–9837 (1996).

25. Koda, Y., Soejima, M., Liu, Y. & Kimura, H. Molecular basis for secretor type alpha(1,2)-fucosyltransferase gene deficiency in a Japanese population: a fusion gene generated by unequal crossover responsible for the enzyme deficiency. Am. J. Hum. Genet. 59, 343–350 (1996).

26. Thurl, S., Henker, J., Siegel, M., Tovar, K. & Sawatzki, G. Detection of four human milk groups with respect to Lewis blood group dependent oligosaccharides. Glycoconj. J. 14, 795–799 (1997).

27. Stahl, B. et al. Detection of four human milk groups with respect to Lewis-blood-group-dependent oligosaccharides by serologic and chromatographic analysis. Adv. Exp. Med. Biol. 501, 299–306 (2001).

28. Mohammad, M. A., Hadsell, D. L. & Haymond, M. W. Gene regulation of UDP-galactose synthesis and transport: potential rate-limiting processes in initiation of milk production in humans. Am. J. Physiol. Endocrinol. Metab. 303, E365–76 (2012).

29. Mohammad, M. A. & Haymond, M. W. Regulation of lipid synthesis genes and milk fat production in human mammary epithelial cells during secretory activation. Am. J. Physiol. Endocrinol. Metab. 305, E700–16 (2013).

30. Maningat, P. D. et al. Gene expression in the human mammary epithelium during lactation: the milk fat globule transcriptome. Physiol. Genomics 37, 12–22 (2009).

31. Lemay, D. G. et al. RNA sequencing of the human milk fat layer transcriptome reveals distinct gene expression profiles at three stages of lactation. PLoS One 8, e67531 (2013).

32. Carithers, L. J. et al. A novel approach to high-quality postmortem tissue procurement: the GTEx project. Biopreserv. Biobank. 13, 311–319 (2015).

33. Blank, D., Dotz, V., Geyer, R. & Kunz, C. Human milk oligosaccharides and Lewis blood group: individual high-throughput sample profiling to enhance conclusions from functional studies. Adv. Nutr. 3, 440S–9S (2012).

34. Wu, S., Tao, N., German, J. B., Grimm, R. & Lebrilla, C. B. Development of an annotated library of neutral human milk oligosaccharides. J. Proteome Res. 9, 4138–4151 (2010).

35. Brodbeck, U. & Ebner, K. E. Resolution of a soluble lactose synthetase into two protein components and solubilization of microsomal lactose synthetase. J. Biol. Chem. 241, 762–764 (1966).

36. Nakhasi, H. L. & Quasba, P. K. Quantitation of milk proteins and their mRNAs in rat mammary gland at various stages of gestation and lactation. J. Biol. Chem. 254, 6016–6025 (1979).

37. Morrow, A. L. et al. Fucosyltransferase 2 non-secretor and low secretor status predicts severe outcomes in premature infants. J. Pediatr. 158, 745–751 (2011).

38. Autran, C. A. et al. Human milk oligosaccharide composition predicts risk of necrotising enterocolitis in preterm infants. Gut 67, 1064–1070 (2018).

39. Morrow, A. L. et al. Human Milk Oligosaccharide Blood Group Epitopes and Innate Immune Protection against Campylobacter and Calicivirus Diarrhea in Breastfed Infants. in 443–446 (Springer, Boston, MA, 2004).

40. Yu, Z.-T., Nanda Nanthakumar, N. & Newburg, D. S. The Human Milk Oligosaccharide 2′-Fucosyllactose Quenches Campylobacter jejuni–Induced Inflammation in Human Epithelial Cells HEp-2 and HT-29 and in Mouse Intestinal Mucosa. The Journal of Nutrition vol. 146 1980–1990 (2016).

41. Alderete, T. L. et al. Associations between human milk oligosaccharides and infant body composition in the first 6 mo of life. Am. J. Clin. Nutr. 102, 1381–1388 (2015).

42. Uwaezuoke, S. N., Eneh, C. I. & Ndu, I. K. Relationship Between Exclusive Breastfeeding and Lower Risk of Childhood Obesity: A Narrative Review of Published Evidence. Clin. Med. Insights Pediatr. 11, 1179556517690196 (2017).

43. Uwaezuoke, S. et al. Maternal diet during exclusive breastfeeding can predict food preference in preschoolers: A cross-sectional study of mother-child dyads in Enugu, south-east Nigeria. Int. J. Child Health Nutr. 6, 70–79 (2017).

44. Moro, G. et al. Dosage-related bifidogenic effects of galacto-and fructooligosaccharides in formula-fed term infants. J. Pediatr. Gastroenterol. Nutr. 34, 291–295 (2002).

45. Costalos, C., Kapiki, A., Apostolou, M. & Papathoma, E. The effect of a prebiotic supplemented formula on growth and stool microbiology of term infants. Early Hum. Dev. 84, 45–49 (2008).

46. Vos, A. P. et al. A specific prebiotic oligosaccharide mixture stimulates delayed-type hypersensitivity in a murine influenza vaccination model. Int. Immunopharmacol. 6, 1277–1286 (2006).

47. Viverge, D., Grimmonprez, L., Cassanas, G., Bardet, L. & Solere, M. Discriminant carbohydrate components of human milk according to donor secretor types. J. Pediatr. Gastroenterol. Nutr. 11, 365–370 (1990).

48. McGuire, M. K. et al. What’s normal? Oligosaccharide concentrations and profiles in milk produced by healthy women vary geographically. Am. J. Clin. Nutr. 105, 1086–1100 (2017).

49. Furuike, T., Yamada, K., Ohta, T., Monde, K. & Nishimura, S.-I. An efficient synthesis of a biantennary sialooligosaccharide analog using a 1,6-anhydro-β-lactose derivative as a key synthetic block. Tetrahedron 59, 5105–5113 (2003).

50. Fair, R. J., Hahm, H. S. & Seeberger, P. H. Combination of automated solid-phase and enzymatic oligosaccharide synthesis provides access to α(2,3)-sialylated glycans. Chem. Commun. 51, 6183–6185 (2015).

51. Yao, W., Yan, J., Chen, X., Wang, F. & Cao, H. Chemoenzymatic synthesis of lacto-N-tetrasaccharide and sialyl lacto-N-tetrasaccharides. Carbohydr. Res. 401, 5–10 (2015).

52. Prudden, A. R. et al. Synthesis of asymmetrical multiantennary human milk oligosaccharides. Proc. Natl. Acad. Sci. U. S. A. 114, 6954–6959 (2017).

53. Prudden, A. R., Chinoy, Z. S., Wolfert, M. A. & Boons, G.-J. A multifunctional anomeric linker for the chemoenzymatic synthesis of complex oligosaccharides. Chem. Commun. 50, 7132–7135 (2014).

54. Bode, L. et al. Overcoming the limited availability of human milk oligosaccharides: challenges and opportunities for research and application. Nutr. Rev. 74, 635–644 (2016).

55. Guan, N. & Chen, R. Recent Technology Development for the Biosynthesis of Human Milk Oligosaccharide. Recent patents on biotechnology (2018).

56. Lee, W.-H. et al. Whole cell biosynthesis of a functional oligosaccharide, 2\textasciiacutex-fucosyllactose, using engineered Escherichia coli. Microb. Cell Fact. 11, 48 (2012).

57. Chin, Y.-W., Kim, J.-Y., Lee, W.-H. & Seo, J.-H. Enhanced production of 2’-fucosyllactose in engineered Escherichia coli BL21star(DE3) by modulation of lactose metabolism and fucosyltransferase. J. Biotechnol. 210, 107–115 (2015).

58. Baumgärtner, F., Seitz, L., Sprenger, G. A. & Albermann, C. Construction of Escherichia coli strains with chromosomally integrated expression cassettes for the synthesis of 2\textasciiacutex-fucosyllactose. Microb. Cell Fact. 12, 40 (2013).

59. Baumgärtner, F., Conrad, J., Sprenger, G. A. & Albermann, C. Synthesis of the Human Milk Oligosaccharide Lacto-N -Tetraose in Metabolically Engineered, Plasmid-Free E. coli. Chembiochem 15, 1896–1900 (2014).

60. Kumazaki, T. & Yoshida, A. Biochemical evidence that secretor gene, Se, is a structural gene encoding a specific fucosyltransferase. Proc. Natl. Acad. Sci. U. S. A. 81, 4193–4197 (1984).

61. Mollicone, R. et al. Activity, Splice Variants, Conserved Peptide Motifs, and Phylogeny of Two New α1,3-Fucosyltransferase Families (FUT10 and FUT11). J. Biol. Chem. 284, 4723–4738 (2009).

62. Kaneko, M. et al. Assignment1 of the human α 1,3-fucosyltransferase IX gene (FUT9) to chromosome band 6q16 by in situ hybridization. Cytogenetic and Genome Research vol. 86 329–330 (1999).

63. Nishihara, S. et al. α1, 3-Fucosyltransferase 9 (FUT9; Fuc-TIX) preferentially fucosylates the distal GlcNAc residue of polylactosamine chain while the other four α1, 3FUT members preferentially fucosylate the inner GlcNAc residue. FEBS Lett. 462, 289–294 (1999).

64. Niemelä, R. et al. Complementary acceptor and site specificities of Fuc-TIV and Fuc-TVII allow effective biosynthesis of sialyl-TriLex and related polylactosamines present on glycoprotein counterreceptors of selectins. J. Biol. Chem. 273, 4021–4026 (1998).

65. Mondal, N. et al. Distinct human α(1,3)-fucosyltransferases drive Lewis-X/sialyl Lewis-X assembly in human cells Downloaded from. (2018) doi: 10.1074/jbc.RA117.000775.

66. Kurosawa, N., Inoue, M., Yoshida, Y. & Tsuji, S. Molecular Cloning and Genomic Analysis of Mouse Galβ1,3GalNAc-specific GalNAc α2,6-Sialyltransferase. Journal of Biological Chemistry vol. 271 15109–15116 (1996).

67. Kurosawa, N., Kojima, N., Inoue, M., Hamamoto, T. & Tsuji, S. Cloning and expression of Gal beta 1,3GalNAc-specific GalNAc alpha 2,6-sialyltransferase. J. Biol. Chem. 269, 19048–19053 (1994).

68. Okajima, T. et al. Molecular Cloning of Brain-specific GD1α Synthase (ST6GalNAc V) Containing CAG/Glutamine Repeats. J. Biol. Chem. 274, 30557–30562 (1999).

69. Okajima, T. et al. Expression cloning of human globoside synthase cDNAs. Identification of beta 3Gal-T3 as UDP-N-acetylgalactosamine:globotriaosylceramide beta 1,3-N-acetylgalactosaminyltransferase. J. Biol. Chem. 275, 40498–40503 (2000).

70. Sjoberg, E. R., Kitagawa, H., Glushka, J., van Halbeek, H. & Paulson, J. C. Molecular Cloning of a Developmentally Regulated N-Acetylgalactosamine 2,6-Sialyltransferase Specific for Sialylated Glycoconjugates. J. Biol. Chem. 271, 7450–7459 (1996).

71. Tsuchida, A. et al. Molecular cloning and expression of human ST6GalNAc III: restricted tissue distribution and substrate specificity. J. Biochem. 138, 237–243 (2005).

72. Lee, Y.-C. et al. Molecular Cloning and Functional Expression of Two Members of Mouse NeuAcα2,3Galβ1,3GalNAc GalNAcα2,6-Sialyltransferase Family, ST6GalNAc III and IV. Journal of Biological Chemistry vol. 274 11958–11967 (1999).

73. Tsuchida, A. et al. Synthesis of Disialyl Lewis a (Lea) Structure in Colon Cancer Cell Lines by a Sialyltransferase, ST6GalNAc VI, Responsible for the Synthesis of α-Series Gangliosides. J. Biol. Chem. 278, 22787–22794 (2003).

74. Kitagawa, H. & Paulson, J. C. Cloning of a novel alpha 2,3-sialyltransferase that sialylates glycoprotein and glycolipid carbohydrate groups. J. Biol. Chem. 269, 1394–1401 (1994).

75. Kono, M. et al. Mouse beta-galactoside alpha 2,3-sialyltransferases: comparison of in vitro substrate specificities and tissue specific expression. Glycobiology 7, 469–479 (1997).

76. Blixt, O. et al. Glycan microarrays for screening sialyltransferase specificities. Glycoconj. J. 25, 59–68 (2008).

77. Weinstein, J., de Souza-e-Silva, U. & Paulson, J. C. Sialylation of glycoprotein oligosaccharides N-linked to asparagine. Enzymatic characterization of a Gal beta 1 to 3(4)GlcNAc alpha 2 to 3 sialyltransferase and a Gal beta 1 to 4GlcNAc alpha 2 to 6 sialyltransferase from rat liver. J. Biol. Chem. 257, 13845–13853 (1982).

78. Gillespie, W., Kelm, S. & Paulson, J. C. Cloning and expression of the Gal beta 1, 3GalNAc alpha 2,3-sialyltransferase. J. Biol. Chem. 267, 21004–21010 (1992).

79. Miyazaki, H. et al. Expression Cloning of Rat cDNA Encoding UDP-galactose:GD2 β1,3-galactosyltransferase That Determines the Expression of GD1b/GM1/GA1. J. Biol. Chem. 272, 24794–24799 (1997).

80. Amado, M. et al. A family of human beta3-galactosyltransferases. Characterization of four members of a UDP-galactose:beta-N-acetyl-glucosamine/beta-nacetyl-galactosamine beta-1,3-galactosyltransferase family. J. Biol. Chem. 273, 12770–12778 (1998).

81. Isshiki, S. et al. Cloning, Expression, and Characterization of a Novel UDP-galactose:β-N-Acetylglucosamine β1,3-Galactosyltransferase (β3Gal-T5) Responsible for Synthesis of Type 1 Chain in Colorectal and Pancreatic Epithelia and Tumor Cells Derived Therefrom. Journal of Biological Chemistry vol. 274 12499–12507 (1999).

82. Schwientek, T. et al. Cloning of a Novel Member of the UDP-Galactose:β-N-Acetylglucosamine β1,4-Galactosyltransferase Family, β4Gal-T4, Involved in Glycosphingolipid Biosynthesis. J. Biol. Chem. 273, 29331–29340 (1998).

83. Sato, T., Aoki, N., Matsuda, T. & Furukawa, K. Differential effect of alpha-lactalbumin on beta-1,4-galactosyltransferase IV activities. Biochem. Biophys. Res. Commun. 244, 637–641 (1998).

84. Shiraishi, N. et al. Identification and Characterization of Three Novel β1,3-N-Acetylglucosaminyltransferases Structurally Related to the β1,3-Galactosyltransferase Family. J. Biol. Chem. 276, 3498–3507 (2001).

85. Chen, G. Y., Kurosawa, N. & Muramatsu, T. A novel variant form of murine beta-1, 6-N-acetylglucosaminyltransferase forming branches in poly-N-acetyllactosamines. Glycobiology 10, 1001–1011 (2000).

86. Bierhuizen, M. F. & Fukuda, M. Expression cloning of a cDNA encoding UDP-GlcNAc: Gal beta 1-3-GalNAc-R (GlcNAc to GalNAc) beta 1-6GlcNAc transferase by gene transfer into CHO cells expressing polyoma large tumor antigen. Proceedings of the National Academy of Sciences 89, 9326–9330 (1992).

87. Schwientek, T. et al. Control of O-glycan branch formation. Molecular cloning of human cDNA encoding a novel beta1,6-N-acetylglucosaminyltransferase forming core 2 and core 4. J. Biol. Chem. 274, 4504–4512 (1999).

88. Yeh, J. C., Ong, E. & Fukuda, M. Molecular cloning and expression of a novel beta-1, 6-N-acetylglucosaminyltransferase that forms core 2, core 4, and I branches. J. Biol. Chem. 274, 3215–3221 (1999).

89. Schwientek, T. et al. Control of O-Glycan Branch Formation: MOLECULAR CLONING OF HUMAN cDNA ENCODING A NOVEL β1,6-N-ACETYLGLUCOSAMINYLTRANSFERASE FORMING CORE 2 AND CORE 4. J. Biol. Chem. 274, 4504–4512 (1999).

90. Ujita, M., Misra, A. K., McAuliffe, J., Hindsgaul, O. & Fukuda, M. Poly-N-acetyllactosamine Extension inN-Glycans and Core 2-and Core 4-branchedO-Glycans Is Differentially Controlled by i-Extension Enzyme and Different Members of the β1, 4-Galactosyltransferase Gene Family. J. Biol. Chem. 275, 15868–15875 (2000).

91. Bode, L. et al. Human milk oligosaccharide concentration and risk of postnatal transmission of HIV through breastfeeding. Am. J. Clin. Nutr. 96, 831–839 (2012).

92. Heirendt, L. et al. Creation and analysis of biochemical constraint-based models using the COBRA Toolbox v.3.0. Nat. Protoc. 14, 639–702 (2019).

93. Kane, M. J., Emerson, J. W., Haverty, P. & Others. bigmemory: Manage massive matrices with shared memory and memory-mapped files. R package version 4, (2010).

94. Dewey, M. metap: Meta-analysis of significance values. R package version 0.7. (2016).

95. Goldberg, D., Sutton-Smith, M., Paulson, J. & Dell, A. Automatic annotation of matrix-assisted laser desorption/ionization N-glycan spectra. Proteomics 5, 865–875 (2005).

96. Hossler, P., Mulukutla, B. C. & Hu, W.-S. Systems analysis of N-glycan processing in mammalian cells. PLoS One 2, e713 (2007).

97. Krambeck, F. J. et al. A mathematical model to derive N-glycan structures and cellular enzyme activities from mass spectrometric data. Glycobiology 19, 1163–1175 (2009).

98. McDonald, A. G. et al. Galactosyltransferase 4 is a major control point for glycan branching in N-linked glycosylation. J. Cell Sci. 127, 5014–5026 (2014).

99. Mantovani, V., Galeotti, F., Maccari, F. & Volpi, N. Recent advances on separation and characterization of human milk oligosaccharides. Electrophoresis 37, 1514–1524 (2016).

100. Ninonuevo, M. R. et al. A strategy for annotating the human milk glycome. J. Agric. Food Chem. 54, 7471–7480 (2006).

101. Wu, S., Grimm, R., German, J. B. & Lebrilla, C. B. Annotation and structural analysis of sialylated human milk oligosaccharides. J. Proteome Res. 10, 856–868 (2011).

102. Wiederschain, G. Y. & Newburg, D. S. Glycoconjugate stability in human milk: glycosidase activities and sugar release. J. Nutr. Biochem. 12, 559–564 (2001).

103. Miura, K., Hakamata, W., Tanaka, A., Hirano, T. & Nishio, T. Discovery of human Golgi β-galactosidase with no identified glycosidase using a QMC substrate design platform for exoglycosidase. Bioorg. Med. Chem. 24, 1369–1375 (2016).

104. Dudzik, D. et al. Activity of N-acetyl-β-D-hexosaminidase (HEX) and its isoenzymes A and B in human milk during the first 3 months of breastfeeding. Adv. Med. Sci. 53, (2008).

105. Orth, J. D., Thiele, I. & Palsson, B. Ø. What is flux balance analysis? Nat. Biotechnol. 28, 245–248 (2010).

106. Mahadevan, R. & Schilling, C. H. The effects of alternate optimal solutions in constraint-based genome-scale metabolic models. Metab. Eng. 5, 264–276 (2003).

107. Gudmundsson, S. & Thiele, I. Computationally efficient flux variability analysis. BMC Bioinformatics 11, 489 (2010).

108. Reed, J. L. & Palsson, B. Ø. Genome-scale in silico models of E. coli have multiple equivalent phenotypic states: assessment of correlated reaction subsets that comprise network states. Genome Res. 14, 1797–1805 (2004).

109. Lee, S., Phalakornkule, C., Domach, M. M. & Grossmann, I. E. Recursive MILP model for finding all the alternate optima in LP models for metabolic networks. Comput. Chem. Eng. 24, 711–716 (2000).

110. Fredman, M. L., Komlós, J. & Szemerédi, E. Storing a Sparse Table with 0 (1) Worst Case Access Time. Journal of the ACM (JACM) vol. 31 538–544 (1984).

111. E., W. P. & W., P. E. Statistical Methods for Research Workers. By Fisher R. A.. [Pp. 239 ix VI Tables. Edinburgh and London: Oliver & Boyd. 1925. Price 15s.]The Fundamentals of Statistics. By Thurstone L. L.. [Pp. 237 xvi. New York: The Macmillan Company. 1925. Price 8s. 6d.]. Journal of the Institute of Actuaries vol. 56 326–327 (1925).

112. Moremen, K. W. et al. Expression system for structural and functional studies of human glycosylation enzymes. Nat. Chem. Biol. 14, 156–162 (2018).

113. Edgar, R., Domrachev, M. & Lash, A. E. Gene Expression Omnibus: NCBI gene expression and hybridization array data repository. Nucleic Acids Res. 30, 207–210 (2002).

114. Smyth, G. K., Thorne, N. P. & Wettenhall, J. LIMMA: Linear Models for Microarray Data Version 1.6. 6. User’s Guide (2004).

115. Benjamini, Y. & Hochberg, Y. Controlling the False Discovery Rate: A Practical and Powerful Approach to Multiple Testing. J. R. Stat. Soc. Series B Stat. Methodol. 57, 289–300 (1995).

116. Bailey, T. L. & Elkan, C. The value of prior knowledge in discovering motifs with MEME. Proc. Int. Conf. Intell. Syst. Mol. Biol. 3, 21–29 (1995).

117. Fornes, O. et al. JASPAR 2020: update of the open-access database of transcription factor binding profiles. Nucleic Acids Res. 48, D87–D92 (2020).

118. Gupta, S., Stamatoyannopoulos, J. A., Bailey, T. L. & Noble, W. S. Quantifying similarity between motifs. Genome Biol. 8, R24 (2007).

119. Taniguchi, N., Honke, K. & Fukuda, M. Handbook of Glycosyltransferases and Related Genes. (Springer Science & Business Media, 2011).

120. Narimatsu, H. Construction of a human glycogene library and comprehensive functional analysis. Glycoconj. J. 21, 17–24 (2004).

121. Schomburg, I. et al. The BRENDA enzyme information system-From a database to an expert system. J. Biotechnol. 261, 194–206 (2017).

122. Magrane, M. & UniProt Consortium. UniProt Knowledgebase: a hub of integrated protein data. Database 2011, bar009 (2011).

123. Caspi, R. et al. The MetaCyc database of metabolic pathways and enzymes and the BioCyc collection of Pathway/Genome Databases. Nucleic Acids Res. 42, D459–71 (2014).

124. Kanehisa, M., Furumichi, M., Tanabe, M., Sato, Y. & Morishima, K. KEGG: new perspectives on genomes, pathways, diseases and drugs. Nucleic Acids Res. 45, D353–D361 (2017).

125. Praissman, J. L. et al. B4GAT1 is the priming enzyme for the LARGE-dependent functional glycosylation of α-dystroglycan. Elife 3, (2014).

126. Willer, T. et al. The glucuronyltransferase B4GAT1 is required for initiation of LARGE-mediated α-dystroglycan functional glycosylation. Elife 3, (2014).

127. Yanagidani, S. et al. Purification and cDNA Cloning of GDP-L-Fuc:N-Acetyl-β-D-Glucosaminide:α1-6 Fucosyltransferase (α1-6 FucT) from Human Gastric Cancer MKN45 Cells. J. Biochem. 121, 626–632 (1997).

128. Uozumi, N. et al. Purification and cDNA cloning of porcine brain GDP-L-Fuc: N-acetyl-β-D-glucosaminide α1→ 6fucosyltransferase. J. Biol. Chem. 271, 27810–27817 (1996).

129. Kataoka, K. & Huh, N.-H. A novel beta1,3-N-acetylglucosaminyltransferase involved in invasion of cancer cells as assayed in vitro. Biochem. Biophys. Res. Commun. 294, 843–848 (2002).

130. Kitayama, K., Hayashida, Y., Nishida, K. & Akama, T. O. Enzymes responsible for synthesis of corneal keratan sulfate glycosaminoglycans. J. Biol. Chem. 282, 30085–30096 (2007).

131. Seko, A. & Yamashita, K. β1,3-N-Acetylglucosaminyltransferase-7 (β3Gn-T7) acts efficiently on keratan sulfate-related glycans. FEBS Lett. 556, 216–220 (2004).

132. Bai, X. et al. Biosynthesis of the linkage region of Glycosaminoglycans cloning and activity of galactosyltransferase ii, the sixth member of the β1, 3-galactosyltransferase family (β3GalT6). J. Biol. Chem. 276, 48189–48195 (2001).

133. Ju, T., Brewer, K., D’Souza, A., Cummings, R. D. & Canfield, W. M. Cloning and Expression of Human Core 1 β1,3-Galactosyltransferase. J. Biol. Chem. 277, 178–186 (2002).

134. Almeida, R. et al. Cloning and expression of a proteoglycan UDP-galactose: β-Xylose β1, 4-galactosyltransferase IA seventh member of the human β4-galactosyltransferase gene family. J. Biol. Chem. 274, 26165–26171 (1999).

135. Okajima, T., Yoshida, K., Kondo, T. & Furukawa, K. Human homolog of Caenorhabditis elegans sqv-3 gene is galactosyltransferase I involved in the biosynthesis of the glycosaminoglycanprotein linkage region of proteoglycans. J. Biol. Chem. 274, 22915–22918 (1999).

136. Inshaw, J. R. J., Cutler, A. J., Burren, O. S., Stefana, M. I. & Todd, J. A. Approaches and advances in the genetic causes of autoimmune disease and their implications. Nat. Immunol. 19, 674–684 (2018).

137. Rouillard, A. D. et al. The harmonizome: a collection of processed datasets gathered to serve and mine knowledge about genes and proteins. Database 2016, (2016).

138. Hu, Z.-Z., Zhuang, L., Meng, J. & Dufau, M. L. Transcriptional Regulation of the Generic Promoter III of the Rat Prolactin Receptor Gene by C/EBPβ and Sp1. J. Biol. Chem. 273, 26225–26235 (1998).

139. Sugiyama, A., Fukushima, N. & Sato, T. Transcriptional Mechanism of the β4-Galactosyltransferase 4 Gene in SW480 Human Colon Cancer Cell Line. Biol. Pharm. Bull. 40, 733–737 (2017).

140. Sato, T. & Furukawa, K. Transcriptional Regulation of the Human β-1,4-Galactosyltransferase V Gene in Cancer Cells: ESSENTIAL ROLE OF TRANSCRIPTION FACTOR Sp1. J. Biol. Chem. 279, 39574–39583 (2004).

141. Sato, T. & Furukawa, K. Sequential Action of Ets-1 and Sp1 in the Activation of the Human β-1,4-Galactosyltransferase V Gene Involved in Abnormal Glycosylation Characteristic of Cancer Cells. J. Biol. Chem. 282, 27702–27712 (2007).

142. Zhou, L., Jiang, J. & Gu, J. β1,4-Galactosyltransferase: Regulation and Signaling in Cancers. Glycoscience: Biology and Medicine 1141–1148 (2015) doi: 10.1007/978-4-431-54841-6_74.

143. Kurcon, T. et al. miRNA proxy approach reveals hidden functions of glycosylation. Proc. Natl. Acad. Sci. U. S. A. 112, 7327–7332 (2015).

144. Liu, B. et al. MiR-29b/Sp1/FUT4 axis modulates the malignancy of leukemia stem cells by regulating fucosylation via Wnt/β-catenin pathway in acute myeloid leukemia. J. Exp. Clin. Cancer Res. 38, 200 (2019).

145. Dhordain, P., Dewitte, F., Desbiens, X., Stehelin, D. & Duterque-Coquillaud, M. Mesodermal expression of the chicken erg gene associated with precartilaginous condensation and cartilage differentiation. Mech. Dev. 50, 17–28 (1995).

146. Taniguchi, A., Itaru, Y. & Matsumoto, K. Genomic structure and transcriptional regulation of human Galβ1,3GalNAc α2,3-sialyltransferase (hST3Gal I) gene. Glycobiology 11, 241–247 (2001).

147. Vandenplas, Y. et al. Human Milk Oligosaccharides: 2’-Fucosyllactose (2’-FL) and Lacto-N-Neotetraose (LNnT) in Infant Formula. Nutrients 10, (2018).

