## Supplemental Text for "Elucidating Human Milk Oligosaccharide biosynthetic genes through network-based multiomics integration"

### 1 SUPPLEMENT

#### 1.1 GLOSSARY

| Term | Definition | Full Definition |
|---|---|---|
| Reaction rules, elementary or fundamental reactions | The generic reaction types the can occur in the model | Section 4.1 |
| Candidate structures, isomers | Alternative and proposed structure for HMOs that are not yet fully characterized | **Figure** S 12 |
| Candidate gene, isoform | All genes in a glycosyltransferase family that could reasonably perform the corresponding reaction. | **Table 1** |
| Complete Network | The collection of all reactions into a network that will produce all possible glycans up to a pre-specified size | Section 4.1 |
| Reduced Network | A trimming, via FVA (Section 5.4), of to Complete Network to remove reactions unnecessary to reach observed glycans | Section 4.2 |
| Candidate Models | Many subnetworks of the Reduced Network, via MILP (Section 5.5), where each subnetwork is capable of uniquely reproducing the observed data | Section 4.3 |
| Flux & Normalized flux | Relativistic measure of the stoichiometrically balanced movement of material through a metabolic model. Flux can be normalized relative to an upstream reaction to consider substrate limitations. Flux is calculated using Flux balance analysis. | **Figure** S 11 |
| Gene-Linkage Score (GLS) | The highest Spearman correlation between the total normalized flux of through a reaction type and the expression of all corresponding candidate genes. The selection of the "best" genes to describe a linkage within a model. | Section 4.5.2 |
| Model Score | The average gene-linkage score across all 10 fundamental reaction types for a model | Section 4.4.1 |
| High-performing models | Top 5% of candidate models ranked by model score relative to a normal distribution. | Section 4.4.2 |
| Commonly high-performing models | High-performing models using data from both cohorts | Section 4.4.2 |

| | | |
|---|---|---|
| Summary Network | A visualization of the most important pathways though the Reduced Network; those most frequently and substantially used in commonly high-performing models | Section 4.4.3 |
| Proportion (PROP) | The hypergeometric enrichment of models where a specific gene was best (highest gene-linkage score within the linkage) in the set of commonly high-performing models vs the background of all models. | Section 4.5.2 |
| Model Contribution Score (MSC) | The Pearson correlation between model score and the gene-linkage score for a particular gene. | Section 4.5.2 |
| Reaction Support Score | The degree to which a gene is likely to “support” a fundamental reaction based on the aggregation of PROP, GLS, and MSC across independent datasets. | Section 4.5.2 |

### 2 Supplementary Figures

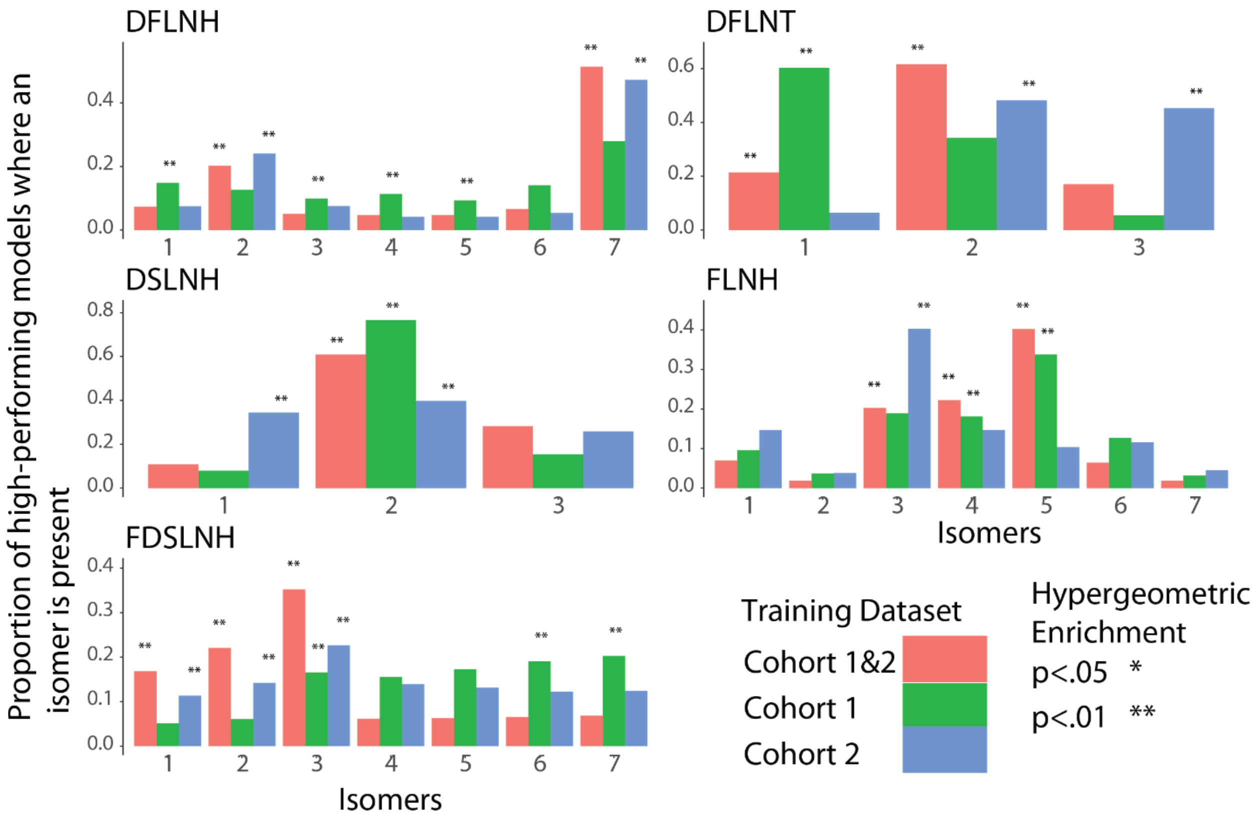

***Figure** S 1 - **Proportion of Each Structure Isoform Appearing in Top-performing Models.** Barplots describing the proportion of models in top-performing model sets containing each isoform of an ambiguous HMO structure. Top-performing model sets include those top-performing when parameterized on cohort 1 data (red), models that performed well with data from cohort 2 data (green), and models that performed well in both cohort 1 and cohort 2 (blue). Specific structures are illustrated in **Figure** S 12.*

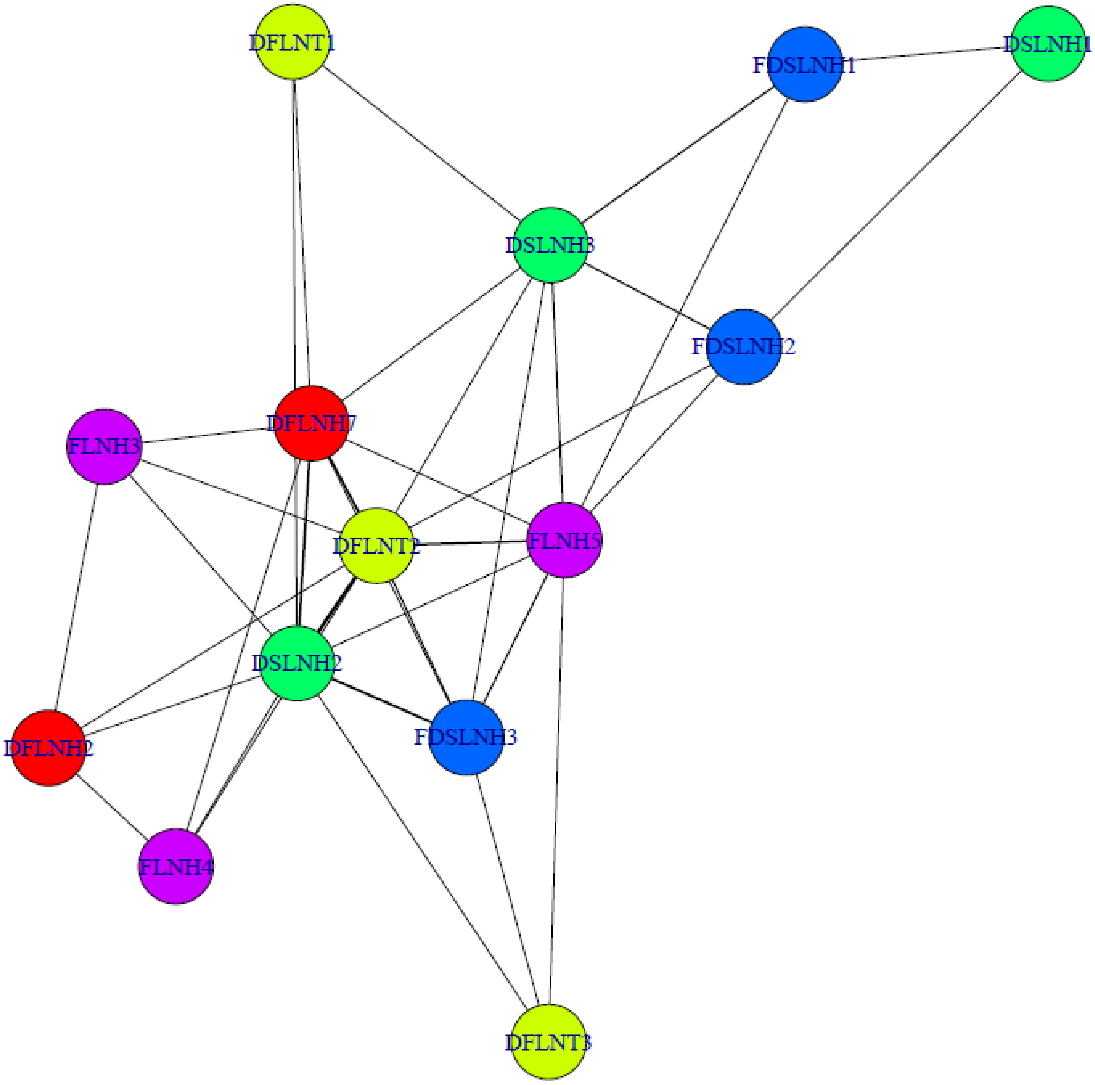

***Figure** S 2 - **Jaccard Structure Co-occurrence.** Network of the Jaccard index measuring co-occurrence of each pair of ambiguous HMOs structure within the set of best performing models. For exact numbers, see **Table** S 2.*

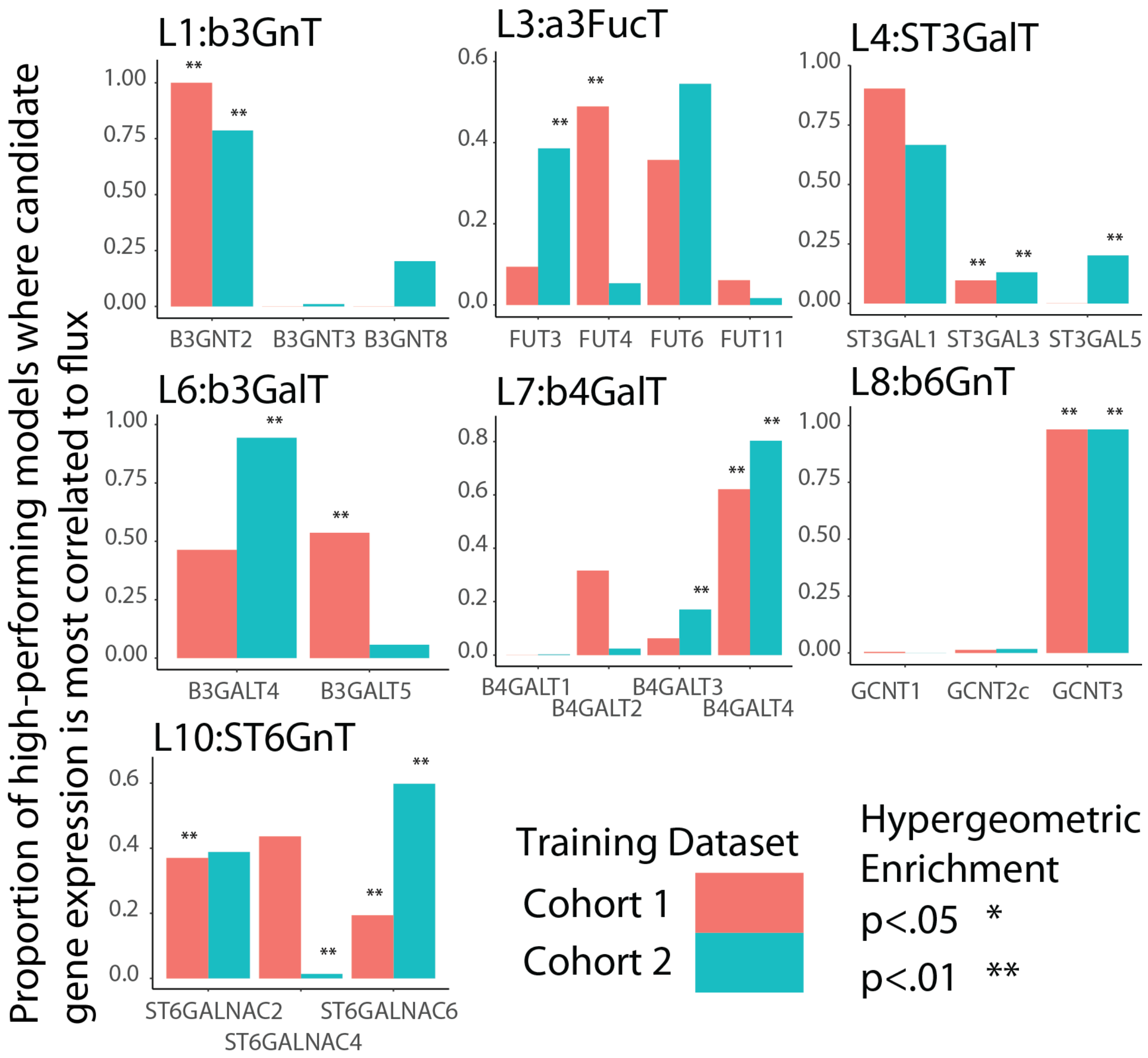

***Figure** S 3 **- Proportion of Models Where Expression of a Given Gene is Most Consistent with Predicted Flux.*** *Gene Expression was measured in dataset 1 (red) or dataset 2 (blue) then compared to normalized flux predicted for models commonly high-performing with cohort1 and cohort2 data as described in Methods section Ambiguous Gene Selection and supplemental methods section 1.1.5)*

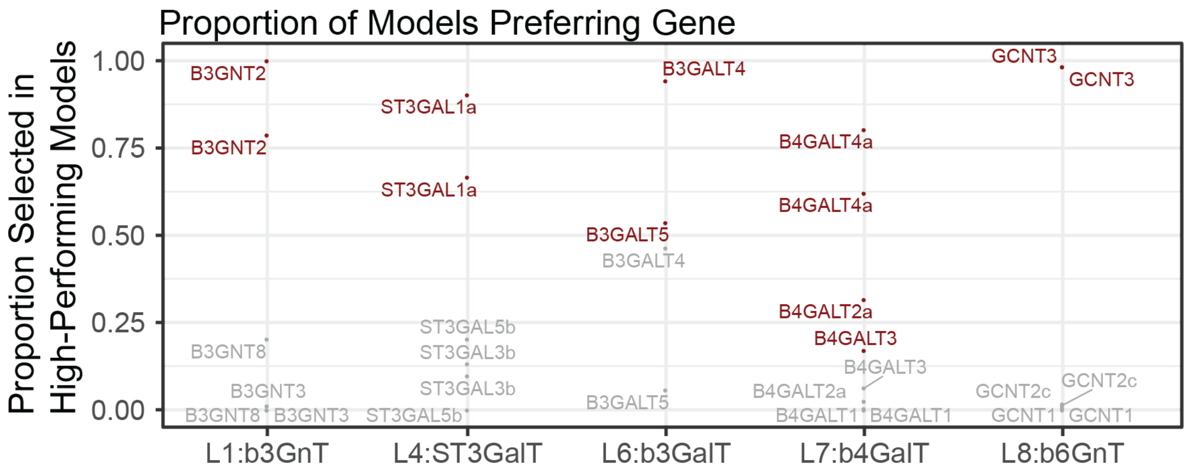

***Figure** S 4* ***– Gene expression correlation with model flux predicts enzymes involved in HMO biosynthesis.*** *The proportion of commonly high-performing models where each isoform was most correlated to the model predicted flux. Significance (p<0.001, red) is calculated by hypergeometric enrichment of representation in high-performing models compared to all models. Each gene appears twice since data are analyzed separately for the two cohorts.*

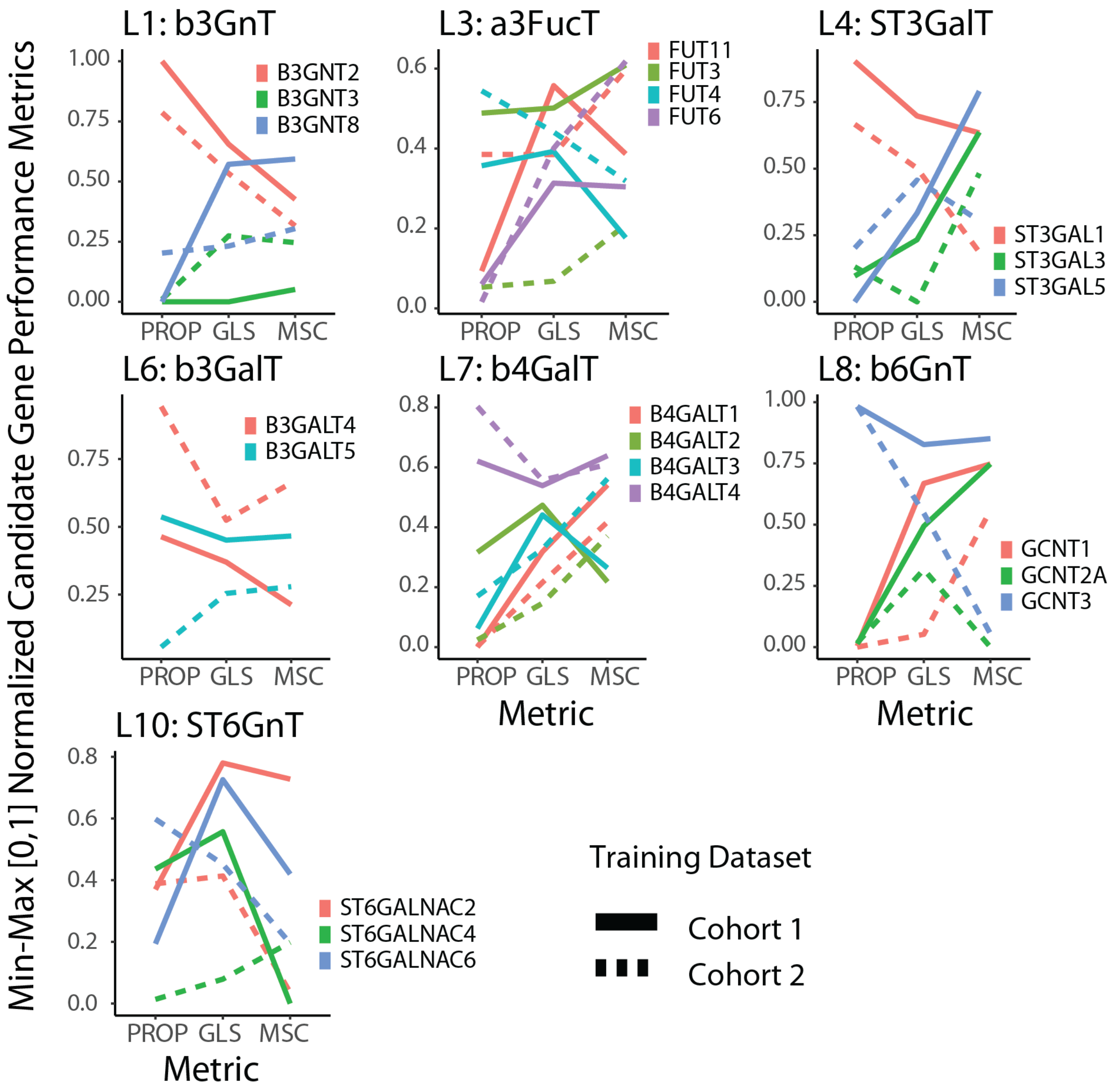

***Figure S 5 - Proportion (Prop), Gene Linkage Score (GLS) and Model Score Contribution (MSC) for each candidate gene within each gene linkage.***

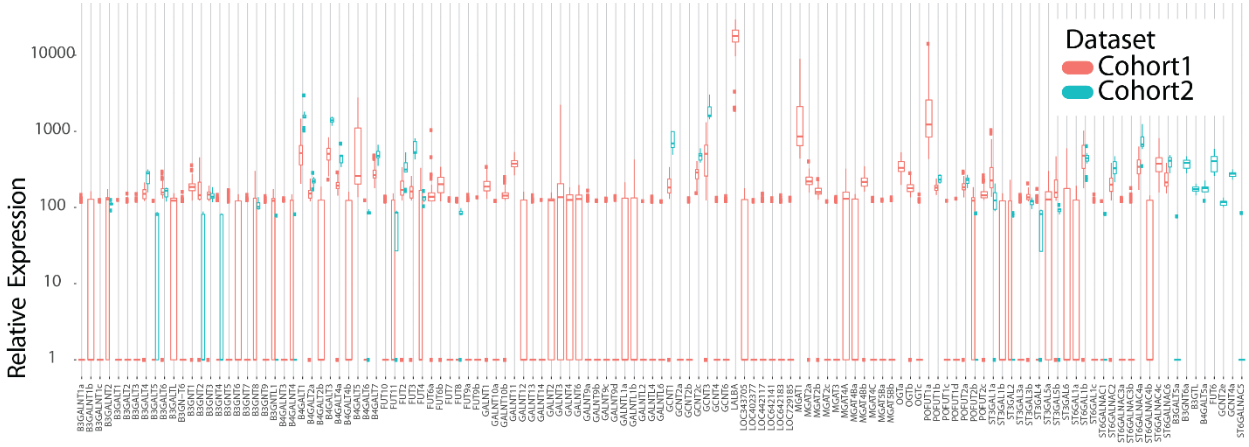

***Figure** S 6* - ***Distribution of Gene Expression Values of Each Gene Candidate.*** *Distribution of gene expression values from microarray experiments for each gene examined across all samples stratified between dataset1 (red) and dataset2 (blue).*

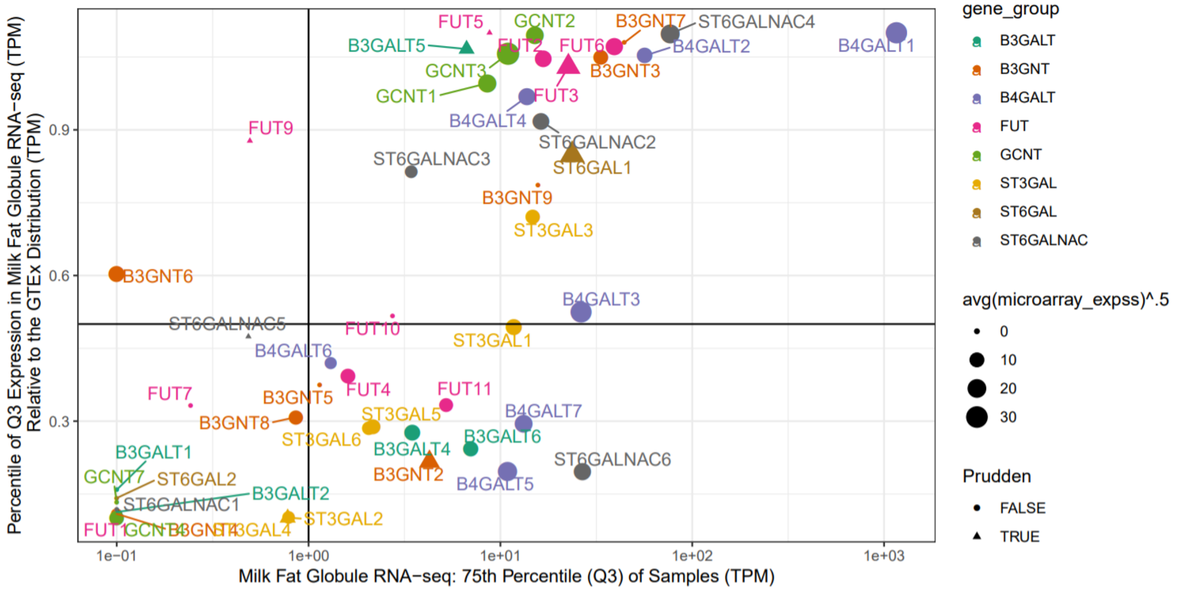

***Figure** S 7* - **Expression (75th percentile of TPMs, Q3) of glycosylatransferase genes in breast epithelium vs the percentile of Q3-TPM expression relative to GTEx expression**. The RNA-Seq (GSE45669[31]) was used to validate low expression in the microarrays (size). Genes used in Prudden et. al. 2017 were highlighted with triangles.

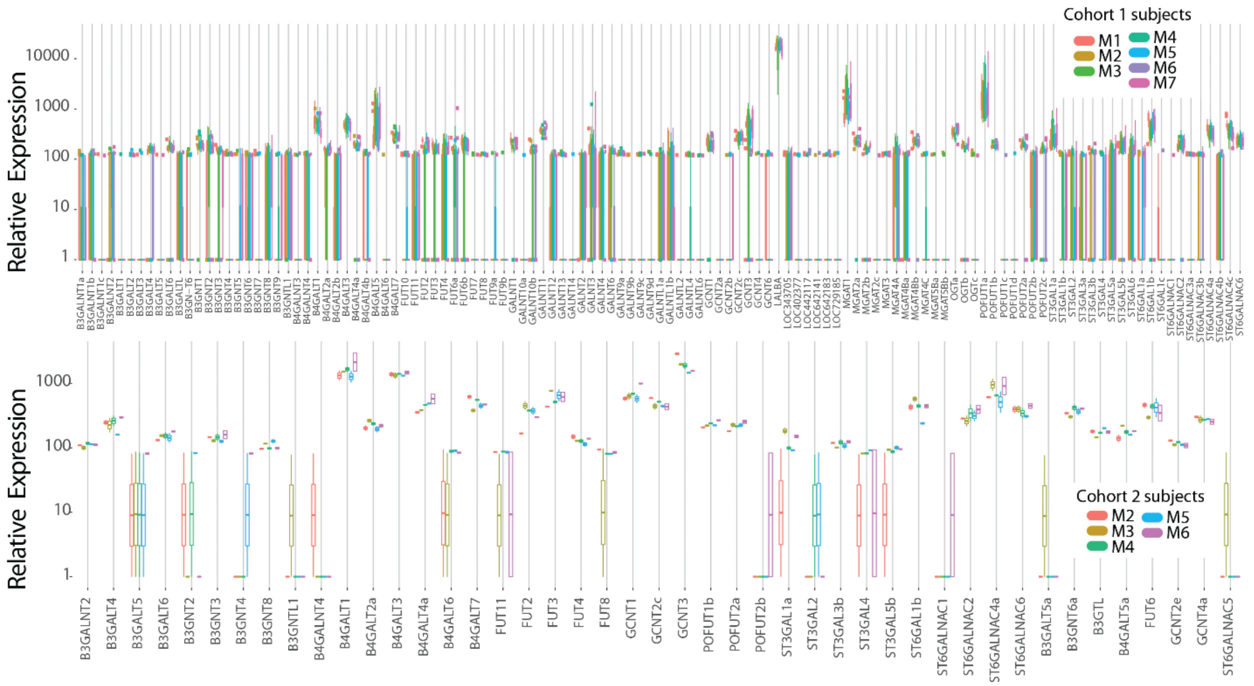

***Figure S 8 - Distribution of Gene Expression Values of Each Gene Candidate Stratified by Subject.*** *Distribution of gene expression values from microarray experiments for each gene examined across all samples stratified between dataset (dataset1 (top) and dataset2 (bottom)), and subject (color).*

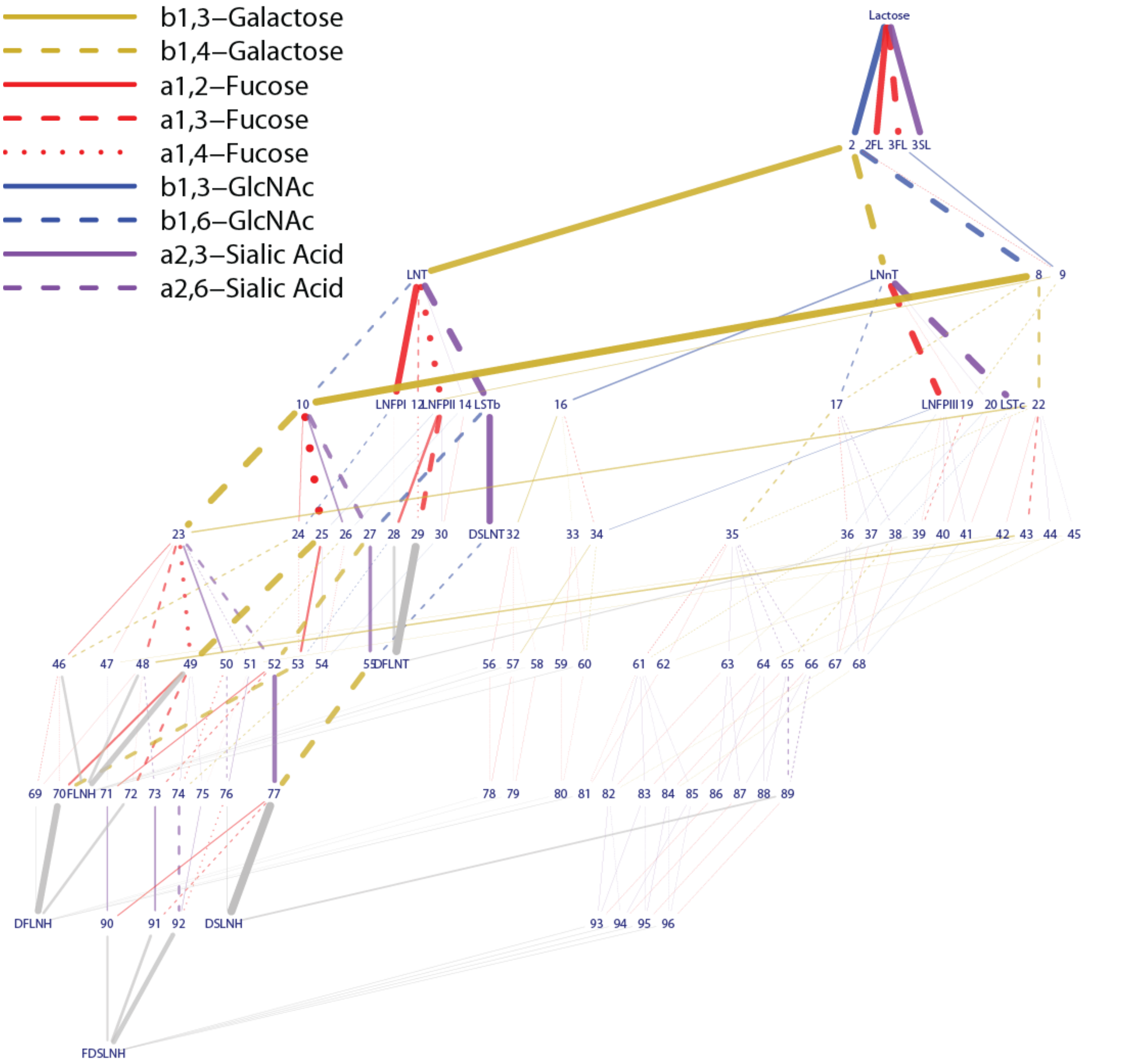

***Figure** S 9 **- The reduced network produced by flux variability analysis.*** *The Reduced Network retains all reactions from the Complete Network,* ***Figure*** *3B, necessary to perfectly predict the HMO profiles collected by High Throughput Liquid Chromatography (HPLC). Numerical nodes are unmeasured intermediary HMOs specified in* ***Table*** *S 2.*

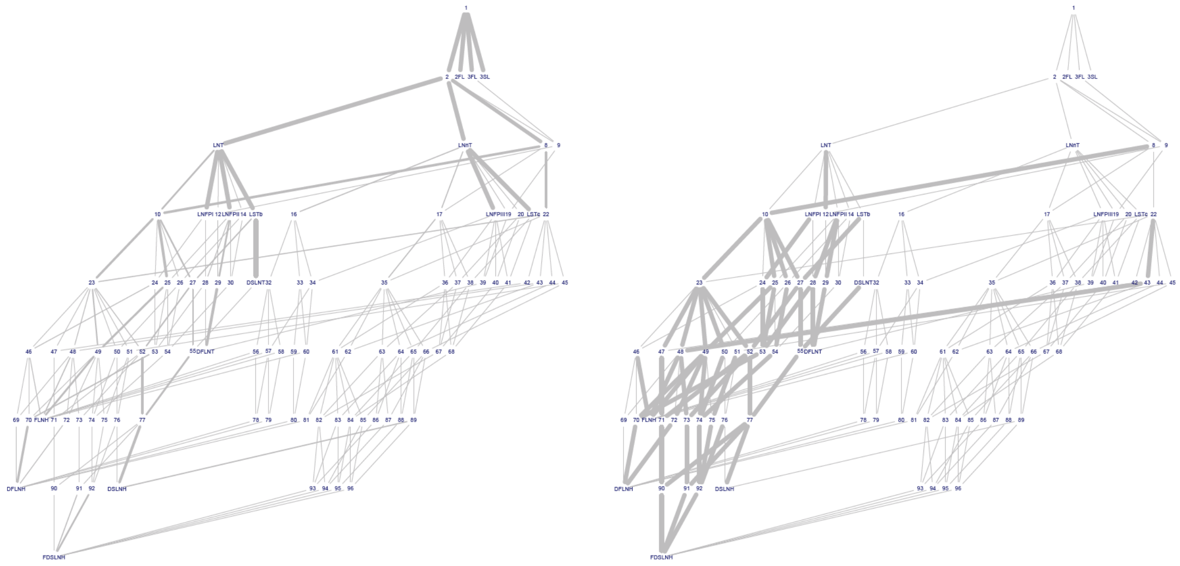

***Figure** S 10* ***- Reduced Network weighted by proportion and enrichment of each reaction in the high-performing models.*** *Edge thickness indicates proportion of top-performing models which include a reaction (left), or enrichment of reaction in the top-performing model set as compared to the background model set (right). Each node is numbered referring to an HMO* ***Table*** *S 2: HMOs*

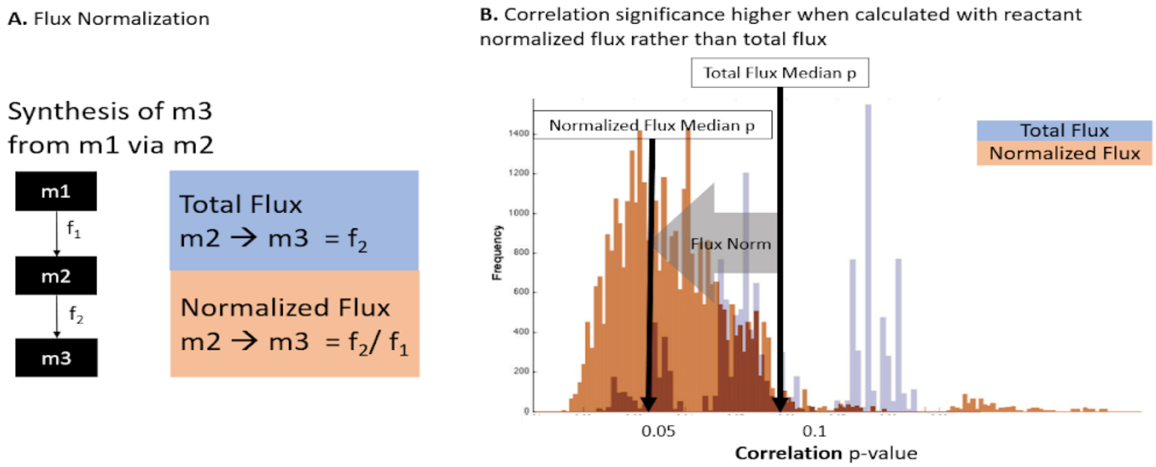

***Figure** S 11* ***- Comparison of total flux and normalized flux.*** *In this method we used prior-normalized flux to control for the availability of the precursor. Considering the enzyme abundance or presence can be informative but it is only one of the limiting factors in a synthesis reaction. Another essential factor to consider is the availability of the precursor.* ***(A)*** *shows the difference in calculating the total flux for reaction 2, conversion of m2 to m3. Total flux is contrasted by normalized flux f2/f1. This is to say that f2 can never exceed f1. If f1 is pulled to multiple reaction, this converts f2 to a measure of the proportion of m2 converted to m3.* ***(B)*** *shows the distribution of p-values for flux-expression Spearman correlations. Clearly, there is a shift towards significance when normalized flux is employed.*

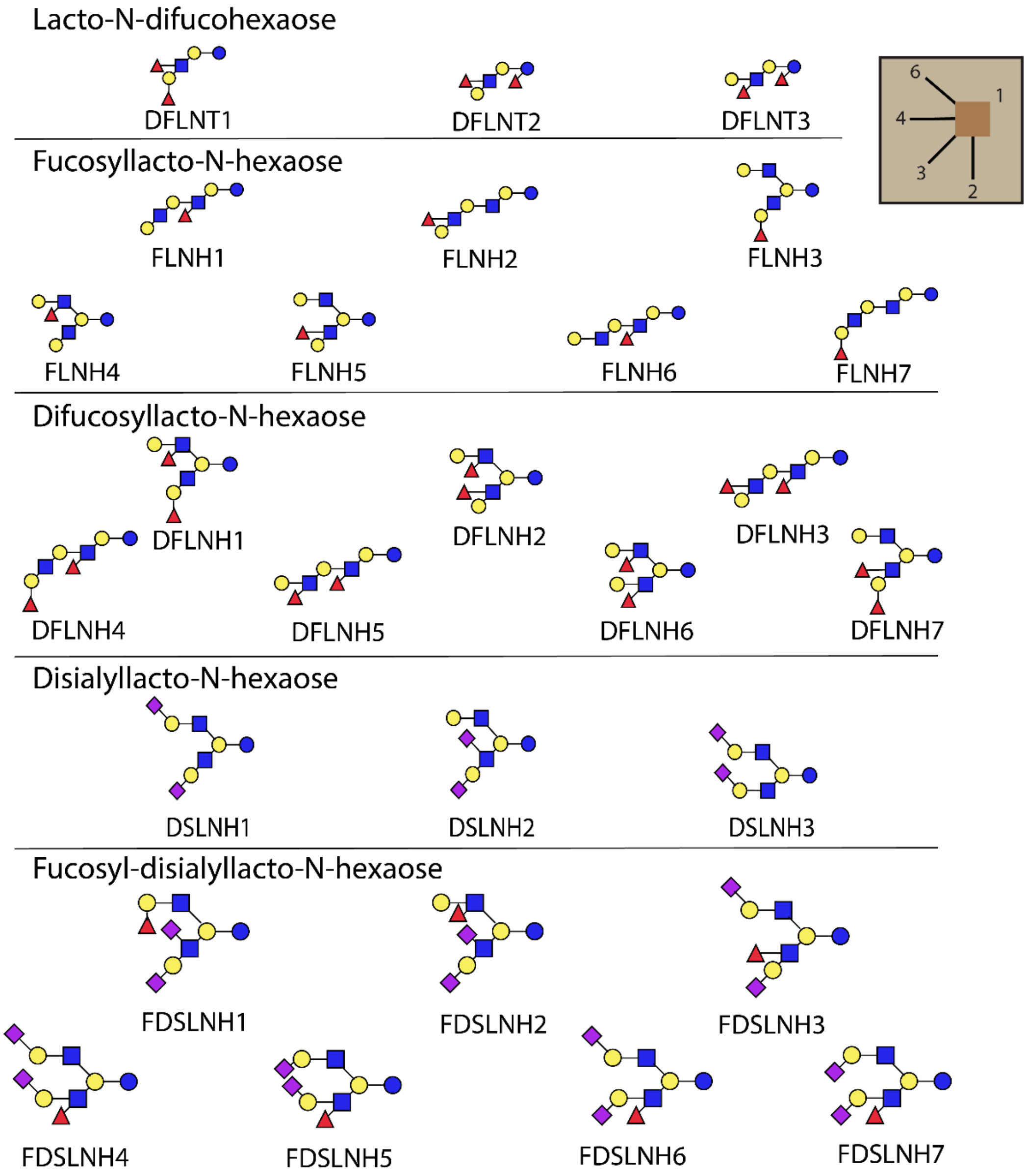

***Figure** S 12* ***- Candidate structures of uncharacterized human milk oligosaccharides.*** *5 HMOs quantified for this study are not structurally well-characterized. Each of these HMOs have been associated to candidates structures identified based on published work[8]. The structures of some HMOs are unclear. Due to the diversity of monosaccharides that could be attached through different linkages, a huge number of structurally distinct HMOs are possible. Of the many possible HMOs, more than 150 have been identified[100]. Several of the most abundant observed HMOs remain to have ambiguous structures. The natural heterogeneity (branching, isomerization and polarization) of HMO mixture present in milk makes their structural identification and quantitative detection a prohibitive challenge to many current studies[6,8,99]. This is due in part to the scarcity of standards. In this study we use High Performance Liquid Chromatography (HPLC) to measure the abundance of the 16 most abundant HMOs. Of the measured HMOs, 11 have fully determined molecular structures, while the remaining five have multiple candidate structures.*

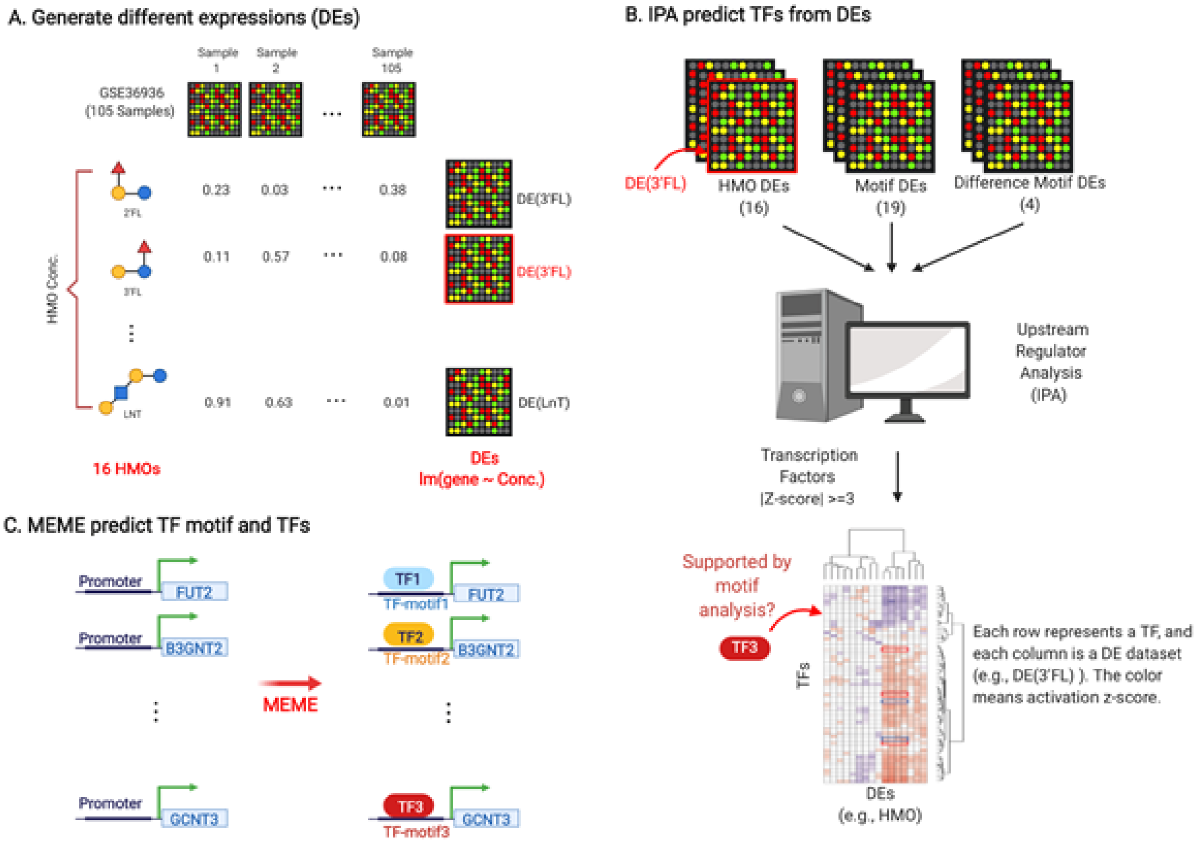

***Figure** S 13 - **Procedures of the Transcriptional Factor Analysis.** **(A)** We first conducted 39 differential expression (DE) analyses to identify gene expression associated with variance in glycan and motif abundance (e.g. limma(expression ~ [3FL])). The 39 abundance measures include: 16 HMO concentrations, 19 glycan motif abundances, and 4 glycan motif abundance ratios. **(B)** We used IPA to predict transcriptional factors (TFs) that could explain differential expression associated with changes in HMO and motif abundance. **(C)** To corroborate the identified TFs, we used MEME to perform TFs binding site analysis for the glycosyltransferases (GTs) based on their promoter sequences.*

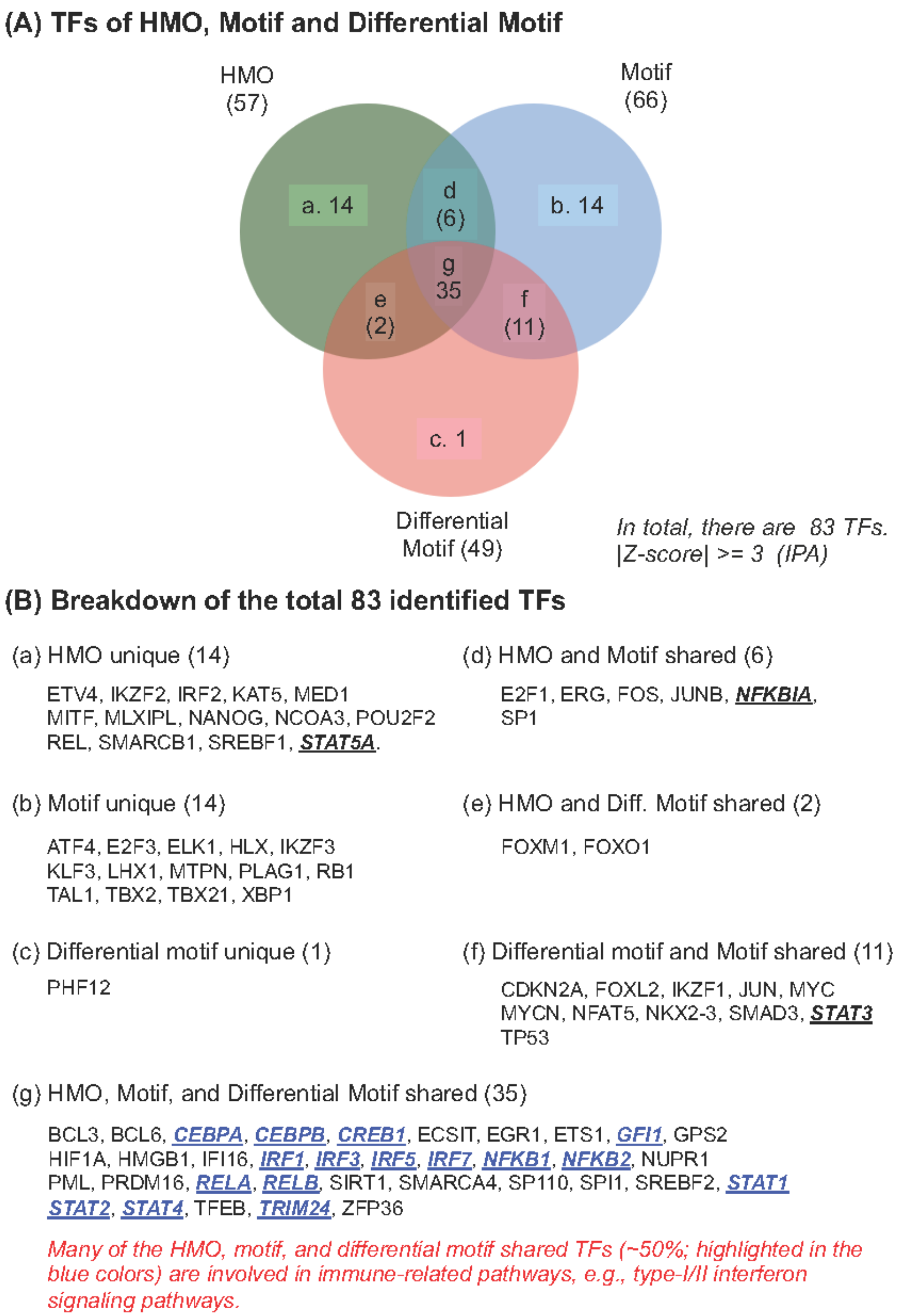

***Figure** S 14* ***- Transcriptional factors (TFs) identified by IPA using the gene expression data.*** *(A) Based on the Z-score predicted by IPA, we identified a total of 83 significant TFs with |Z value|>=3 in this study. (B) We show the breakdown of these 83 TFs into different categories.*

#### A. TF motif – I

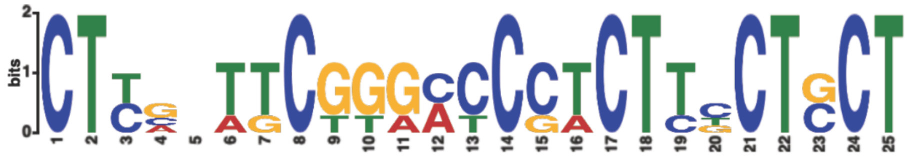

| GlycosylT. | Strand | Start | P-value | Sites |
|---|---|---|---|---|
| GALNTL6 | + | 76 | 2.56e-13 | tcagctctgtCTTGCTTCGGGATCCTCTTCCTCCT |
| ST6GALNAC6 | + | 46 | 1.14e-11 | caaggcctgaCTTCTTTCTGGCCCCACTTTCTGCTgtagtggtct |
| ST3GAL1 | + | 14 | 2.76e-11 | ctgccatcctCTCGGTGCGTGCCCGTCTTCCTGCTgaaacggacc |
| OGT | + | 35 | 4.29e-11 | tgaagacgggCTCAAATCGGAACCCTCTCGCTCCTcgttttgagc |

#### B. TF motif – II

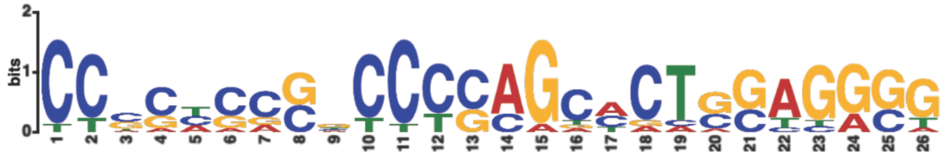

| GlycosylT. | Strand | Start | P-value | Sites |
|---|---|---|---|---|
| B3GNT2 | + | 72 | 1.39e-12 | cggggccattCCTCACCCGCCCCAGCACTGGAGGGGgcg |
| B4GALNT4 | + | 12 | 7.67e-11 | acagctcggcCCCCTGCGGCCCCAGCCCTGCAGGCTgagatcccgc |
| POFUT2 | + | 22 | 2.71e-9 | tgcgcggctcCCGCCCCGACCCCAGCTCCAGAGGGGcgagagcacc |
| POFUT1 | - | 45 | 3.01e-9 | AGCCACCGGTCCCCTCCCTTCCGAGACCTGGAGACGCAGTCCTGAG |
| GALNT9 | + | 54 | 4.12e-8 | gggactcagcTTCCCCAGTCCTCAGCCCTGGACGGGggagagggag |
| MGAT3 | - | 69 | 5.61e-8 | ACATGCCCGAAGCCACCTGCGCACTCCTGGGGTAACTGGGGG |
| GALNT2 | + | 66 | 7.54e-8 | agcaccgaggCCACTACCGCCCCCGTAATCGAGAGGccccgtgtg |
| FUT11 | + | 39 | 1.00e-7 | tcctttccagCTGGAGAGGCCCCAACACTGGATGGTtctgtaggga |
| ST6GALNAC2 | + | 32 | 1.08e-7 | ctccaccctgCCCGCCCCCTCCCAGCTGTCCTGGCGtccgcagact |
| B4GALT6 | - | 18 | 1.62e-7 | GTGAGGCGCCCCCGCCGGGCCCGCGGACACCAGGCGAAGAGCAACC |
| B3GALT6 | - | 60 | 3.03e-7 | CATCGTCCGTCCGCTCGGCCTTGAGCCCTGGCGAGACTCCGCCGCC |

#### C. TF motif – III

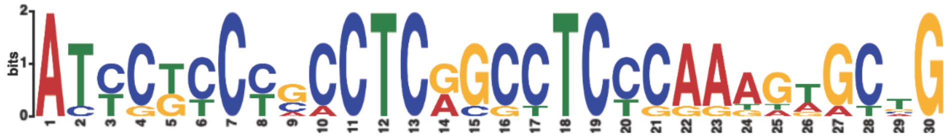

| GlycosylT. | Strand | Start | P-value | Sites |
|---|---|---|---|---|
| B4GALT7 | + | 14 | 1.77e-17 | tgacctcgtgATCCGCCCGCCTCGGCCTCCCAAAGTGCTGggattacagg |
| MGAT1 | + | 2 | 1.23e-16 | gATCCGTCTGCCTCGGCCTCCCAAAGTGCTGggattacagg |
| FUT2 | - | 11 | 3.63e-16 | AGTTCAAGGGATTCTCCTGCCTCAGCTTCCCAAAGTGCTGGGATTACAGA |
| FUT6 | + | 45 | 1.75e-14 | ggttcacaccATTCTCCTGCCTCAGCCTCCCAAGTAGCTGggactacagg |
| B3GNT3 | + | 27 | 1.55e-12 | ggttccagcgATCCGCCCCCCTCGGCCTCTGGAAGTACAGgagtgagcca |
| FUT10 | + | 58 | 6.86e-12 | gactcaagagATCCTCCCACCTCAGCCTCCCAGTAGGTGGgactacaggc |
| ST3GAL2 | - | 27 | 2.69e-11 | GCCGGAGGGCACTGTTCCCACTCGCGCTCTCAAAGAGCCGCGACCTGAGG |

***Figure** S 15 – **TF motifs identified by MEME using the promoter sequence of GTs.** There are three TF motifs that their known TFs overlapped with the IPA predict TFs: (A) TF motif-I. (B) TF motif-II, and (C) TF motif-III. The red arrows highlighted GTs are those major GTs involved in the HMO biosynthesis selected in this study (**Figure** 5**C** in the main text). The p-value represents the significance of the MEME identified TF motif associated with the GT by the aligned potential TF binding site on the promoter sequence.*

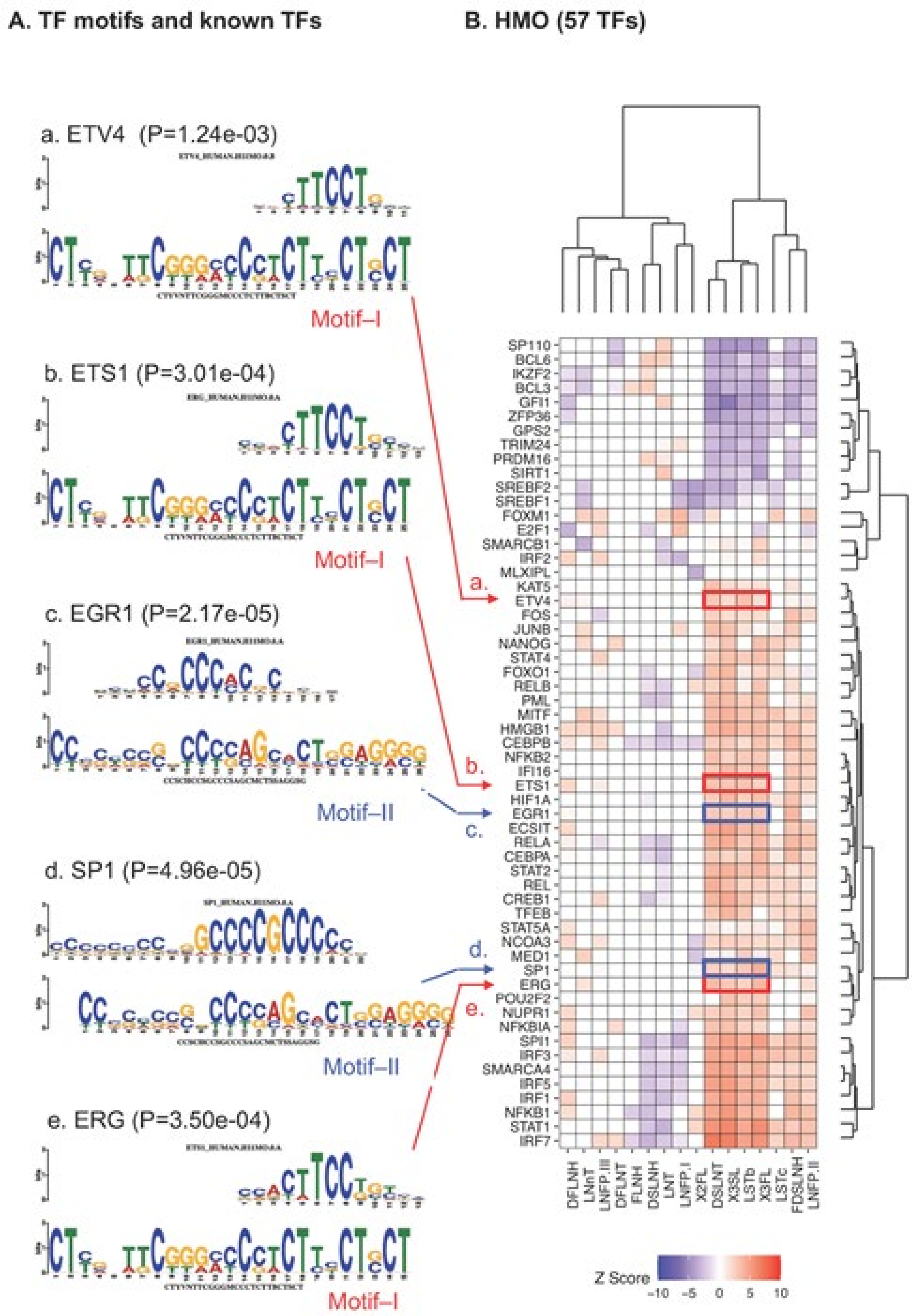

*Figure S 16* – ***MEME promoter-enriched TF motifs and IPA predicted TFs using differential expression analyses with respect to 16 HMOs.*** *(A) MEME identified TF motifs and 5 known TFs (ETV4, ETS1, EGR1, SP1, and ERG) associated with them (see* ***Table*** *S 6). MEME-discovered TFs were cross-referenced with known TF binding sites using TOMTOM. Logos for the matched known and discovered motifs are shown in the top and bottom of each subpanel (A.a-e); the p-value is a logo matching significance calculated by TOMTOM. (B) Biclustering of activation z-score computed by IPA indicating the likelihood that a TF activates (z>0) or inhibits (z<0) an HMO concentration signature (gene expression associated with changes in HMO concentration).*

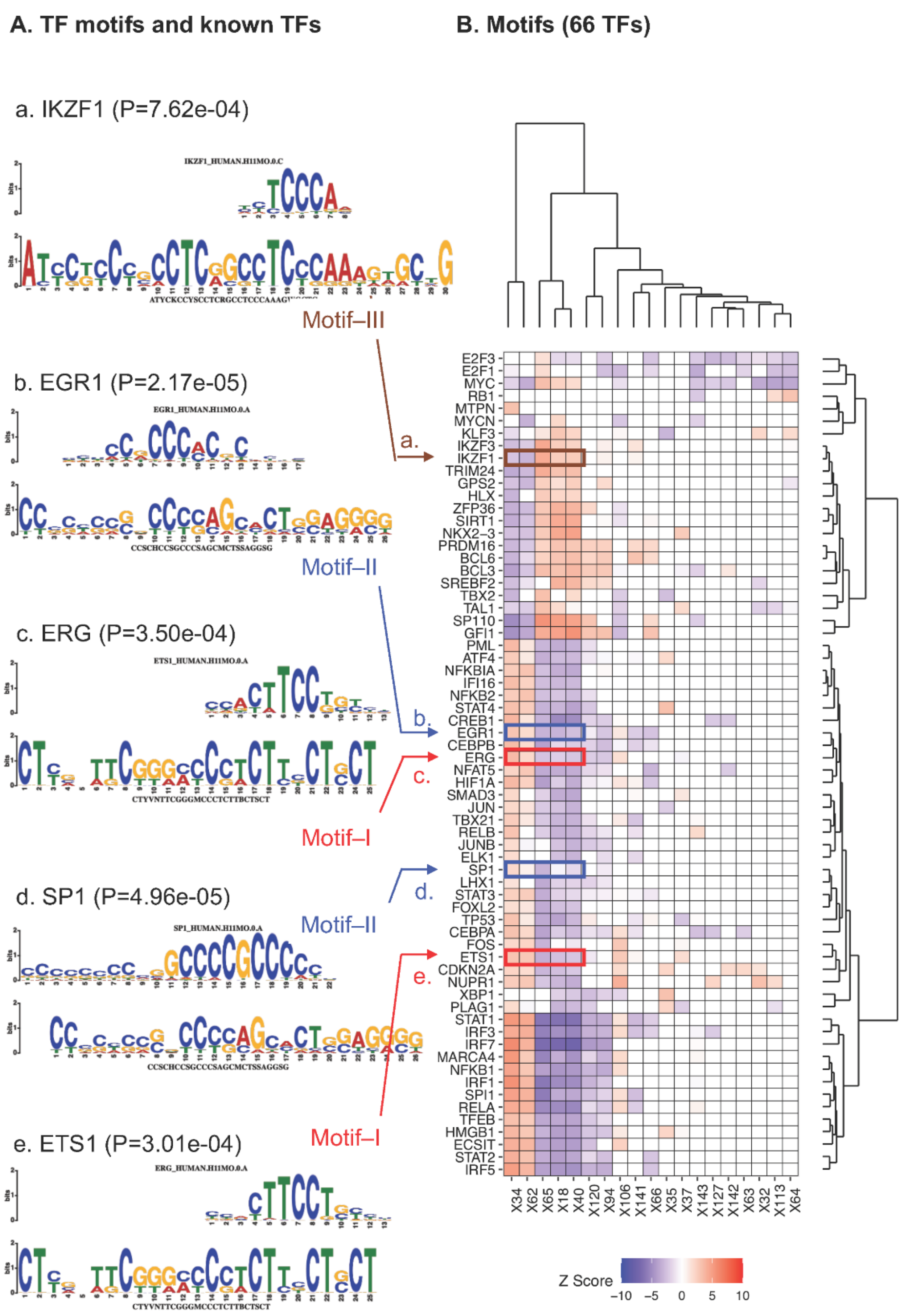

***Figure** S 17 – Clustering results and MEME identified TFs of the IPA predicted TFs using the gene expression data (19 Motifs).* *(A) MEME identified TF motifs and 5 know TFs (IKZF1, EGR1, ERG, SP1, and ETS1) associated with them (see **Table** S 1). The known TF binding site motifs discovered by TOMTOM is given on the first line of each box, followed by the motif discovered by MEME, all shown as aligned logos. The p-value showing in the parenthesis represents the significance of the known TF motif associated with the MEME identified TF motif. (B) Clustering results of the IPA identified TFs using the 19 motif DE data. The color in the heatmap denotes the IPA predicted activation (Z-score) of each TF in the indicated glycan motif. X[number] glycan substructures defined in **Figure** S 19.*

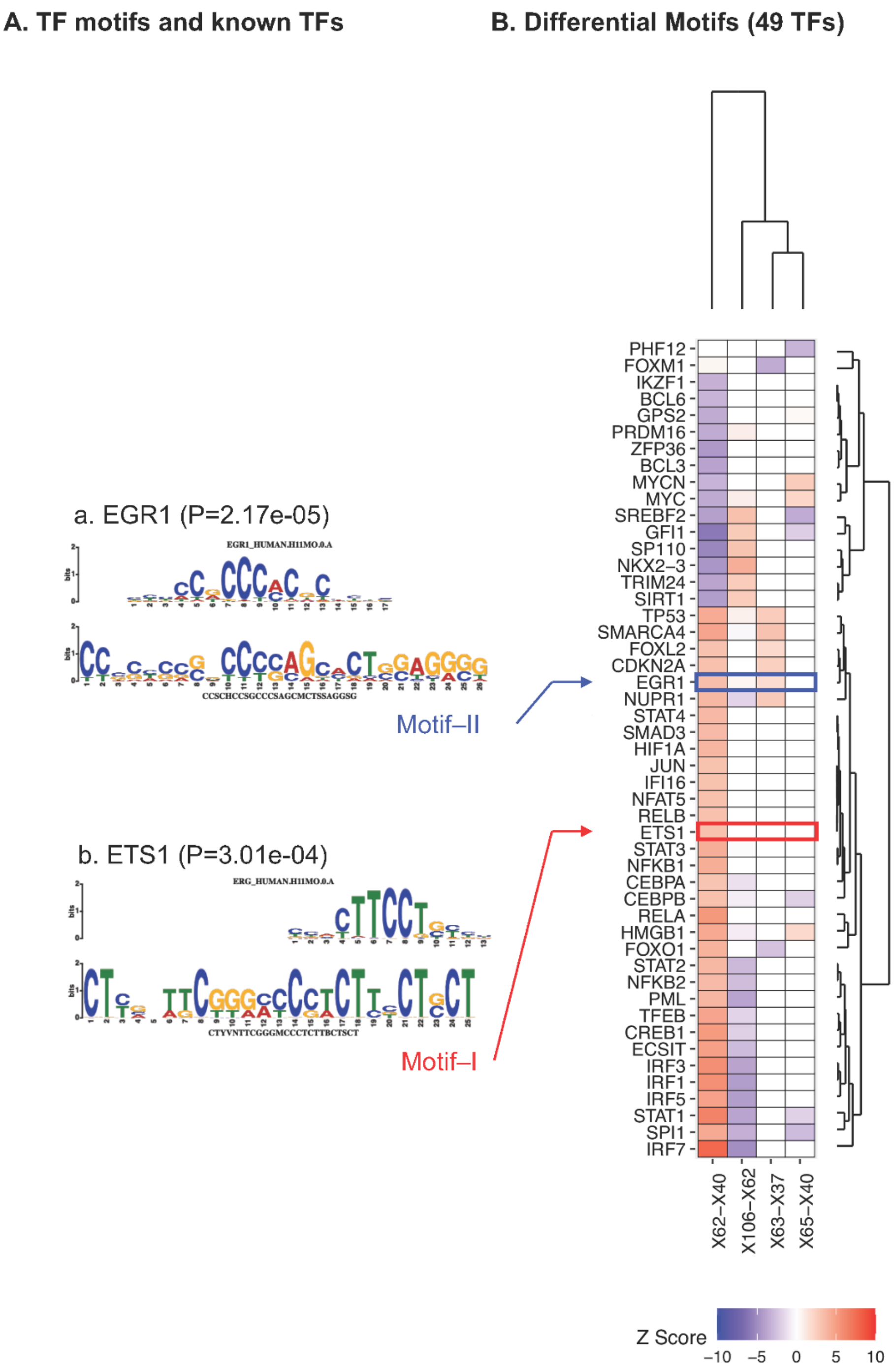

***Figure S 18 – Clustering results and MEME identified TFs of the IPA predicted TFs using the gene expression data (4 differential motifs).*** *(A) MEME identified TF motifs and two know TFs (EGR1 and ETS1) associated with them (see **Table** S 1). The known TF binding site motifs discovered by TOMTOM is given on the first line of each box, followed by the motif discovered by MEME, all shown as aligned logos. The p-value showing in the parenthesis represents the significance of the known TF motif associated with the MEME identified TF motif. (B) Clustering results of the IPA identified TFs using the differential motifs DE data. The color in the heatmap denotes the IPA predicted activation (Z-score) of each TF in the indicated differential motif. X[number] glycan substructures defined in **Figure S 19**.*

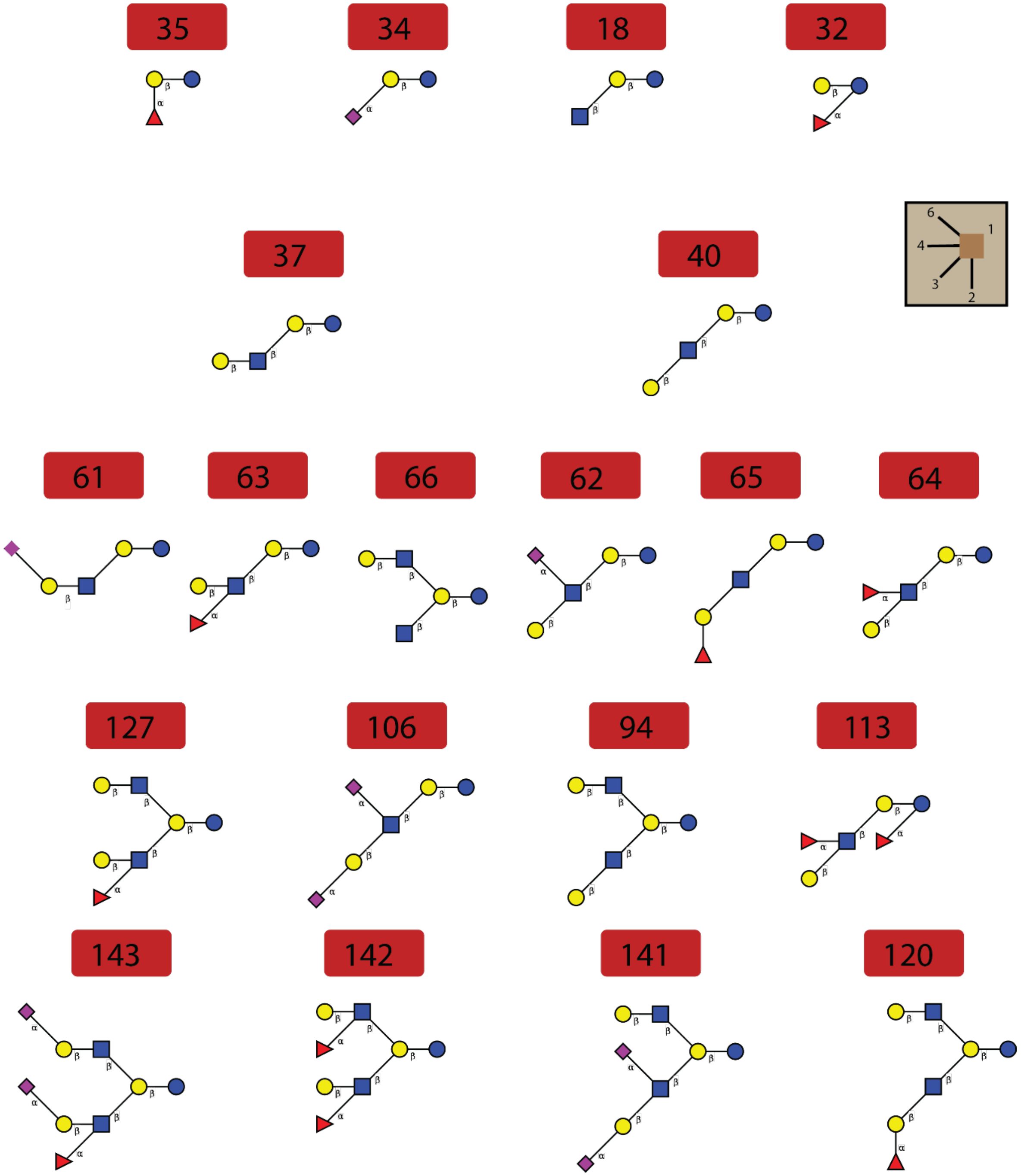

***Figure S 19 - HMO substructure network with dependent substructure removed.*** *All the glyco-motifs examined in the IPA transcription factor analysis. Reproduced with permission*[17]

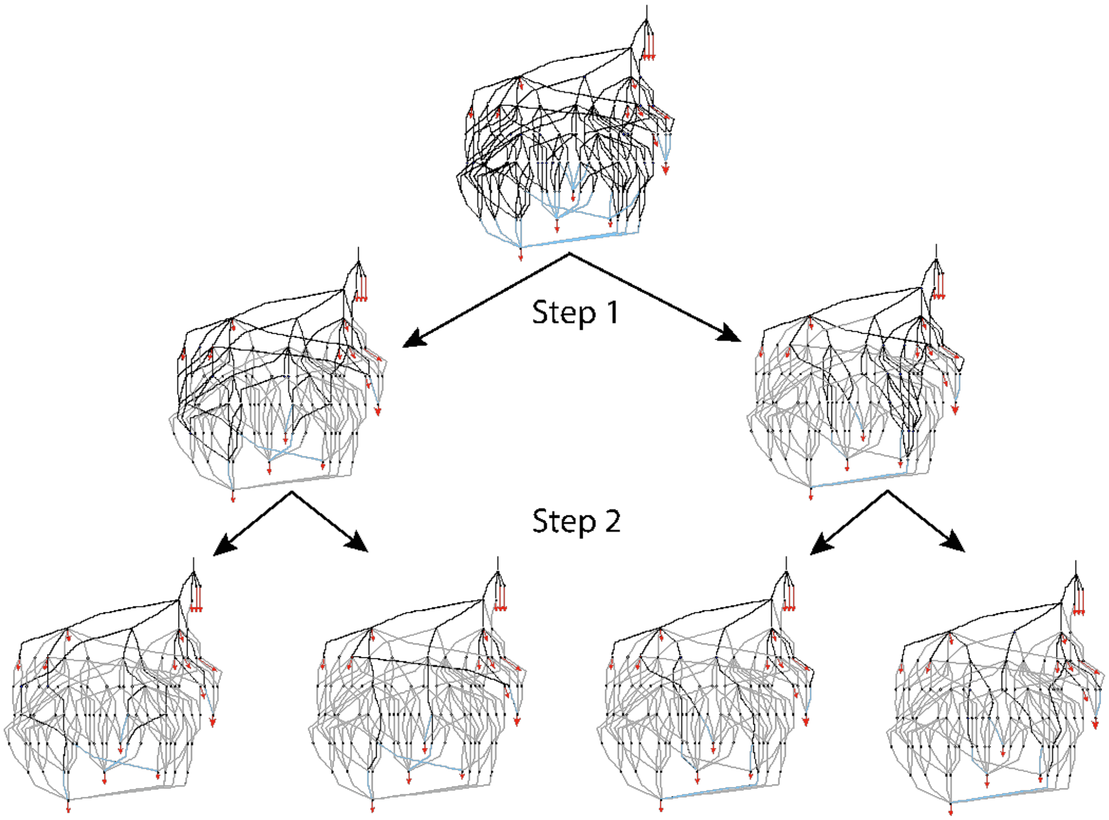

***Figure** S 20 - **Subnetwork generation methodology.** Different combinations (i.e. only two combinations of the 5 uncharacterized HMOs are shown in the **Figure**) are fixed to tailor the solution space (step 1) and a MILP algorithm is further applied to these reduced networks to enumerate all alternate subnetworks (step 2, only two alternate subnetworks by combination are showed in this **Figure**) that can synthesize all HMOs of interest.*

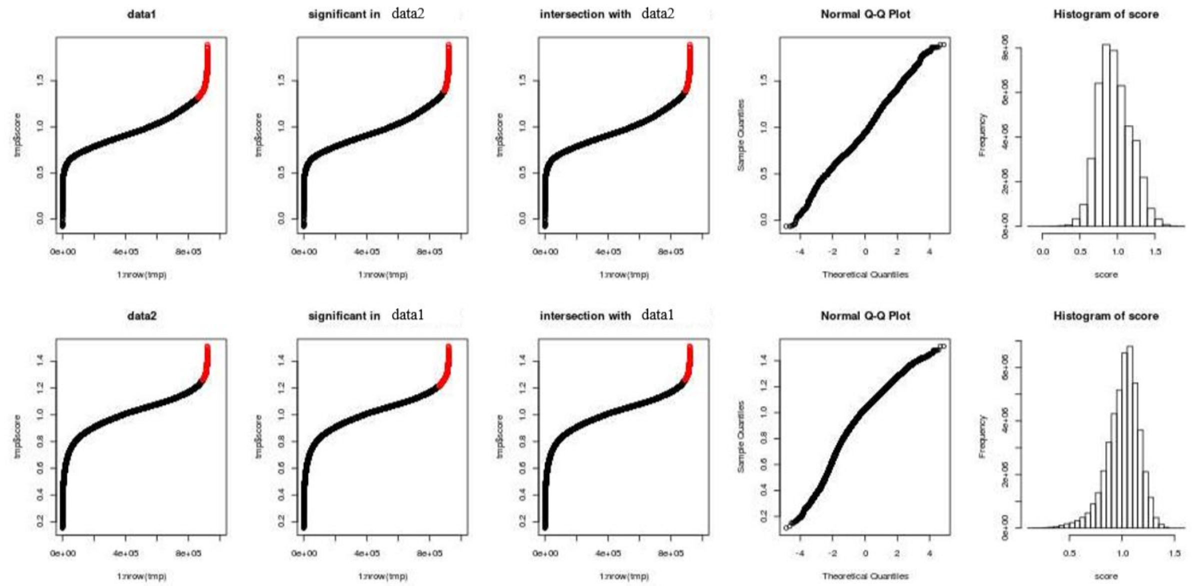

***Figure** S 21 - **Selection of models presenting a high model score** – (letters will be added by row, left to right) A-C. Candidate Model Scores, $S_{\omega}(D_1)$, sorted by magnitude; sorted index vs. magnitude. F-H. Candidate Model Scores, $S_{\omega}(D_2)$, sorted by magnitude; sorted index vs. magnitude. A and F show the sorted scores for $S_{\omega}(D_1)$, and $S_{\omega}(2)$, respectively. Red points are top-performing candidate models. B and G show the top-performing models (red) from F and A respectively mapped onto the distributions from A and F to show cross-comparability. C and H show the commonly top-performing models (red) between dataset1 and dataset2 mapped onto score sigmoid 1 and 2 respectively to show the models in the intersection perform well. D and I show Q-Q plots of the score distributions to evaluate the normality of the distribution. E and J show distributions of the scores to further visualize normality of the distributions.*

### 3 SUPPLEMENTARY TABLES

***Table** S 1 – **Linkages and genes considered throughout this analysis.*** *List of glycosyltransferase genes associated with each reaction and associated expression in datasets within and beyond this work. Here we list the 56 candidate glycosyltransferase genes considered. Columns **A-E** indicate the corresponding reaction, gene name and various identifiers. Column **G**, "Prudden," indicates if the gene was used Prudden et. al. 2017. Columns **H-J** show expression on cohort 1 and 2 microarrays (relative abundance) and an independent RNA-seq in the same tissue (TPM). Column **M-O** specify the distribution of TPMs in GTEx (min, Q1, Q2, Q3, Q4, max), the percentile where Q3 from the RNA-seq falls in the GTEx distribution, and the log2 Fold Change of RNA-seq Q3 vs GTEx Q3. Column **P** specifies any low expression events. Columns **Q-S** compare microarray and RNA-seq expression and usage in Prudden et. al. 2017. Column **T** indicates if previous literature found a gene relevant ("yes"), partially relevant ("similar") or irrelevant ("no") to the reaction in question; elaborated in **Table** S 1. Columns **U-V** delimited and justify which genes will be included for further analysis. The remailing columns provide some relevant detail on substrate specificity for each gene.*

See associated file: **Table** S 1

***Table** S 2 - **The FVA reduced HMO biosynthetic network including HMO and reaction calculation referenced throughout the manuscript***

See associated file: **Table** S 2

***Table** S 3 - **Reduction in model complexity across various examined models.***

| model Size | Complete | Reduced | Candidate |
|---|---|---|---|
| HMOs | 2,961 | 101 | 27-38 |
| Reactions | 9,918 | 221 | 43-54 |

***Table** S 4 - **Number of total subnetworks High-performing candidate models from given dataset 1 or 2 with all sample or just secretor samples from the background of 44,984,988 subnetwork models.***

| Data | Dataset 1 | Dataset 2 | Intersection |
|---|---|---|---|
| All Data | 2,658,052 | 2,322,262 | 241,589 |

***Table S 5 – Results of the NTP-Glo™ Glycosyltransferase Assay (Promega) to test GT candidates on relevant HMO acceptors.*** *++ indicates a strong effect, + indicates a weak effect, - indicates an insignificant effect, a blank cell indicates a negligible effect and a grey cell indicates an unmeasured reaction.*

| Linkage | Reaction | Enzyme | Lactose | LNT | LNnT | Gal b-1,3-GalNAc |
|---|---|---|---|---|---|---|
| L1 b3GnT | b-1,3 N-acetylglucosamine | B3GNT2 | ++ | | ++ | |
| L4 ST3GalT | (b-Gal) a-2,3 sialytransferase | ST3GAL1 | | + | | ++ |
| | | ST3GAL2 | | - | | + |
| | | ST3GAL3 | | ++ | ++ | + |
| | | ST3GAL4 | | - | + | - |
| | | ST3GAL5 | | | | |
| | | ST3GAL6 | | + | ++ | - |
| L10 ST6GnT | (b-1,3-GlcNac) a-2,6 sialytransferase | ST6GALNAC2 | | | | |
| | | ST6GALNAC5 | | - | | |

***Table S 6 – MEME discovered 3 novel TF motifs and their associated known TF motifs that were supported by IPA predicted TFs***

| MEME TF binding site motif | Known TF | Known TF binding site consensus | Overlap | P-value[*1] |
|---|---|---|---|---|
| **TF motif – I** 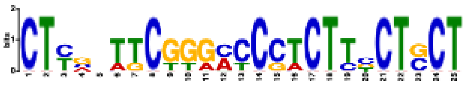 | ETV5 | CTCACTTCCTGCTC | 13 | 6.74E-06 |
| | ELF5 | CCACTTCCTCCTTCC | 12 | 1.78E-04 |
| | ETS2 | TCCTCTTCCTTCC | 13 | 2.58E-04 |
| | **ERG[*2]** | **CCACTTCCTGCCC** | **12** | **3.01E-04** |
| | **ETS1[*2]** | **CCACTTCCTGTCT** | **12** | **3.49E-04** |
| | ELF2 | CCACTTCCGGGTT | 12 | 6.75E-04 |
| | EHF | GCCACTTCCTGGTTC | 13 | 1.03E-03 |
| | **ETV4[*2]** | **CCCTTCCTGTT** | **11** | **1.24E-03** |
| | ELF3 | CCACTTCCTGGTTC | 12 | 1.26E-03 |

| | | | | |
|---|---|---|---|---|
| | SP4 | CCCGGCCCCGCCCCCTTCCC | 19 | 1.33E-03 |
| | ETV2 | CCCACTTCCTGTTTCC | 13 | 1.62E-03 |
| | NKX25 | CCTCTCCA | 8 | 1.94E-03 |
| | GABPA | CCACTTCCGGTTCC | 12 | 2.39E-03 |
| | **EGR1[*2]** | **CCCCCGCCCACGCCCTC** | **17** | **2.17E-05** |
| | PATZ1 | CCTCCCCCCCGCCCCCTCCCC | 19 | 3.62E-05 |
| | **SP1[*2]** | **CCCCCCCCCGGCCCCGCCCCCC** | **20** | **4.96E-05** |
| | SP3 | CCCCGGCCCCGCCCCCCCCC | 18 | 1.08E-04 |
| | KLF1 | CCCGGCCCCGCCCC | 14 | 3.27E-04 |
| **TF motif – II** | KLF15 | CCCCCCCCTGCTCCTCCCC | 19 | 4.10E-04 |
| | WT1 | CCCCCCCTCCTCCCCCGCCC | 18 | 4.33E-04 |
| 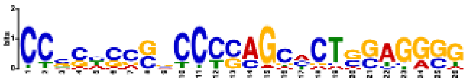 | SP2 | CCCCCGGCCCCGCCCCCCCCCC | 19 | 7.52E-04 |
| | KLF6 | CCCCCGGCCCCGCCCTTCC | 19 | 8.71E-04 |
| | E2F6 | CCCTTCCCGCCCC | 13 | 1.02E-03 |
| | ZN770 | GATCCTCCCGCCTCAGCCTCCC | 21 | 1.65E-03 |
| | ZBTB6 | CGGCTCCAGCACC | 13 | 1.91E-03 |
| | MAZ | CCCCCCCCCCCCCCCCTCCCCC | 20 | 2.13E-03 |
| | ZN770 | GATCCTCCCGCCTCAGCCTCCC | 21 | 2.72E-11 |
| | ZSC22 | CCTCCTCCCTCAGACC | 16 | 2.29E-04 |
| | WT1 | CCCCCCCTCCTCCCCCGCCC | 20 | 5.61E-04 |
| | ZN263 | CTCCTCCTCTCCCTCCTCCC | 20 | 5.61E-04 |
| **TF motif – III** | **IKZF1[*2]** | **TCTCCCAA** | **8** | **7.62E-04** |
| | PATZ1 | CCTCCCCCCCGCCCCCTCCCC | 22 | 1.40E-03 |
| 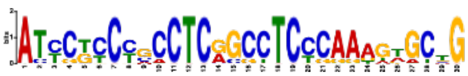 | KLF6 | CCCCCGGCCCCGCCCTTCC | 19 | 1.45E-03 |
| | MAZ | CCCCCCCCCCCCCCCCTCCCCC | 22 | 1.84E-03 |
| | ZN281 | TCCCCTCCCCCACCC | 15 | 2.01E-03 |
| | ZN467 | CCCCCCCCCCCCTCCCCTCCCC | 22 | 2.10E-03 |

*1 The P-value (see **Figure** S 13, **Figure** S 17, **Figure** S 18) is the significance of know TF identified to the MEME identified TF motif.

*2 The red highlighted six known TFs (IKZF1, EGR1, SP1, ERG, ETS1, and ETV4) are the ones that also predicted as upstream regulators by IPA using gene expression data (**Figure** S 16, **Figure** S 17, **Figure** S 18).

***Table** S 7 – **MEME discovered 7 novel TF motifs and their associated known TF motifs that were not supported by IPA predicted TFs.***

| **MEME TF binding site motif** | **Known TF** | **Know TF binding site consensus** | **Overlap** | **P-value[*1]** |
|---|---|---|---|---|
| **TF motif – IV** 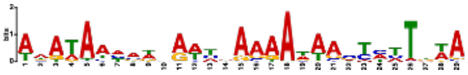 | MEF2D | CTAAAAATAGCA | 12 | 3.06E-04 |
| | AIRE | TTAACCAATAAAACCAAT | 18 | 3.41E-04 |
| | PRDM6 | AAAAAGAAAAAAA | 13 | 7.98E-04 |
| | MEF2C | GCTAAAAATAGCA | 13 | 7.98E-04 |
| | MEF2B | CCAAAAATAGCAAC | 14 | 8.02E-04 |
| | FOXJ3 | AAAAAATAAACAA | 13 | 1.15E-03 |
| | ANDR | AGCAAACAAAAAAGAACA | 18 | 1.43E-03 |
| | NKX61 | CATTAAATTCCCATTAATC | 17 | 1.57E-03 |
| | MEF2A | GCTAAAAATAGAA | 13 | 2.03E-03 |
| **TF motif – V** 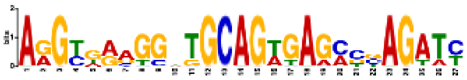 | TBX3 | GGAGGTGGGAA | 11 | 1.04E-03 |
| | MYB | GGTGGCAGTTGG | 12 | 1.83E-03 |
| | RARG | AGGTCAGAGTGACCTGGG | 18 | 1.90E-03 |
| | ETV5 | GAGCAGGAAGTGAG | 14 | 2.12E-03 |
| | SMAD2 | AGGTGACAGACA | 12 | 2.40E-03 |
| **TF motif – VI** 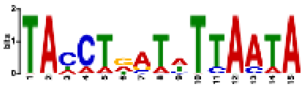 | TWST1 | ACATCTGGTTTTAATTA | 15 | 1.18E-03 |
| **TF motif – VII** | ZN320 | GTGGGACCAGGGGGCCAGTG | 14 | 1.66E-03 |

| | | | | |
|---|---|---|---|---|
| 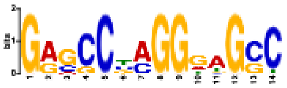 | | | | |
| **TF motif – VIII** | ZN257 | AGAGGCAAGAGC | 12 | 5.47E-04 |
| 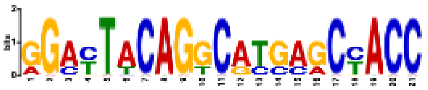 | TEAD4 | ACATTCCTGGCAT | 13 | 2.49E-03 |
| | SOX2 | TCCCTTTGTTCTC | 13 | 1.06E-04 |
| **TF motif – IX** | PRDM6 | TTTTTTTCTTTTT | 13 | 7.26E-04 |
| | ZN680 | CCTCATTCTTCTTGGACATG | 20 | 1.21E-03 |
| 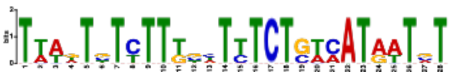 | NFAC1 | TTTCTTTTTTTCCAT | 15 | 1.22E-03 |
| | SOX4 | TCCTTTGTTCTC | 12 | 1.31E-03 |
| | CDX2 | TTTTATTGCTGT | 12 | 1.99E-03 |
| **TF motif – X** | ZN502 | GAATGGAATCGAATGGAATC | 14 | 1.65E-03 |
| | ZN140 | TAGGAGCGGAATTGCTGGGTCATA | 14 | 1.80E-03 |
| 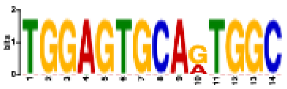 | ZNF18 | GGTGTGAACTGG | 12 | 2.09E-03 |

*1 The P-value (see **Figure** S 13, **Figure** S 17, **Figure** S 18) is the significance of know TF identified to the MEME identified TF motif.

*2 The red highlighted six known TFs (IKZF1, EGR1, SP1, ERG, ETS1, and ETV4) are the ones that also predicted as upstream regulators by IPA using gene expression data (**Figure** S 16, **Figure** S 17, **Figure** S 18).

### 4 SUPPLEMENTAL METHODS

#### 4.1 GENERATION OF COMPLETE HMO BIOSYNTHESIS NETWORK

Similar to Spahn et al (2016), a generic reaction network for HMO biosynthesis is constructed based on a set of reaction rules (**Figure 3**A).

Based on the knowledge of enzyme specificities for each structure feature, the algorithm constructs the complete biosynthesis reaction network required to generate all HMOs of a user-defined complexity level. Specifically, the complexity level $CL^k$ refers to the set of HMOs that can be produced from the initial lactose in $k$ reaction steps (iterations) or less, following the reaction rules (**Figure 3**B).

The network generated is a complete network representing all reactions that, theoretically, could occur in human for a defined HMO structure complexity level, given the known enzyme specificities. We set the maximum complexity level to $CL^7$, as it captures the most complex HMO within the scope of this study (FDSLNH candidates, **Figure** S 12). In total, the network includes 9918 reactions and unique 2961 HMO structures.

#### 4.2 REDUCTION OF THE GENERIC NETWORK TO A REDUCED NETWORK USING FLUX VARIABILITY APPROACH

To make the network specific to the HMOs of interest in this study, the generic network is tailored by removing all the reactions that are not required to describe the synthesis of 16 measured HMOs. First, several additional reactions were introduced into the generic network to account for the structural ambiguity of uncharacterized HMOs (**Figure 3**B-C).

Second, the size of the generic network is reduced by removing all reactions that cannot contribute to the synthesis of the measured HMOs using a Flux Variability Analysis (**Figure 3**C, S1.2.4). The reduced generic network includes 221 reactions and 101 HMO structures (**Figure** S 9).

#### 4.3 GENERATION OF ALL MINIMAL SUBNETWORKS FOR EVERY POTENTIAL HMO STRUCTURE COMBINATION

From the reduced generic network, it is possible to enumerate all the subnetworks presenting specific combinations of candidate HMO structures for a defined number of reactions using an MILP approach (S1.2.5). To this end, all potential combinations of the HMOs associated with unknown structures have first been generated (i.e. number of combinations equal to 3087). For each combination, we used a mixed integer linear programming algorithm to extract all subnetworks, which present unique biosynthetic pathways for every HMO (**Figure** S 20). Doing so, the number of reactions included in the subnetwork will range between the minimal number of reactions required for the simultaneous biosynthesis of all observed HMOs and the maximal number of reactions after which the extracted subnetwork no longer present unique solution when predicting the flux distribution. This methodology allowed the systematic analysis of the entire solution space of the reduced generic network and identification of 44,984,988 existing alternative minimal subnetworks capable of synthesizing all measured HMOs.

#### 4.4 ASSESSMENT OF CONSISTENCY OF EXPERIMENTAL DATA WITH SUBNETWORKS AND SELECTION OF CANDIDATE MODELS

We computed a normalized flux, i.e. the sum of the ratio between the flux of a reaction and its downstream reaction flux for every reaction associated to a specific linkage type within a subnetwork. For this, we first computed the flux values for every reaction within each subnetwork based on a measured glycoprofile in a milk sample, using Flux Balance Analysis (FBA, *S1.2.4*). Since subnetwork extraction identifies the minimum number of reactions, no alternate optimal solutions were needed, thus resulting in a unique flux distribution for each subnetwork. We normalized the flux value (f) of a reaction by the value of flux associated with the previous reaction (f') within the subnetwork to obtain a ***flux ratio*** ($\underline{f}$) for each reaction.

$$\underline{f} = f/f'$$

Based on the assumption that only one enzyme catalyzes the formation of a linkage type, we compute the ***normalized flux*** ($\underline{f}_{l\omega D_m}$) as the sum of all flux ratios associated with a specific linkage ($l$). Since each flux ratio corresponds to a candidate model, $\omega$, and parameterized on the glycomic data, $D_m$:

$$for\ a\ given\ \omega\ \&\ D_m, \qquad \underline{f_l} = \sum_{r \in l_r} \underline{f_r}$$

Doing so, every subnetwork is associated with 10 normalized flux values for each HMO measurement. The normalized flux values represent the activity levels of the 10 types of transferase reactions required for the biosynthesis of measured HMOs. We then computed the Spearman correlations ($\rho$) within linkage, $l$, between the flux ratio, $\underline{f}_{l\omega D_m}$, and the mRNA expression data ($E: g \rightarrow D_g$) of one of its associated candidate genes ($g \in l$, **Table 1**) for the subnetwork ($\omega$). The ***gene-linkage score*** ($\rho_{gl}$) is defined as

$$for\ a\ given\ \omega, D_g, \&\ D_m, \qquad \rho_{gl} = cor(E(g_l) \sim \underline{f_l})$$

##### 4.4.1 Computation of Subnetwork Score

The ***model score*** ($S_\omega$) for each candidate model $\omega$ is defined as the average of the maximum z-score normalized gene-linkage score. The z-score, *z()*, mean and standard deviation is computed from the correlations of all gene-flux correlations within a linkage. The function z will denote this normalization from a set of correlations $x$ to a normalized set of values relative to other correlations within that linkage $z_l: x \rightarrow N(x, \mu_l, \sigma_l)$.

$$for\ a\ given\ \omega, D_g, \&\ D_m, \qquad S_\omega = mean_{l \in L}\ \{\ z_l(max_{g\ \in\ l_g}\ \rho_{gl}\ )\}$$

The model score ($S_\omega$) describes the average consistency between the most consistent genes and the associated fluxes in a subnetwork model.

###### 4.4.2 Ranking subnetwork performance and extraction of candidate models

The distribution of subnetwork scores is approximately normal. Thus, we rely on a z-statistic to define the 95$^{th}$ percentile of high-performing models. Z-scores were calculated using the mean and standard deviation of the subnetwork scores for all subnetworks. Models with a z>1.646 were retained as high-performing models.

A careful reader may note that, though the total number of models does not change, the number of high-performing models is different between cohorts. This is because of small differences in skew. We used a z-score defined 95$^{th}$ percentile because the extrema were more robust to skew.

###### 4.4.3 Summary network Extraction from the Reduced Network

For each reaction in the Reduced Network (**Figure** 3D) we measured the proportion of models in the top-performing model set which used each reaction ($p^*$), the proportion of models in the background model set which used each reaction ($p$), and the enrichment of each reaction in the top-performing model set compared to the background model set ($e(p^*|p)$ (**Figure** S 10).

The Summary network contains the most important paths in the reduced network where importance ($i$) is defined as $i = p^* + p^* e(p^*|p)$. This provides an additional reward for enrichment without inflicting a penalty on essential reactions which are not enriched in the top-performing model set. The important paths are defined as the top 5% of paths from lactose to an observed HMO where a path-score is defined as the sum of reaction importance between lactose and each observed HMO. The reduced network (**Figure** S 9) and Summary network (**Figure** 4) are presented with line thickness corresponding to importance.

##### 4.5 Resolution of ambiguous genes and HMO structure through analysis of candidate models and subnetwork statistics

The top-performing candidate models from all subnetwork models were examined for distinguishing and common features that contributed to their extreme model score. To determine model features associated with improved model performance, we compared model features of the high scoring candidate models to the background of all subnetwork models.

###### 4.5.1 Enrichment of ambiguous HMO structures within top-performing candidate models

Towards disambiguating HMO structures enumerated in **Figure** S 12, we employed two strategies. First, we asked which structures were most enriched. Then, to further clarify ambiguous structures, co-inclusion of ambiguous structures was considered.

We first identified HMO structures that are more prevalent in the top-performing models, compared to all other models. We only consider structures that were included in >1% of all models. Hypergeometric enrichment was computed by comparing the occurrence of each structure in a top-performing set with its occurrence in the background. Those structures with both the highest prevalence in the subnetwork models and significant enrichment over the background were top-performing as the representative structure for that HMO. Those HMOs for which a single clear representative did not arise were examined for co-occurrence with sugars clearly resolved in the initial examination of prevalence and occurrence.

Co-occurrence analyses involved an initial filtration to retain only the top 10% of co-occurring structures. Co-occurrence was measured by the Jaccard index, the ratio of co-inclusions to all inclusions of two HMO structures. The Jaccard index was computed on HMO structure co-occurrence within common models top-performing under cohort 1 and cohort 2. The size of the intersection of retained co-occurrences, the top 10% of Jaccard indexes, were confirmed to be extremely significant (p<<.01) using hypergeometric enrichment. Structures with high centrality in the network of retained co-occurrences were selected from this analysis. Specifically, structures chosen in this analysis showed retained and significant co-occurrence with at least three members of the central clade; the group containing DFLNT1/2 and DFLNH2.

###### 4.5.2 Enrichment of ambiguous enzymes within top-performing candidate models

We identified genes potentially responsible for reactions listed in **Table** 1 as follows. Candidate model scores, $S_{\omega}$, were calculated as the arithmetic mean of maximum gene-linkage scores, $\rho_{gl}$ (S1.1.4). The maximum gene-linkage score will be contributed by the gene expression that best explains the flux predicted by a given model. To expand the model score to a gene-specific interrogation we employ three metrics.

The ***enrichment proportion*** (PROP) describes the increase in the proportion of high-performing models (relative to all models) that prefer a specific gene(**Figure** S 4); the proportion of models where a gene provides the maximum gene-linkage score within a model. It asks if that proportion is significantly higher in the top-performing models ($\Omega^{*}$ top 5% of model score) compared to the background models ($\Omega$). We can ask the number of times a gene ($\gamma$) has the best correlation within a linkage for models in the top-performing ($\Omega^{*}$) and background ($\Omega$) sets:

$$B_{\gamma l} = \{\omega \in \Omega | argmax_{g \in l_g} \; \rho_{gl} = \gamma\} \qquad B^{*}{}_{\gamma l} = \{\omega \in \Omega^{*} | argmax_{g \in l_g} \; \rho_{gl} = \gamma\}$$

Then we can ask, for each gene, if $B^{*}{}_{\gamma l} > B_{\gamma l}$ using a hypergeometric enrichment:

$$HypGeo_{\gamma l} \, (k = |B^{*}{}_{\gamma l}| \, , n = |\Omega| \; , K = |B_{\gamma l}| \, , N = |\Omega| \, )$$

The enrichment measure helps to identify high performing genes by focusing only on the best performing gene in each model.

The ***gene-linkage score metric*** (GLS), described in section S1.1.4, the Spearman correlation between gene expression and normalized flux: $\rho_{gl} = cor(E(g) \sim \underline{f}_{l_r \, \omega \, D_m})$ . This metric provides information on every gene in every model regardless of the relevance of the gene to the model. This helps identify genes with low congruence because every gene is evaluated.

The ***model score contribution*** (MSC) is the Pearson correlation between the model score and the gene-linkage score for a specific gene, $cor(S_{\omega} \sim \rho_{gl})$. This is a more continuous approach to the question posed by the enrichment metric. This is a high granularity metric that can distinguish genes that highly influence the overall score.

### 5 THEORY, CALCULATIONS AND DEFINITIONS

#### 5.1 DEFINING THE MODEL-SPACE: INPUT DATE, REACTION RULES, COMPLETE NETWORK AND REDUCED NETWORK

Our examination relies on two datatypes: glycomic data from High-Performance Liquid Chromatography (HPLC) and microarray data from Illumina Beadchip arrays. These datasets will be referred to as $D = \{D_m, D_g\}$ where $D_m$ refers to the glycomic data an $D_g$ refers to the gene expression data.

Let $M_1$, be a set of monosaccharides: $M_1 = \{Fuc, NeuAc, Glc, Gal, GlcNAc\}$. Let the set of multimeric sugars, $M$, be defined as all biochemically feasible HMOs up to a specified complexity; those which contain a maximum of $k$ monosaccharides. Let $R$, be the set of all feasible reactions, specifically monosaccharide additions, resulting in an HMO in $M$. These reactions will be represented as an ordered set including a reactant (q), product (p), and monosaccharide (a).

$$R = \{r = < q, p, a > \in R \mid \forall_{p \in M} \exists_{q \in M} \exists_{a \in M_1} \; q + a \rightarrow p \}$$

The definition of HMOs (**M**) and reactions (**R**), allows us to define the complete HMO biosynthetic network $\Psi = \{M, R\}$, a directed acyclic graph rooted in a lactose (**Figure** 3B). Let $\Omega$, the Reduced Network (**Figure** 3C-D), be a subgraph of the complete HMO biosynthetic network. We define the Reduced network as a subgraph of the Complete network (**Figure** 3B), $\Omega \subset \Psi = \{M, R\}\; s.t. M \subset M \wedge R \subset R$. The vertices of this network are taken from the set of HMOs, $M$, and the directed edges are taken from the set of reactions, $R$, such that $\Omega = \{M, R\}$. All HMOs which are retained in the reduced network, $\Omega$, $m \in M$, $m$ is an observed HMO ($m \in D_m$), or can be transformed into an observed HMO through a combination of reactions $r \in R$.

$$M = \{ m \in M \mid \forall_{m \in M} m \in D_m \vee \exists_{\langle r_1, \ldots, r_n \rangle \in R^{[0,n]}} \; r_n\big(\ldots r_1(m)\big) \in D_m \}$$

For all reactions retained in the reduced network, $\Omega$, $r \in R$, there exists an HMO which requires at least $r$ to be successfully transformed into an HMO observed by HPLC ($m \in D_m$).

$$R = \{r \in R \mid \forall_{r \in R} \exists_{m \in M} \exists_{\langle s_1, \ldots, s_n \rangle \in R^{[0,n]}} \; s_n\big(\ldots s_1(m)\big) \wedge r \in \langle s_1, \ldots, s_n \rangle\}$$

This reduction of the complete network is done using Flux Variability Analysis.

#### 5.2 IDENTIFYING CANDIDATE MODELS WITHIN THE REDUCED NETWORK

The Mixed Integer Linear Programing generated candidate models (**Figure** 3E), are subgraphs of the reduced network $\omega \in \Omega$. These subgraphs are sufficient to simulate the observed glycomic data with Flux Balance Analysis (FBA). Given an HMO dataset ($D_m$), where each quantity $d_{m_i} \in D_m$ defines an observed HMO (m). We refer to $d_{m_i}$ as the quantification of HMO $m_i$ and we refer to the flux through the reaction leading to $m_i$ as $f_{r_i}$. We define flux as $f_{\omega, D_m} = FBA(\omega, D_m) = \langle f_{r_1}, \ldots f_{r_n} \rangle_{\omega, D_m}$ (**Figure** 3F). To obtain the flux for a given reaction, $r_i$, we define a function $h\colon r_i \rightarrow f_{r_i}$. For a candidate model to be valid the flux distribution must match the data and be unique. Uniqueness means that no other flux distribution in the given model would reproduce the observed data. Matching the observed data means that the sum of flux through HMO-specific sink reactions (exiting/observed flux) is equivalent to the values observed in $m \in D_m$.

$$MILP(\Omega) = \{\omega = \{M', R'\} \subset \Omega \mid \forall_{d_{m_i} \in D} \exists_{f_{m_i} \in f_{\omega,D}} d_{m_i} = f_{m_i} \wedge \forall_{f_{m_i} \in f_{\omega,D}} f_{m_i} = f_{m_i} \}$$

#### 5.3 Defining the Candidate Model Score

The Model Score (S) is defined by the best correlations between simulated flux and gene expression which may be supporting that flux. We first group similar reactions into 10 groups (**Table**1) referred to as the linkages (L). The linkages are groups of reaction, $r \in R$, which involve equivalent monosaccharide additions of the same "type;" same monosaccharide added to the same terminal sugar via the same $\frac{\alpha}{\beta}$ configuration on the same carbons. Reactions of the same types are assumed to be supported by the same gene expressions. Therefore, each linkage $l \in L$ maps to both reactions of the same type and genes which may support those reactions: $l = \{ l_r, l_g \}$. Every reaction-linkage set $l_r \in l \in L$ is a set which contains several equivalent reactions $\forall_{l \in L} \forall_{r,s \in l} type(r) = type(s)$. All the linkage sets are disjoint as no reaction contains the addition of multiple monosaccharides,

$$\cup_{l_r \in l \in L}\, l_r = L \text{ but } \cap_{l_r \in l \in L}\, l_r = \emptyset$$

Each flux in this calculation is normalized to a proportion of the flux of its parent reaction $\underline{f} = \frac{f}{f'}$ where $f'$ is the reaction which feeds $f$. Flux through a linkage is calculated as the sum of flux through all reactions within a linkage. Recall, every flux distribution, $f_{\omega,D_m} = FBA(\omega, D_m) = \langle f_{r_1}, \dots f_{r_n} \rangle_{\omega,D_m}$, is defined by the structure specified by a candidate model ($\omega$) and parameterized by the glycomic data ($D_m$). Therefore, we can specify the linkage-flux $\underline{f}_{l_r \in l \in L}$ as:

$$\underline{f}_{l_r\,\omega\,D_m} = \sum_{r \in l_r} \frac{h(r)}{h(r)'} = \sum_{r \in l_r} \frac{f_r \ \in f_{\omega D_m}}{f'_r \ \in f_{\omega D_m}} = \sum_{r \in l_r} \underline{f_r}_{\ \omega\, D_m}$$

Every gene-linkage set $l_g \in l \in L$ contains genes which are suspected to support the reactions in $l_r \in l$. The gene linkage sets are not disjoint because several linkages involve the same monosaccharide and further specification about the type of reaction the associated genes are expected to perform is not yet available. Therefore,

$$\cup_{l_g \in l \in L}\, l_g = L \text{ but } \cap_{l_g \in l \in L}\, l_g \neq \emptyset$$

The model score (S) for each candidate model $\omega$ can be defined as the average of the maximum normalized correlation between predicted flux through a given reaction and the gene expression predicted to support it. The normalization is a z-score normalization where all values are converted to z-scores using the mean ($\mu$) and standard deviation ($\sigma$) of a common group. In this case, the z-score mean and standard deviation is computed from the correlations of all genes-flux correlations within a linkage. The function z will denote this normalization from a set of correlations $x$ to a normalized set of values relative to other correlations within that linkage $z_l : x \rightarrow N(x, \mu_l, \sigma_l)$. We will also refer to the expression of a gene (g) as $E(g)$.

$$S_\omega = \left\{ z_l \left( cor \left( E(g) \sim \underline{f}_{l_r\,\omega\,D_m} \right) \right) \right\}$$

#### 5.4 Flux Variability Analysis (FVA)

The Flux Variability analysis is related to a min-max Linear Programming (LP) problem (i.e. each reaction of the network is maximized and subsequently minimized). It can be written as follows[20,106]

$$\forall v_i, i = 1, \ldots, N \qquad v_{i,upper} = (v_i) \qquad v_{i,lower} = (v_i) \quad s.t. \quad Sv = 0$$

where $v$ is the vector of specific reaction rates (i.e. metabolic fluxes), $N$ is the number of fluxes in $v$ , $v_{i,upper}$ and $v_{i,lower}$ are respectively the upper and lower values of each flux $v_i$ satisfying the system of linear equations. The LP problem formulated as above is solved using the function *fastFVA* in the CobraToolbox 2.0.

#### 5.5 Mixed-Integer Linear Programming (MILP)

The enumeration of all alternate minimal reaction sets solving equally $Sv = 0$ can be formulated with the following MILP problem[108,109]:

$$Min\ Z = \sum_{i=1}^{N} w_i$$

Subject to

$$Sv = 0$$

$$\sum_{i \in NZ^{J-1}} y_i \ \geq 1 \qquad with\ y_i \in \{0,1\}$$

$$\sum_{i \in NZ^{J}} w_i \ \leq \left|NZ^k\right| - 1 \qquad with\ w_i \in \{0,1\}\ and\ k = 1{:}J - 1$$

$$w_i + y_i \leq 1 \qquad \forall i$$

$$w_i.lb \leq v_i \leq \ w_i.ub \qquad \forall i$$

Where $S$ is the stoichiometric matrix, $v$ is the vector of specific reaction rates (i.e. metabolic fluxes), $N$ is the number of fluxes in $v$ , $y$ and $w$ are vector of binary variables and $lb$ and $ub$ are respectively the lower and upper bounds for the individual flux values. The MILP problem formulated as above is solved using the function *solveCobraMILP* in the CobraToolbox 2.0.

### 6 SUPPLEMENTARY RESULTS

#### 6.1 CANDIDATE GENE FILTERING BY EXPRESSION AND ACCEPTOR SPECIFICITY

We initialized our candidate gene lists naively, including all members of a gene family corresponding to each linkage reaction (**Table** 1). Here, we query existing literature and expression data to narrow the initial list of gene family members to those relevant to HMO biosynthesis (**Table** S 1). Overall, a candidate gene was excluded from further consideration if it was not measured in the microarrays and that non-expression was confirmed by an independent RNA-Seq or if the gene was documented to perform an irrelevant reaction. If a gene is unmeasured in microarray and measured in RNA-Seq, it cannot be evaluated or ruled out. We further validated the RNA-Seq data by ranking it relative to expression in normal human tissues from GTEx[32]. If literature and expression data conflict (e.g. a gene performing a relevant reaction is not expressed), conflicts are resolved by the independent RNA-seq data; a relevant gene is not part of our system if it is not expressed.

We first considered the likelihood a gene may perform the corresponding reaction considering the existing literature[119] and databases such as glycogene database[120], BRENDA[121], Uniprot[122], MetaCyc[123], KEGG[124]. Because the characterization of these genes with HMO acceptors is historically limited, we gave special attention to recent work in HMO chemosynthesis[52]. If a gene was used by Prudden et. al. to perform a specific reaction, we know it can perform the reaction. Similarly, if the linkage reaction has been demonstrated previously in HMO, lactoceramide derived, or a similar (GalNAc or Glc instead of GlcNAc) context, we retain that gene for further consideration. Genes were excluded from further examination if characterized to use a different donor including B4GAT1, formerly B3GNT1[125,126], and B3GALNT1, formerly B3GALT3[69,80]). Similarly genes were excluded from further examination if they were known to synthesize a different linkage (e.g. FUT8 and the α-1,6 Fucose addition[127,128]) or use an unrelated acceptor (e.g. glycosaminoglycan (GAG) glycosyltransferases like B3GNT7[129–131], B3GALT6[132,133] and B4GALT7[134,135]. Through review of existing literature, we were able to rule out six glycosyltransferases reducing the candidate genes from 54 to 48.

We further examined microarray gene expression data from cohort 1 (GSE36936) and cohort 2 (GSE12669) to determine which of the candidate genes were expressed; we summarized expression to the third quartile (Q3) or 75th percentile to account for infrequent expression. The microarrays failed to measure the expression of 25 glycogenes; 2 genes lacked probes, 11 genes showed no expression (Q3=0) in one microarray, 15 genes showed no expression (Q3=0) in both microarrays. Negligible expression was confirmed in an independent RNA-Seq study of the milk fat globules in lactating women (GSE45669, [31]) and comparison to global expression distributions in GTEx[32]; genes unmeasured in the microarrays that did not meet the expression criteria--Q3 TPM>2 or (Q3 TPM<=2 and Q3 TPM > 50% of GTEx samples)--were excluded from further consideration. Of the 25 unmeasured glycogenes, 17 were confirmed unmeasured in the Lemay RNA-Seq relative to GTEx and excluded from further consideration while 8 genes could not be confirmed as non-expressed. 2 of the 8 unconfirmed non-expressed genes are known to perform irrelevant reactions--FUT8 and B3GNT7; the remaining 6 genes--B3GNT9, FUT5, FUT9, FUT10,

B3GALT5, and ST6GALNAC3--cannot be ruled out or evaluated in this study due to a lack of measurement. Unfortunately, 3 of the 6 questionably unmeasured genes--FUT5, FUT9 and B3GALT5--were demonstrated to synthesize these reactions in HMO[52]. Because low B3GALT5 expression was marginal and isolated to one microarray, we chose to include it in later analysis. 4 genes used in the Prudden experiment were successfully excluded due to low and RNA-Seq confirmed non-expression--FUT1, ST3GAL4, GCNT2B, and ST6GALNAC5. Non-expression of GCNT2B was confirmed with exon-level re-analysis of the Lemay RNA-seq data (STARv2.5.4b alignment to GRCh38); nearly all of the reads aligning to the three exons (1a, 1b, and 1c) definitive of isoforms A, B and C, aligned to exon 1a. Ultimately, 25 genes were retained for further analysis, 22 genes were excluded due to confirmed low expression or catalytic irrelevance, and 6 were designated removed from further analysis without being ruled out.

#### 6.2 GENERATION OF 44,984,988 CANDIDATE MODELS REPRESENTING HMO BIOSYNTHESIS

To explore how human mammary gland epithelial cells synthesize HMOs, we generated candidate models based on a complete network describing all possible reactions leading to the synthesis of all possible HMO structures. HMO complexity was limited to 7 monosaccharides added to the starting lactose. The complete network was pruned using flux variability analysis, and reactions are removed if they are not used to synthesize the 16 most abundant HMOs. Using mixed integer linear programming, we enumerated all subnetworks that can accurately simulate the synthesis of observed oligosaccharides. The MILP produced 48,914,738 subnetworks. 3,929,750 of these subnetworks were unable to uniquely (upper and lower bound flux were not equal) simulate the HPLC data leaving 44,984,988 candidate models for further examination. Each candidate model retained 43-54 reactions, 0.4-.5% of the reactions found in the complete model and 19.5-24.4% of the reactions retained in the Reduced Network (**Table** S 4). These models covered all the possible combinations of HMO synthesis by the 10 known glycosyltransferase families that could describe the synthesis of the HMOs in this study.

#### 6.3 SELECTION OF BEST PERFORMING MODELS

We computed the model score $S_\omega$ using the glycoprofiling and transcriptomic data from two independent cohorts. When sorted, these scores reveal a sigmoidal trend (**Figure** S 20A, **Figure** S 20F) suggesting an enrichment of high and low scoring models. Scores falling into the top 5% of density of the normal distribution of scores were ***top-performing*** ($\Omega^*$, red) to represent high performing models (Described and justified in *Ranking subnetwork performance and selection of candidate models* in Methods). We found 2.66 million top-performing models using cohort 1, and 2.32 million top-performing models using cohort 2 (**Table** S 4). 241,589 models were common to the top-performing in cohort1 and cohort 2. Fewer top-performing models were expected from dataset 2 because it is a smaller dataset, and therefore its predictive power is limited; this does not present an issue as the primary purpose of dataset 2 is validation. As an initial validation, we mapped top-performing models from dataset2 (red) onto the sigmoid for dataset 1 (**Figure** S 20B) and did the same for top-performing models from dataset 1 (red, **Figure** S 20G). We note that high performing models in both datasets appear at the top of the sigmoid for the other dataset suggesting the selection is consistent. We also note that the intersection of models top-performing in analysis parameterized by dataset 1 and dataset 2 also appear at the top of both sigmoids (red, **Figure** S 20C, **Figure** S 20H).

Normality of the score distributions was also examined using Q-Q plots and simple distribution visualization. The candidate model scores for dataset 1, $S_{\omega}(D_1)$, are normally distributed as evidenced by the nearly linear Q-Q plot (**Figure** S 20D). The candidate model scores for dataset 2, $S_{\omega}(D_2)$, is less normally distributed as evidenced by the moderate deviation in the Q-Q plot (**Figure** S 20I). The deviation in the Q-Q plot for dataset 2 scores is expected as it is a smaller dataset therefore there is a moderate skew to lower scores due to limited power. Since dataset 2 is not being used for discovery, but rather validation, this moderate skew is manageable.

### 7 Supplemental Discussion

#### 7.1 Validity of Selected Transcription Factors

To further validate the selected genes, we analyzed the promoters and gene expression patterns for common regulatory elements; if these genes are co-regulated, during lactation, they should enrich for common transcription regulatory elements. Two studies implicated IKZF1 in regulating α-1,2 fucosylation through high-throughput analyses including type-1-diabetes GWAS [136] and text-mining TF prediction [137]. SP1, a TF already implicated in prolactin reception[138], has been known to transcriptionally regulate gene expression in several β4-galactosyltransferases including B4GALT4 [139] and B4GALT5 [140–142]. Consistent with the miRNA proxy theory [143], SP1 can also regulate α-1,3-fucosyltransferase, FUT4, for Lewis Antigen X biosynthesis [144] through miR-29b. The ETS gene superfamily encodes TFs (e.g., ETS1, ETV4 and ERG) that bind a purine-rich DNA sequence through the ETS domain [145], and ST3GAL1 contains several TFBS in its promoter region, including ETS1 [146]. Together, our results of the discovered TFs and TFBS motifs are consistent with those reported for regulating GTs in similar biosynthesis reactions.

#### 7.2 Leveraging HMO pathway resolution to clarify natural variation in HMO composition

With the newly reduced space of HMO biosynthetic pathways and knowledge of the enzymes and their regulation will enable mechanistic insights into the relationship of maternal genotype and infant development. HMO concentrations vary dramatically between mothers creating sometimes unpredic**Table** health risks for newborns. Numerous factors influence HMO composition. These include genetic factors influencing blood type, secretor and Lewis status[23–26,47,60], diet[7], geography[7,48]. Variations in HMO composition influence infant susceptibility to infection[37,39,40,147], impact risk of developing diseases such as necrotizing enterocolitis[38] and influence development[1,2,5,6] and contribute to childhood obesity[41,42]. To elucidate the mechanisms leading to the heterogeneity among women, and its connection to child health, the differences in HMO composition often need to be interpreted in the context of all the biosynthetic pathways.

#### 7.3 Leveraging HMO pathway resolution to facilitate chemoezymatic synthesis

Finally, once essential HMOs are identified, the knowledge presented here on the HMO biosynthetic network can provide insights for the industrial production of HMO as a nutraceutical. Currently, commercially available formula is fortified with galactooligosaccharide (GOS), fructooligosaccharide (FOS) and 2'-fucosyllactose (2'-FL). These can be probiotic[44], modulate immune development[45], and reduce pathogenic susceptibility[46]. As larger health-relevant HMOs emerge[38], there is a need to scale up their production. Chemoenzymatic synthesis is feasible[49–53], but cell-based production can be more easily scaled. The growth of microbial HMO production is limited by knowledge of the biosynthetic system[54,55]. Small HMOs like 2’FL[56–58] and larger molecules like LNT have been produced in *E. coli*[59]. Increased knowledge of enzymes and reactions collected in this work will expand the toolbox for metabolic engineering by narrowing and prioritizing the list of relevant glycosyltransferases to HMO biosynthesis.
